# Comprehensive characterisation of cell-free tumour DNA in plasma and urine of patients with renal tumours

**DOI:** 10.1101/758003

**Authors:** Christopher G Smith, Tina Moser, Johanna Burge, Matthew Eldridge, Anja L Riediger, Florent Mouliere, Dineika Chandrananda, Katrin Heider, Jonathan CM Wan, Anne Y Warren, James Morris, Irena Hudecova, Wendy N Cooper, Thomas J Mitchell, Davina Gale, Andrea Ruiz-Valdepenas, Tobias Klatte, Stephan Ursprung, Evis Sala, Antony CP Riddick, Tevita F Aho, James N Armitage, Samantha Perakis, Martin Pichler, Maximilian Seles, Gabriel Wcislo, Sarah J Welsh, Athena Matakidou, Tim Eisen, Charles E Massie, Nitzan Rosenfeld, Ellen Heitzer, Grant D Stewart

**Affiliations:** Cancer Research UK Cambridge Institute, Li Ka Shing Centre, Robinson Way, Cambridge CB2 0RE, UK; Cancer Research UK Major Centre – Cambridge, Cancer Research UK Cambridge Institute, Li Ka Shing Centre, Robinson Way, Cambridge CB2 0RE, UK; Medical University of Graz, Diagnostic and Research Center for Molecular Biomedicine, Institute of Human Genetics, Austria; Cambridge University Hospitals NHS Foundation Trust, Cambridge CB2 0QQ, UK; Amsterdam UMC, Vrije Universiteit Amsterdam, department of Pathology, Cancer Center Amsterdam, De Boelelaan 1117, 1081 HV, Amsterdam, The Netherlands; Wellcome Sanger Institute, Hinxton CB10 1SA, UK; Medical University of Graz, Department of Internal Medicine Graz, Austria Division of Oncology, Graz, Austria; Medical University of Graz, Department of Urology, Graz, Austria; Military Institute of Medicine, Department of Oncology, Warsaw, Poland; AstraZeneca, 1 Francis Crick Avenue, Cambridge Biomedical Campus, Cambridge CB2 0AA, UK; Department of Oncology, University of Cambridge, Cambridge, CB2 0QQ, UK; Hutchison/MRC Research Centre, University of Cambridge, Cambridge, CB2 0QQ, UK; Department of Surgery, University of Cambridge, Cambridge CB2 0QQ, UK; Department of Urology, Royal Bournemouth Hospital, Bournemouth, UK; Department of Radiology, University of Cambridge, Cambridge, CB2 0QQ, UK

## Abstract

Cell-free tumour-derived DNA (ctDNA) allows non-invasive monitoring of cancers but its utility in renal cell cancer (RCC) has not been established. Here, untargeted and targeted sequencing methods, applied to two independent cohorts of renal tumour patients (n=90), were used to determine ctDNA content in plasma and urine. Our data revealed lower plasma ctDNA levels in RCC relative to other cancers, with untargeted detection of ∼33%. A sensitive personalised approach, applied to plasma and urine from select patients improved detection to ∼50%, including in patients with early-stage and even benign lesions.

A machine-learning based model predicted detection, potentially offering a means of triaging samples for personalised analysis. In addition, with limited data we observed that plasma, and for the first time, urine ctDNA may better represent tumour heterogeneity than tissue biopsy. Furthermore, longitudinal sampling of >200 plasma samples revealed that ctDNA can track disease course. Additional datasets will be required to validate these findings.

Overall, our data highlight RCC as a ctDNA-low malignancy, but indicate potential clinical utility provided improvement in detection approaches.

**One sentence summary:** Complementary sequencing methods show that cell-free tumour DNA levels are low in renal cancer though, via various strategies, may still be informative.

## Introduction

Renal cell carcinoma (RCC) is the most lethal urological malignancy with 50% of patients that develop the disease dying from it^[1]^. Clinical management challenges include: early diagnosis; differentiation of histological subtypes, i.e. chromophobe RCC (chRCC) from clear cell RCC (ccRCC) or benign oncocytoma; identification of patients with minimal residual disease following intended curative nephrectomy which will allow improved stratification of patients for adjuvant therapy trials; and predicting and tracking of response to targeted therapies. RCC has well-established pathological and genetic heterogeneity^[2]^, which confounds development of personalised medicine^[3]^. Moreover, ccRCC exhibits a broad range of metastatic phenotypes^[4]^ highlighting the need for longitudinal sampling. Due to the invasive nature of the procedure and failure to capture genetic heterogeneity, tissue biopsies potentially inadequately inform treatment decisions^[5]^. A ‘liquid biopsy’, providing an admixture of the entire tumour burden of a patient, may offer a non-invasive alternative to traditional tumour sampling techniques. Cell-free DNA (cfDNA), which in patients with cancer contains cell-free tumour derived DNA (ctDNA), represent one such promising liquid biopsy strategy^[6–8]^.

Despite showing great promise in various cancers^[6]^, there is little and often contradictory data of ctDNA as a tool in RCC in locally advanced and metastatic RCC^[6, 9–11]^. As such, there remains an unmet need for the characterisation of the levels, and potential clinical utility, of ctDNA in renal cancers of differing stage and subtype. Furthermore, whilst evidence suggests ctDNA in the urine can be informative in urological cancers^[12, 13]^, no previous study has assessed the presence of ctDNA in the urine of RCC patients. Here we aimed to determine the presence, levels, heterogeneity and potential clinical applications of ctDNA in plasma and urine of 90 patients with renal tumours ranging from benign oncocytomas through to metastatic RCCs using both untargeted genome-wide, and targeted sequencing approaches.

## Results

### Untargeted analysis of ctDNA in plasma and urine from patients with renal tumours

We applied a combination of rapid and cost-effective untargeted approaches, albeit with limited sensitivity, to establish a first measure of ctDNA presence and levels in patients with benign through to metastatic disease (n=90 patients from the DIAMOND and MonReC studies **(Fig. 1A-C and table S1-S2)**. First, we assessed overall ctDNA levels using the trimmed Median Absolute Deviation (tMAD) score calculated from shallow whole genome sequencing (sWGS)^[14]^ of plasma from 47 DIAMOND patients **(Fig. 1B)**. As this method is dependent on the presence of somatic copy number alterations (SCNA), we applied sWGS/tMAD to matched tumour tissue, available from 28 of 47 patients (59.6%). All had SCNA **(Fig. S1)**, suggesting that SCNA are a valid ctDNA target in patients. However, in plasma we detected SCNA in only 3 of 47 (6.4%) patient samples (**Fig. 2A**, one patient with metastatic ccRCC and 2 with non-metastatic chRCCs).

**Fig. 1.**
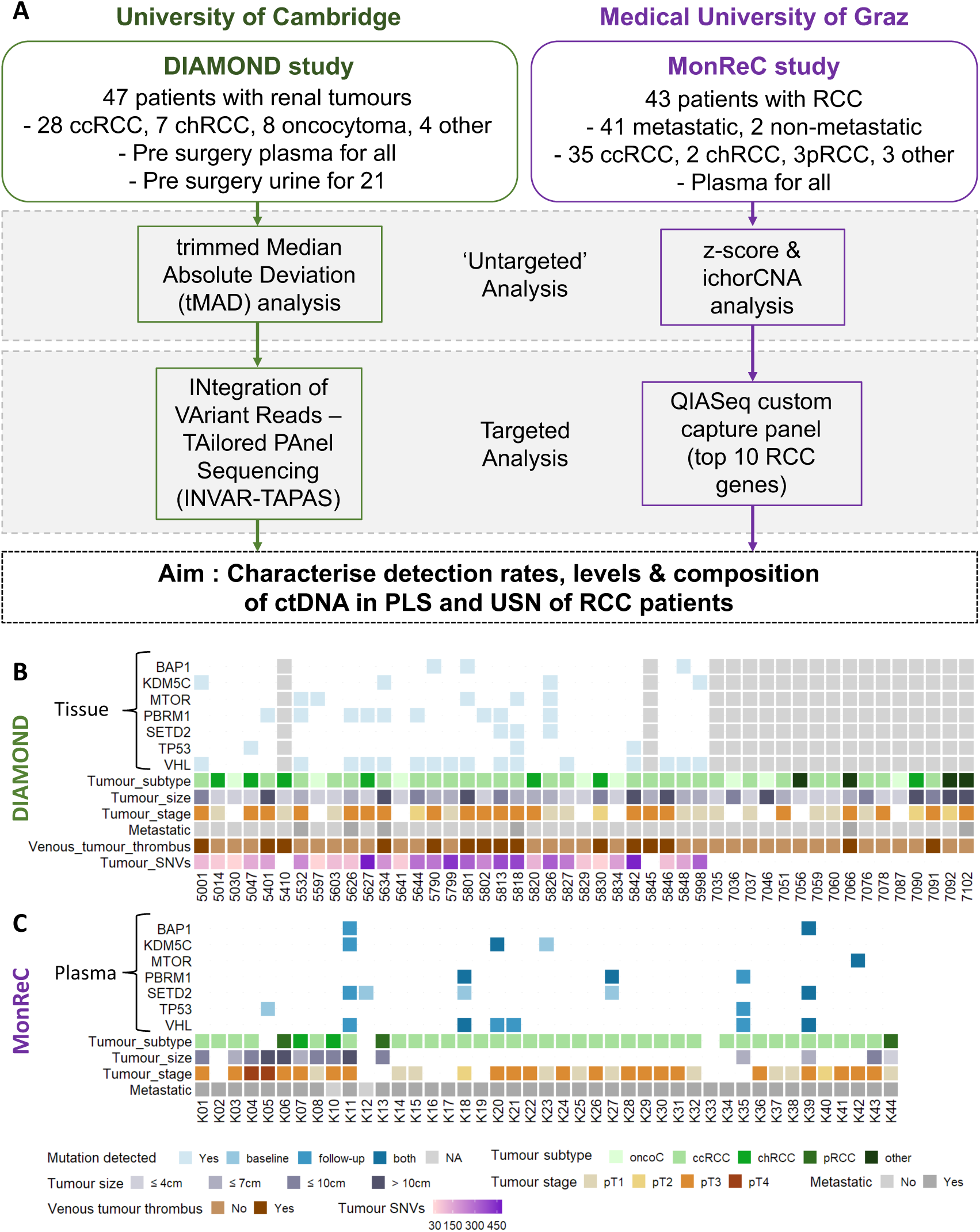
Study design, patient characteristics and tumour genomic profile. **(A)** ctDNA analysis in RCC patients was applied to two patient cohorts, DIAMOND and MonReC. Initially, untargeted sequencing methods were applied to samples; For DIAMOND, tMAD analysis of sWGS data was applied. For MonReC, a combination of z-score analyses of mFastSeq data and ichorCNA analysis of sWGS data. Subsequently, targeted sequencing methods were used; For DIAMOND, INVAR-TAPAS was applied to patient plasma (n=29) and urine (n=20). For MonReC, a QIASeq custom capture panel targeting the 10 most commonly mutated genes in RCC patients was applied. **(B)** For DIAMOND, plasma (n=47) and urine (n=21) were collected from patients with a range of tumour subtypes and stages. Specifically, 28 ccRCCs (11/1/16 stage I, II & III respectively), 7 chRCCs (2/2/3 2 stage I, II & III), 8 oncocytomas, 1 patient with papillary RCC (stage III), 1 patient with a MiT family translocation RCC (stage II) and 2 patients with oncocytic renal neoplasm. Shown, in descending order, are tumour tissue mutation status of frequently mutated RCC genes (pale blue cubes indicate that a mutation was detected, white space indicates that no mutation was detected, grey columns indicate that tissue was not available for that patient), tumour -subtype, -size, -stage, metastatic at baseline, evidence of venous tumour thrombus, and number of tumour SNVs (targeted for INVAR-TAPAS) **(C)** For MonReC, plasma (n=43) was collected from 41 patients with metastatic RCC and two with localized RCC. Shown, in descending order, are plasma mutation status (after QIASeq, blue, medium blue and dark blue cubes indicate that a mutation was detected at baseline, during follow-up, or at both time points respectively) of frequently mutated RCC genes, tumour -subtype, -size and metastatic at time of sampling. More comprehensive versions of **(B)** and **(C)** provided in **Fig. S13**.

**Fig. 2.**
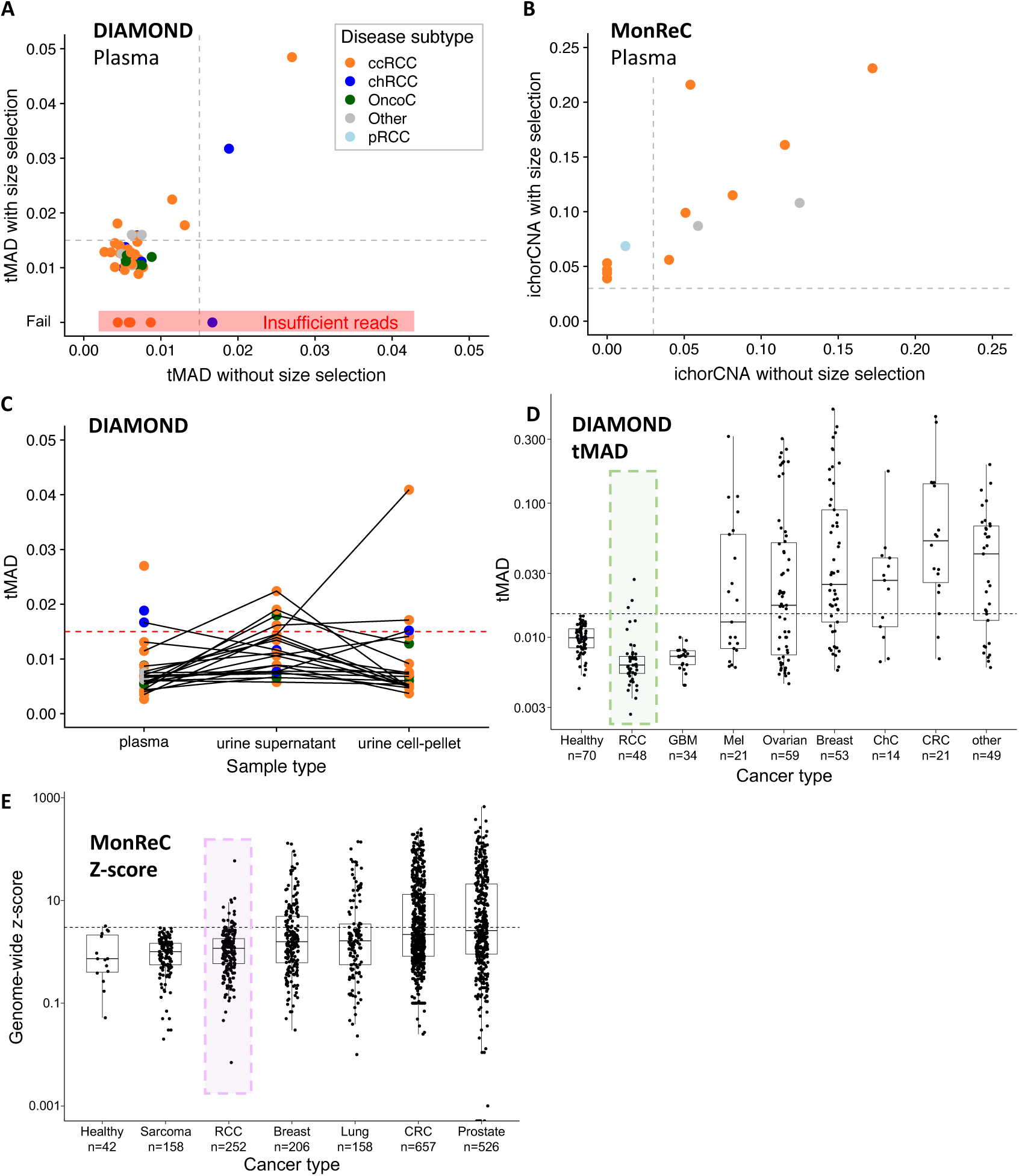
ctDNA detection using untargeted assays. **(A)** Distribution of tMAD scores across DIAMOND plasma samples (x-axis). Data points are coloured according to disease subtype. A tMAD score of >0.015 (grey dashed line) indicates SCNA, and thus ctDNA. ctDNA was detected in 3/47 (6.4%) plasma samples. Data on the y-axis show tMAD scores for the same plasma samples after *in silico* size selection for sequencing reads 90-150bp in length. On average the tMAD score increased 2.2 fold (range 1.25-4.83) and led to ctDNA detection in 8 additional patient samples, resulting in ctDNA detection in 11/47 (23.4%) DIAMOND patients. Four patient samples had insufficient sequencing reads after size selection for tMAD analysis (red highlight). **(B)** Tumour fraction of MonReC ctDNA positive plasma samples (n=14), as calculated by ichorCNA, before and after *in silico* size selection for sequencing reads 90-150bp in length. On average tumour fraction increased 2.2-fold (range 0.9 – 5.7) and revealed six patients with detected ctDNA, in addition to the eight patient samples detected without size-selection. **(C)** Plot showing distribution of tMAD scores across DIAMOND plasma, urine supernatant (USN) and urine cell pellet (UCP) samples. Samples from the same patient are connected by grey lines. The detection threshold is indicated by a red dashed line. **(D)** tMAD and **(E)** z-scores score distribution of RCC samples were compared to samples from other cancer types collected at the University of Cambridge^[14]^ and Medical University of Graz respectively. Renal samples are highlighted. GBM = Glioblastoma, Mel = Melanoma, ChC = Cholangiocarcinoma, CRC = Colorectal cancer. A similar comparison was carried out using the ichorCNA metric (**Fig. S5C**).

Next, we employed *in-silico* selection of sequence reads within particular DNA fragment size ranges, an approach demonstrated to enrich for mutant signal in plasma^[14, 15]^. After selection of reads between 90-150bp, 41/47 plasma samples from DIAMOND met the criteria for evaluation by tMAD analysis (>2 million reads) **(Fig. S2A)**. On average, tMAD scores increased 2.2-fold (range 1.25-4.83) and led to ctDNA detection in 8 additional patients **(Fig. 2A**) including a patient with an oncocytoma (**Fig. S3A**). Thus, in total we detected plasma ctDNA in 11/47 (23.4%) DIAMOND patients.

The MonReC cohort consisted primarily of metastatic patients (two non-metastatic), most with their primary tumour removed (35/43 patients) (**Fig. 1C**). Samples were initially analysed with untargeted methods, mFAST-SeqS^[16]^ and sWGS (ichorCNA tumour fraction). A mFAST-SeqS z-score of ≥3 indicates tumour fractions of >3-10% (depending on the number and amplitude of SCNA)^[17]^. At baseline, z-scores ranged from −0.6-3.7 (median=0.7) with only 2/43 patients (4.7%) surpassing the detection threshold. The ichorCNA algorithm^[18]^ revealed six further patients with detected ctDNA (8/43=18.6%; **Fig. 2B**) with tumour fractions up to 0.17 (median 0.07, range 0.04-0.17). As above, *in-silico* size selection further improved the ichorCNA detection rate to 14/43 (32%) **(Fig. 2B, Fig. S4 and 5)**. On average, tumour fraction from ctDNA positive samples increased 2.2-fold (range 0.9-5.7) to a median of 0.08 (range 0.04-0.23).

As sampling of biofluids collected in close proximity to the tumour site may improve detection^[13, 19]^, we analysed urine supernatant (USN) samples from 21 DIAMOND cohort patients. Applying tMAD to urine for the first time **(Fig. S2B)**, ctDNA was detected in 4 patients (19.0%) **(Fig. 2C)**, including three patients with ccRCC and a patient with oncocytoma (not detected in plasma, **Fig. S3B**). In addition to USN, we had access to urine cell pellet (UCP) DNA for these 21 patients. UCP DNA is not cell-free but allows non-invasive detection of tumour DNA^[13, 20]^. UCP tMAD analysis revealed 3/20 (15%) patients with detected ctDNA, including one with localised ccRCC, the largest tumour of the cohort with a diameter of 23cm. Comparison of plasma, USN and UCP data did not reveal a clear relationship in detection amongst these compartments **(Fig. 2C)**, confirming previous observations in bladder cancer^[13, 21]^. Considering only those patients for which we had access to plasma and urine, the detection rate increased to 9/21 (42.9%). Of note, only one patient (5842, **table S1**) had detected ctDNA in both plasma and urine.

Overall, untargeted sequencing methods employed here suggested low ctDNA detection rates in plasma and/or urine of patients with renal tumours (16/47, 34% in DIAMOND, 14/43, 33% in MonReC). Even in metastatic disease, the detection rate was only 34% (MonReC 13/41, DIAMOND 3/6). Comparison of plasma ctDNA levels, as quantified by tMAD, mFAST-SeqS and ichorCNA, against other cancer types confirmed that ctDNA levels are lower in renal tumours **(Fig. 2D&E, Fig. S5C)**. Parallel analysis of urine (USN and UCP) also revealed low detection rates. ctDNA was detected in either fluid in patients with a range of tumour subtypes including, unexpectedly, patients with benign oncocytoma. Given these low detection rates, we hypothesised that techniques with greater sensitivity were required to quantify ctDNA in renal tumours.

### Targeted analysis of ctDNA yields improved detection rates

For DIAMOND patients, INtegration of VAriant Reads – TAilored PAnel Sequencing (INVAR-TAPAS) was used, an approach demonstrated to detect plasma ctDNA to parts per million^[22]^ (**Fig. S6**). This method relies on *a priori* knowledge of tumour specific mutations and thus, for 29 DIAMOND patients, we carried out whole exome sequencing (WES) of matched tumour tissue and buffy coat (**Fig. S7**). We observed extensive disease heterogeneity as previously described^[2, 4]^ (**Fig. S8**). Patient-specific mutations of key RCC genes are listed in **table S4**. A personalised capture panel was designed targeting all patient specific SNV from tissue WES as well as the coding regions of 109 genes commonly mutated in renal tumours (**table S5**). Based on the requirements of the INVAR algorithm, seven plasma samples (7/29, 24.1%) had insufficient reads, resulting in limited ability to detect ctDNA, and were thus excluded as technical failures (**Fig. S7A & 9A**).

In the 22 remaining plasma samples, ctDNA was detected in 12 (54.5%) (**Fig. 3A**). Ten of these (83.3%) were ccRCC samples, including three detected by tMAD analysis of sWGS data. It is noteworthy that 9 of these patients had the largest evaluable ccRCC tumours in the cohort (7.4cm - 23cm). However, ctDNA was also detected in a patient with a small (2.8cm) ccRCC with global mutant allele fraction, gmAF=6.4×10^−5^. Of the remaining patients with detected ctDNA, one had a small chRCC (2cm) with gmAF 1.8×10^−4^, and the other had a benign oncocytoma (3.9cm) with gmAF of 2.7×10^−4^. Assessment of all tumour subtypes revealed a significant (p=0.033, Mann Whitney’s U test) correlation between ctDNA detection and tumour size (**Fig. 3E, Fig. S10A**). Similarly, ctDNA detection was more likely in patients with locally advanced RCC denoted by renal vein or inferior vena cava tumour thrombus (p<0.05 for detection by INVAR-TAPAS +/− tMAD, Fisher’s exact test; **Fig. 3F, Fig. S11A-B**). Conversely, Ki-67 assessed cellular proliferation rate did not correlate with detection (**Fig. S11C-E**).

**Fig. 3.**
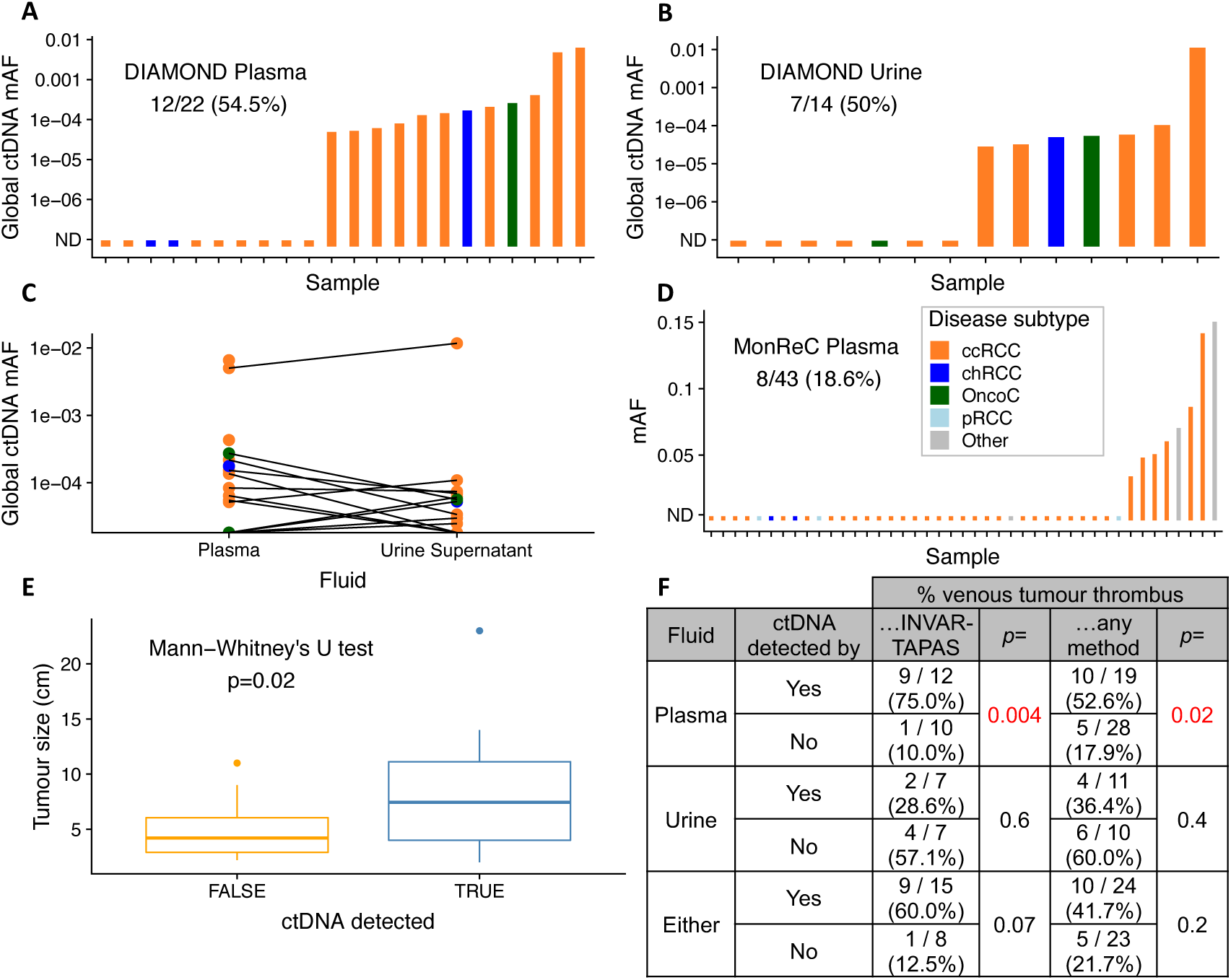
ctDNA detection using targeted assays. **(A)** Application of INVAR-TAPAS to DIAMOND plasma samples. ctDNA was detected in plasma of 12/22 (54.5%), with global ctDNA mAF (gMAF) shown on the y-axis (**Fig. 4**). Disease subtype is indicated by bar colour (legend in Fig. 3D). **(B)** The same analysis was applied to DIAMOND USN samples. ctDNA was detected in 7/14 (50%) patients. **(C)** Comparison of gmAF of plasma and USN samples. In patients for whom we had access to both fluids, lines connect data points (Spearman’s rho = 0.28, p=0.3). **(D)** Summary of targeted sequence analysis using a 10-gene QIASeq panel. Mutations at baseline were detected in 8/43 (18.6%) MonReC plasma samples. The y-axis denotes mAF which ranged from 3.5×10-2-0.15 (if two or more mutations detected, the mean was calculated). **(E)** Assessment of the correlation between primary tumour size (diameter, cm), and ctDNA detection. Detection was via tMAD and/or INVAR-TAPAS, and in either fluid. This observation was driven by plasma (**Fig. S6A**) with no apparent relationship in urine (**Fig. S6B**). **(F)** ctDNA detection in plasma was significantly more frequent amongst patients with venous tumour thrombus as compared to those without. This was not the case when considering ctDNA in urine or ctDNA in either fluid (**Fig. S11**).

For the first time we applied INVAR-TAPAS to USN from 20 patients. As in plasma, six samples (30%) had insufficient sequence reads and were excluded as technical fails. We detected ctDNA in USN of 7/14 patient samples (50%) (**Fig. 3B**; ccRCC n=5, chRCC n=1, oncocytoma n=1). Two of these patients had detected urine ctDNA by tMAD analysis, whilst four had detected ctDNA in plasma (by INVAR-TAPAS or tMAD). Of note, the oncocytoma patient (histology confirmed; **Fig. S12**) had ctDNA detected in plasma (gmAF USN 5.7×10^−5^ vs plasma 2.7×10^−4^). In contrast to plasma, there was no correlation between USN ctDNA detection and lesion size across all patients (**Fig. S10B**), venous tumour thrombus invasion (**Fig. 1B, Fig. 3F, Fig. S11A-B**) or proliferation rate (**Fig. S11C**). There was no correlation between the global ctDNA mAF in plasma and urine (Spearman’s rho = 0.28, p=0.3; **Fig. 3C**), though in the majority of patients, levels proved too low for accurate quantification of tumour fraction.

For metastatic RCC patients recruited to MonReC, no tumour tissue was available and so a *de novo* mutation calling approach was applied to plasma DNA. A gene panel targeting ten significantly mutated genes in renal cancers (*BAP1, KDM5C, MET, MTOR, PBRM1, PIK3CA, PTEN, SETD2, TP53, VHL)*^[23]^ was used with a maximal achievable sensitivity of 5×10^−3^ mAF. Based on existing data, one would expect a somatic mutation to be present in >80% of metastatic ccRCC patients in at least one of these gene^[24]^. However, ctDNA was detected in only 8/43 (18.6%) baseline samples (**Fig. 3D, table S1, S6**). mAF ranged from 3.5×10^−2^ - 0.18 with an average of 8.3×10^−2^. Except for patient K42 (**Fig. 1C**), SCNA were detected in all of these samples after size selection. In four patients, two or more mutations were identified and *SETD2* was the most frequently mutated gene in the cohort with a mutation being observed in 4 of the 8 mutation positive patients (50%). *KDM5C* and *VHL* were the next most frequently mutated genes with mutations being observed in 2/8 (25%) in both instances (**Fig. 1C**).

ctDNA detection across all patient samples are summarised in **Fig. 4** and **Fig. S13**. Targeted analysis with a personalised approach (INVAR-TAPAS) improved ctDNA detection over untargeted analysis. Overall (targeted and untargeted data), 24 of 47 (51.5%) DIAMOND patients had detected ctDNA, whether in plasma or urine, though not all patient samples had the most sensitive approach applied to them or had both plasma and urine available. Of those that did, 11/13 (84.6%) patients had ctDNA detected. Of note, only one patient had ctDNA detected in both plasma and urine by both untargeted and targeted methods. For the MonReC cohort the overall ctDNA detection rate of 34.5% was lower than for DIAMOND, most likely due to the use of less sensitive *de novo* methods. Taken together, ctDNA detection in plasma and urine is challenging in patients with renal tumours, with tumour size (**Fig. 3E**) and renal vein or inferior vena cava tumour thrombus locally advanced disease (**Fig. 3F**) being the single greatest factors contributing to detection (as assessed in DIAMOND, **Fig. S11A-B)**.

**Fig. 4.**
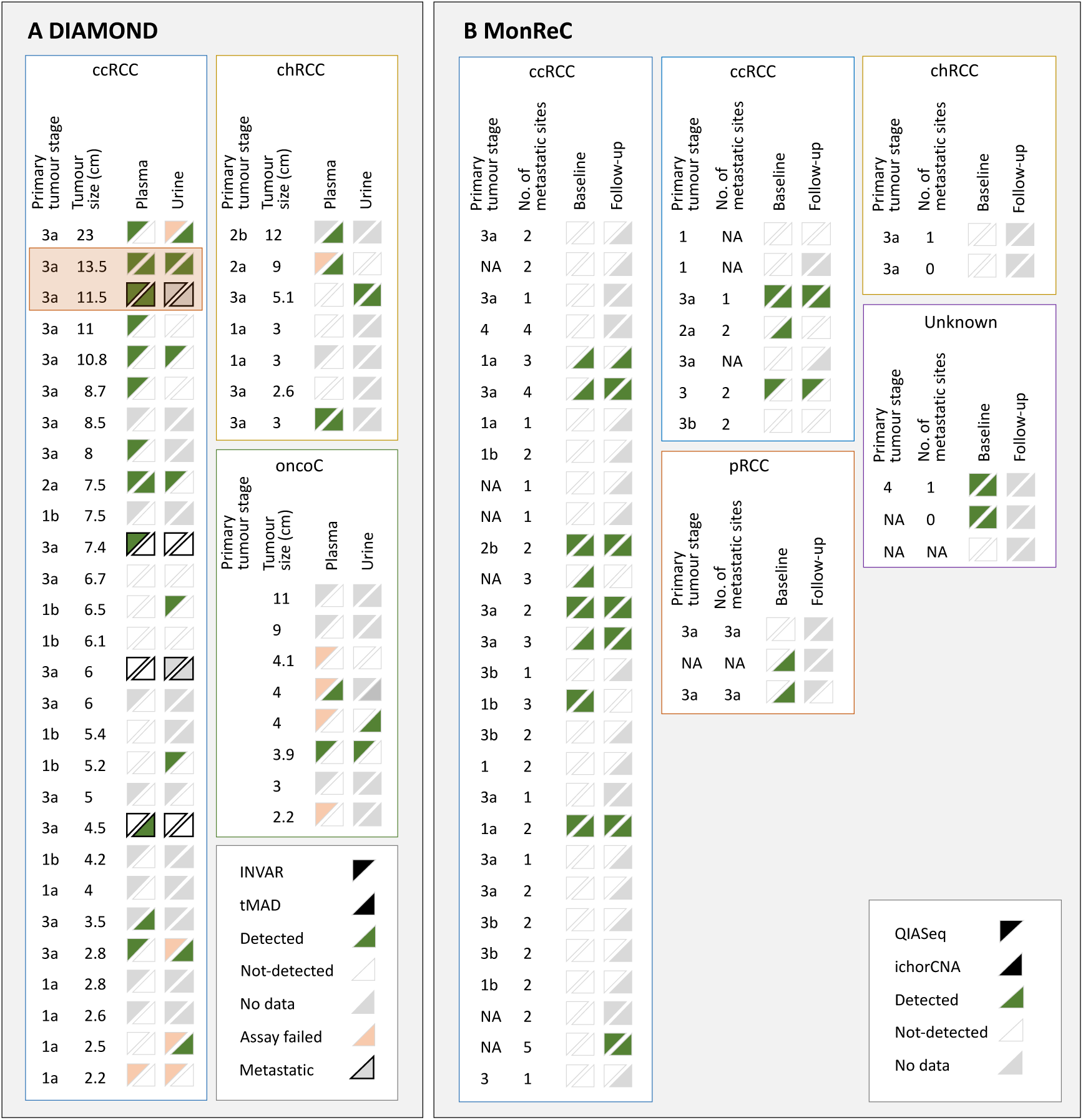
Summary of ctDNA detection in all patients and all biofluids. **(A)** Summary of ctDNA detection in baseline plasma (left of triangle box pair) and urine (right of pair) of DIAMOND ccRCC (left), chRCC (top right) and oncocytoma (oncoC, middle right) patients. Four patients with ‘other’ disease sub-types are not shown. Samples are ranked in descending order according to tumour size (cm). For each data point, the upper left triangle shows the results of INVAR-TAPAS analysis and the bottom right the results of tMAD analysis. Green triangles indicate samples in which ctDNA was detected, white triangles indicate samples in which ctDNA was not detected, grey triangles indicate no data available (because the assay was not applied to that sample, or no sample was available), and pink triangles indicate failed assay. Data points with a black outline indicate patients with metastatic disease at the time of sampling. DIAMOND patients 5842 and 5634 (longitudinal section) are highlighted with an orange box. **(B)** summary of ctDNA detection in baseline (left of triangle box pair) and follow-up (right of pair) plasma of MonReC patients. Each subtype is shown in a separate box (ccRCC, clear cell; pRCC, papillary; chRCC, chromophobe; NA, unknown) The upper left triangle shows the results of QIASeq analysis, the bottom right the results of ichorCNA analysis. Triangle colour, as above. Forty-one patients had metastatic disease and, where data was available, the number of metastatic sites is indicated. ctDNA detection are plotted alongside patient characteristics in **Fig. S13**.

### Supervised machine learning to predict patients with sufficient ctDNA for detection

Our data demonstrate that there is great interpatient variability of ctDNA detection in plasma and urine of patients with RCC. In order to triage patient samples likely to have sufficiently high ctDNA levels for subsequent analysis by more expensive and time-consuming targeted methods we applied our recently developed random forest (RF) disease classification model^[14]^ to plasma sWGS data of DIAMOND patients. Used as originally intended, the RF model enables classification of plasma DNA samples into “normal” and “cancer” based on fragmentation features of cfDNA (**Fig. S14A**). Here we compared the output of the model with the results of targeted analysis and found that amongst those samples surpassing a >50% probability of ctDNA detection threshold, 75% (9/12) had detected ctDNA in plasma by INVAR-TAPAS (**Fig. S14B**) and 91.7% (11/12) had detected ctDNA in plasma and/or urine by INVAR-TAPAS and/or tMAD. Conversely, only 36.4% (4/11) of plasma samples with <50% probability of ctDNA detection by the RF model, had ctDNA detected in any fluid by any method (p=9.4×10-3 Fisher’s exact test; **Fig. 5**). This data suggests that fragmentation features may predict which RCC patients are likely to have sufficient ctDNA (in plasma or urine) for targeted sequencing by other, more sensitive, methods.

**Fig. 5.**
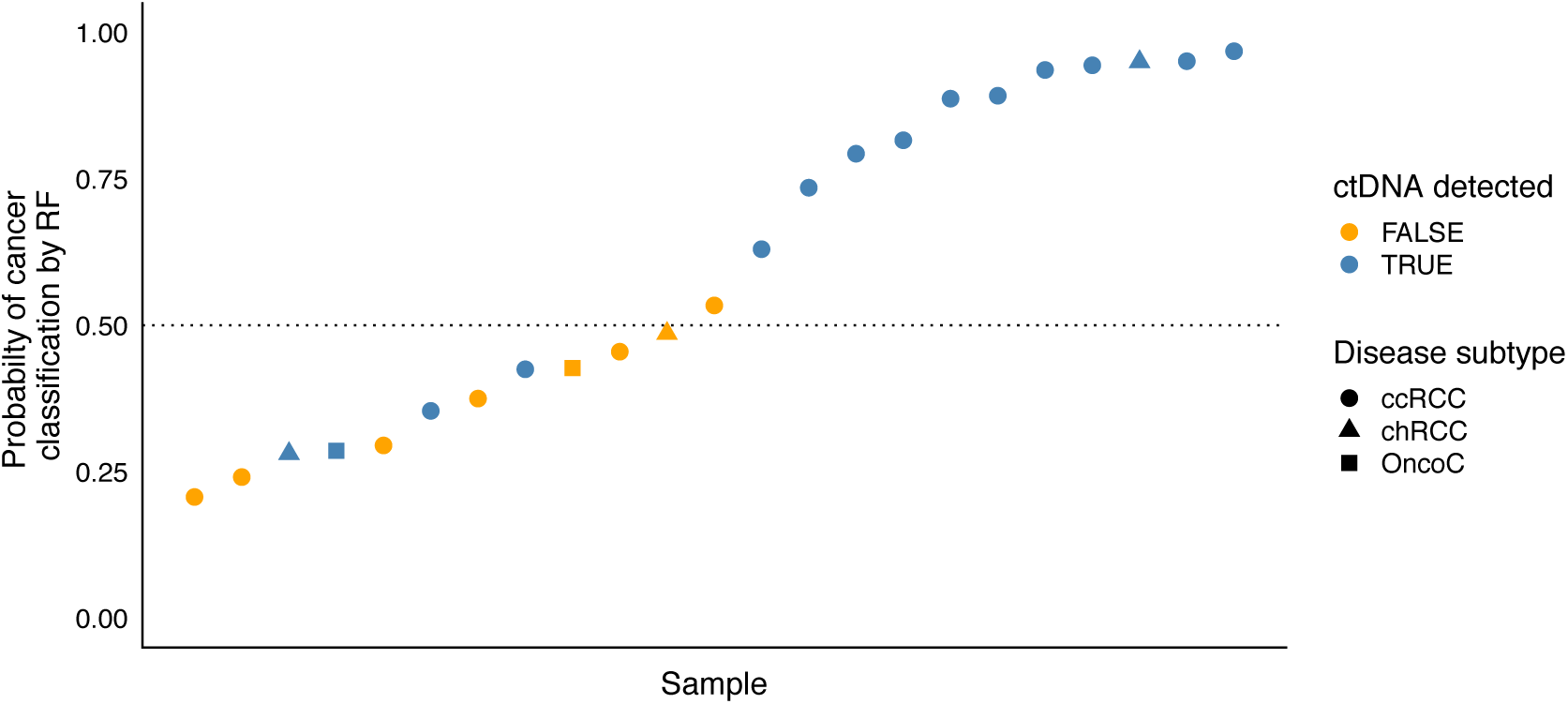
Application of a fragmentation feature based random forest model to predict patients with detected ctDNA. An RF model utilised for cancer detection based on fragmentation features^[14]^ was applied to plasma samples from DIAMOND patients. Amongst those with >50% probability of ctDNA detection (above dotted line), 11/12 (91.7%) had detected ctDNA by INVAR in either fluid. Conversely, of those with <50% probability, 4/11 (36.4%) had detected ctDNA (p=9.4×10^−3^, Fisher’s exact test). A blue point indicates patients with detected ctDNA while orange points indicate patients in which ctDNA was not detected. Disease subtype is indicated by data point shape with circle=ccRCC, triangle=chRCC and square=oncocytoma. Detection in plasma (not urine) by INVAR is shown in **Fig. S14B**. Detection in plasma (not urine) by INVAR and/or tMAD is shown in **Fig. S14C**. The equivalent data for urine are shown in **Fig. S14D-E**.

### Longitudinal analysis of ctDNA in renal tumours

An aim of the MonReC study was to investigate the potential of plasma ctDNA to monitor treatment response in metastatic RCC, as such we had access to longitudinal plasma samples for 37/43 (86%) MonReC patients. During a median follow-up period of 6 months (range 0.4-19.2) serial plasma samples (median 5, range 2-21) were collected before and during treatment. mFAST-SeqS was used as an initial measure of tumour content^[16, 25]^. For all samples the median genome-wide z-score was 1.0 (range −0.9 - 58.0), and an elevated z-score was observed in 19/252 samples (7.5%) from 9 patients. In those samples SCNA profiling revealed expected RCC aberrations, including 3p loss. Using a linear mixed model with a random intercept at the patient level, we found significant differences between baseline and treatment (Wald test, p<0.001), and progression (Wald test, p=0.0294) (**Fig. S15)**. Longitudinal mutation analysis was performed in 14 patients. Of those 6 patients (K18, K20, K23, K27, K39, K42) had mutations called at baseline (**Fig 6A-C, Fig. S16**). Due to low detection rates at baseline, we analysed additional samples whose collection coincided with clinical progression, and identified three further patients (K11, K21, K35) with detected ctDNA. Moreover, ichorCNA was applied to available follow-up samples of 5 patients (K08, K13, K19, K40, K44) with detected ctDNA at baseline (**table S7**).

**Fig. 6.**
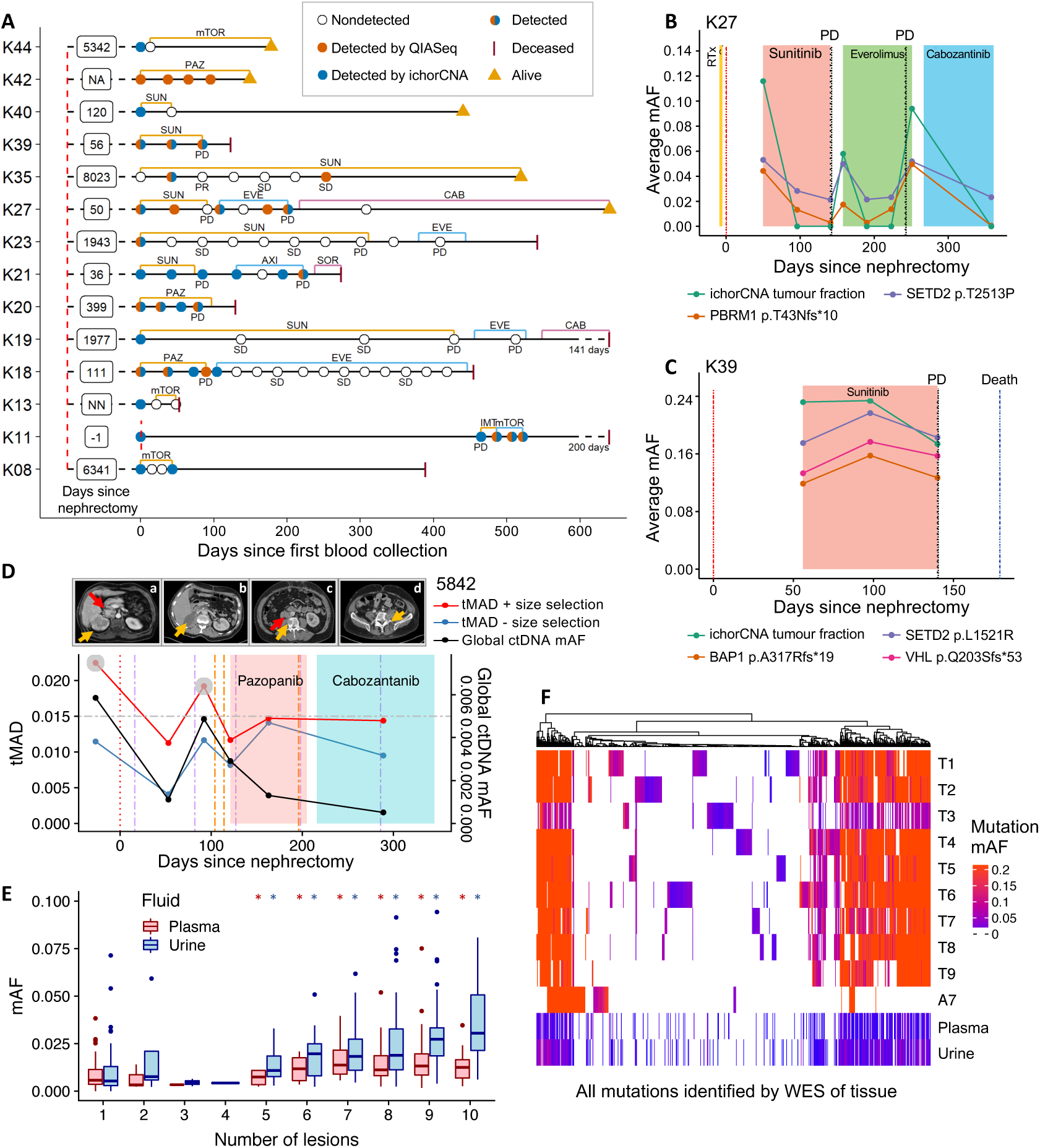
Longitudinal ctDNA analysis and assessment of intratumoural heterogeneity in plasma and urine. **(A)** Longitudinal cfDNA assessment of MonReC patients with metastatic RCC. Shown are disease courses of patients who had detected ctDNA with QIASeq and/or ichorCNA analysis. Time between nephrectomy and first blood draw is indicated in days (NA, not available; NN, no nephrectomy). Type and duration of treatment (mTOR=mTOR inhibitor; PAZ=pazopanib, SUN=sunitinib; EVE=everolimus; CAB=cabozantinib; AXI=Axitinib; SOR=sorafenib; IMT=immune therapy) are indicated by coloured lines. Most patients had detected ctDNA at progression (PD), whereas during stable disease (SD) or response (partial response, PR) ctDNA was undetected. **(B)** and **(C)** Plots demonstrating dynamic changes in ctDNA in longitudinal plasma from MonReC patients K27 and K39 respectively. Further details, and patient specific plots are in **Fig. S16**. **(D)** tMAD (left y-axis) and INVAR-TAPAS (right) analysis of plasma taken throughout the clinical course of DIAMOND patient 5842. Following nephrectomy (**scan a**; orange arrow=right renal tumour, red arrow=tumour thrombus), whilst INVAR-TAPAS global ctDNA levels (black line) drop, it remains detected (gmAF=9.5×10-4) at day 53, indicating residual disease. Conversely, imaging did not detect residual disease at day 16 (**b**; normal renal fossa). ctDNA levels rise with disease spread, before falling again upon the commencement of radio- and chemotherapy. Of note, ctDNA levels continue to fall despite evidence of clinical progression. Further details are provided in **Fig. S18A-B**. tMAD values before (blue line) and after (red line, grey circles=detected ctDNA) size selection are shown. Urine data are shown in **Fig. S18C. (E)** Comparison of baseline ctDNA mAF in plasma (red) and USN (blue) and the number of tumour regions that mutation was observed in after multi-region sampling of 5842. * indicate significant difference as compared to mutations detected in just one region. **(F)** Heatmap of mutations detected across 10 tumour biopsies (T1-T9=fresh frozen, A7=FFPE) and baseline fluid samples from 5842, with vertical coloured lines indicating individual SNVs. Hierarchical clustering was by mutation according to Euclidean distance. Colour intensity corresponds to mutation mAF. Whilst mutations show different representation in pre-surgery fluids (**Fig. S23**), all mutation clusters, even those private to individual regions, are represented by at least one mutation in plasma and urine (**Fig. S24**).

For most patients, ctDNA levels assessed either by the QIASeq panel or ichorCNA were elevated at treatment initiation but decreased with response (**Fig. 6B, Fig. S16A-G**). At progression, or when a treatment response was not gained, ctDNA increased or remained elevated (**Fig. 6C, Fig. S16A-G**). For example, patient (K39) showed high levels of tumour DNA (>0.1 mAF) at all available time points and died 2.8 months after the first blood draw, mirroring the rapidly progressing disease (**Fig. 6C**).

We obtained longitudinal plasma and urine from two DIAMOND patients (highlighted in **Fig. 4, Fig. S17**). Data are summarised in **Fig. S18-20**. ctDNA levels largely fluctuated in accordance with clinical response as determined by standard imaging. Of note, the data highlighted that the detection of ctDNA (9.5×10^−4^ gmAF) 53 days after radical nephrectomy pre-empted CT scan detection of minimal residual disease 82 days after surgery (**Fig. 6D, Fig. S17**).

### Representation of tumour heterogeneity in plasma and urine

For all but two DIAMOND patients, the low ctDNA levels precluded meaningful assessment of representation of tumour heterogeneity in fluids samples. For DIAMOND patient 5842 we carried out WES of 10 spatially distinct tumour biopsies obtained after nephrectomy (**Fig. S21**), identifying somatic mutations with varying apparent clonality. We compared the number of regions a mutation was called in, against the mAF of that mutation in plasma, urine and tissue (**Fig. 6E, fig. S22**) and observed an incrementally rising mAF as more tumour regions were considered (Wilcoxon T-test p<0.05). We assessed whether private mutations from each tumour region were represented in plasma and USN and found that both overcame this apparent heterogeneity with 90% and 100% of regions represented by at least one mutation in plasma and urine respectively (**Fig. 6F and Fig. S23**). There was no evidence of one or more tumour regions having greater representation in fluids than others (**Fig. S24**). These data confirm that plasma ctDNA can overcome tumour heterogeneity^[26]^ and, for the first time, demonstrates that USN ctDNA is capable of the same (**Fig. 6F**). The ctDNA mAF varied between plasma and urine (mean mAF of detected mutations = 2.2×10^−2^ vs 1.2×10^−2^ respectively), with differing representation of likely driver genes including VHL (ENST00000256474.2:c.333_340+1delCTACCGAGG) (**Fig. S25**). In patient 5634, plasma showed a similar ability to overcome heterogeneity with private clusters of mutations all represented in baseline plasma (**Fig. S26**).

## Discussion

Here we present the most comprehensive assessment of ctDNA in renal tumour patients to date, using state-of-the-art approaches applied to tissue and liquid biopsy samples from two independent, prospective clinical cohorts. These cohorts were complementary, with DIAMOND representing patients with the full range of renal tumours from benign to locally advanced and metastatic patients. MonReC evaluated metastatic RCC patients treated with multiple systemic therapies and longitudinal follow-up/biosampling allowing the predictive ability of ctDNA to be assessed. We sought to take a bottom-up approach to determine if inexpensive, untargeted liquid biopsy approaches could be applied to RCC, as they have been successfully employed in breast^[27]^, colorectal^[28]^ and prostate cancer^[29]^. These methods enabled ctDNA detection in only a third of patients with RCC. Even in metastatic patients these methods achieved only moderate detection rates indicating generally low levels of tumour-derived DNA. Personalised high-resolution methods, which are more expensive, were used with incremental success.

RCC is often an aggressive, angiogenesis-driven malignancy, in a vascular organ with frequent cellular necrosis. As such it is surprising that we observed such low ctDNA levels. Our data suggest that the probability of detecting ctDNA rises with increasing size of the primary tumour. Furthermore, amongst patients with locally advanced tumour growth, such as growth of a tumour thrombus into the renal vein or inferior vena cava, ctDNA detection in plasma, but not urine, was significantly more frequent. In contrast, tumour proliferation rate did not predict ctDNA detection as observed in patient derived xenograft models^[30]^ and lung cancer^[31]^. Surprisingly ctDNA detection was also limited to approximately a third of patients with metastatic disease from the MonReC cohort, albeit with substantially higher tumour fractions than observed in DIAMOND.

There is little published data characterising ctDNA levels in RCC. Initial studies suggested low detection rates and/or levels of ctDNA in locally advanced and metastatic RCC^[6, 9, 10]^. More recently targeted sequencing detected ctDNA in 30% of 53 RCC patients^[11]^. Conversely, Pal and colleagues (2017a) detected ctDNA in 78.6% of 200 metastatic patients using the Guardant360 plasma assay (Guardant Health), though with a median of one genomic alteration per sample^[32]^. The same authors detected ctDNA in a further 18/34 (53%) metastatic RCC patients, and, as echoed by our data, observed a possible correlation between detection and lesion diameter^[33]^. Likely reasons for the lower detection rate amongst metastatic patients in the MonReC cohort include our use of a smaller gene panel which has a detection limit of 5.0×10^−3^ as compared to 2.0×10^−4^ for Guardant360. Moreover, their study analysed more than 70 RCC associated genes, including *EGFR*, *NF1* and *ARID1A*, many of which were not included in our assay. Of note, neither of these two studies reported the range of detected mAFs, meaning that a direct comparison with our data was not possible. Nevertheless, the use of larger gene panels, or personalized assays such as INVAR-TAPAS, are likely to increase ctDNA detection rates. Considered with our data, it is clear that a consensus concerning ctDNA levels in RCC has yet to be reached.

For select DIAMOND patients we also assessed tumour DNA content in urine (supernatant and cell-pellet) and found similar levels and detection rates with minimal overlap between fluids, with only six patients having detected ctDNA in both plasma and urine. This data suggests that the mechanisms that determine the release and levels of ctDNA in plasma and urine of patients with renal tumours vary, a finding that requires further mechanistic analysis. We further aimed to assess intratumoural heterogeneity through the comparison of multi-region sampled tumour tissue and ctDNA. Unfortunately, analysis of further patients was hampered by their low ctDNA levels. Nevertheless, analysis of two well characterised DIAMOND patients revealed that mutations from the majority of sampled tumour regions were detected in plasma. In addition, we show for the first time that genetic heterogeneity is represented in ctDNA from urine. As such, whilst limited, our data suggest that ctDNA analysis of both fluids has the potential to overcome intratumoural heterogeneity that is prevalent in renal cancers^[33]^.

We also found that ctDNA indicated minimal residual disease after nephrectomy, pre-empting detection by imaging by 29 days. Another noteworthy finding was the detection of ctDNA in the plasma and urine of patients with benign oncocytomas and early stage ccRCCs. For the former this is particularly surprising given the benign nature of oncocytomas. Whilst differentiation of small renal masses into ccRCC, chRCC or oncocytoma can be challenging using renal tumour biopsy, these data hint at the possibility of non-invasively differentiating small renal masses to guide decisions over invasive surgery versus active surveillance.

We recognise the limitations of our study. By including a broad range of renal mass patients there was a limited number of patients with each disease stage. Furthermore, our pre-analytical knowledge evolved during recruitment to the DIAMOND cohort meaning not all sample types and time points were available for all patients.

Moreover, different techniques were applied to different patient cohorts, though this in turn reinforces the general statement that ctDNA in RCC is challenging and further developments are needed for clinical utility. Larger studies of ctDNA in ccRCC are now required to better determine its clinical utility and validate any prognostic or predictive utility. We would suggest that together with harmonised pre-analytical conditions for urine and plasma collection, a triaging method, such as the fragmentation feature based RF model explored here, is needed to move the field forward. Furthermore, assays that target multiple biomarkers, including proteins^29^ and methylated cfDNA^[34]^, will improve sensitivity. Multiple questions remain about this ctDNA-low malignancy but there is no doubt that further study is warranted and will inform approaches for other ctDNA-low tumour types.

## Methods

### DIAMOND study sample collection

Patients with a range of renal malignancies were recruited to the DIAMOND study according to local ethical guidelines (REC ID 03/018). Patient characteristics and renal tumour pathological details are presented in **Fig. 1** and **table S1**.

Patients underwent partial or total nephrectomy as part of curative treatment or cytoreductive surgery. Tumour tissue, from 29 patients, was obtained during these procedures and samples were stored as either fresh frozen-(FF) or formalin fixed paraffin embedded (FFPE) specimens. An average of 4 spatially separate tumour regions per patient (range 2 to 10 regions, 128 across all patients) were obtained, in order to study and overcome tumour heterogeneity prevalent in renal cancer^[2]^.

For FF samples, a small piece of tissue weighing <20mg was removed and DNA was extracted using a DNeasy Blood & Tissue kit (QIAGEN) according to the manufacturer’s protocol. For the FFPE samples, 2mm diameter and 3mm^3^ deep cores were obtained and DNA was extracted using the GeneRead DNA FFPE kit (QIAGEN) according to the manufacturer’s protocol, apart from the 56°C incubation step which was carried out overnight instead of for 1 hour. This protocol utilises Uracil-N-Glycosylases enzymes in order to remove artefacts resulting from the deamination of cytosine during the fixation process. All extracted DNA was quantified using the Qubit assay run on the PheraStar FSX platform (BMG LabTech).

From all DIAMOND patients we collected blood plasma prior to surgery (mean 5.0, range 0-35 days pre-surgery) and samples were processed as follows. For samples collected prior to April 2016 (21/32 patients), 8ml of blood were collected into EDTA tubes and, within 1 hour, centrifuged at room temperature at 4,000 rpm for 20 minutes. The plasma layer was subsequently decanted into separate cryotubes. The buffy coat layer was transferred into a sterile 2ml microfuge tube for parallel use. For samples collected after April 2016 (11/32 patients), 12ml of blood were collected into EDTA tubes and centrifuged at room temperature at 1,600g for 10 minutes within an hour of collection. Avoiding the buffy coat layer, 4ml of plasma was transferred to RNase-free microfuge tubes and spun on a bench top centrifuge at 13,300 rpm for 10 minutes. The supernatant was transferred to a 2ml sterile microfuge tube and the pellet was discarded. The buffy coat layer was transferred from the original collection tube into a sterile 2ml microfuge tube for parallel use. Once processed, all samples were stored at −80’C.

For 22 patients, urine samples were also collected prior to surgery (mean 8.6, range 0-35 days pre-surgery). From the same urine sample we isolated both urine supernatant (USN) and urine cell pellet (UCP), as follows. From each patient, 30-50ml urine was collected in a 50ml falcon tube, and 0.5M EDTA was added within an hour of collection (pH 8.0; 600µl for 30ml, final concentration 10mM. For larger volumes of urine the volume of EDTA was adjusted accordingly). After gentle inversion, the sample was spun at 2,400g for 10 minutes. Subsequently, ∼3.6ml of supernatant was transferred into a separate cryotube. For UCP collection, an additional 1ml of supernatant was transferred to a separate microfuge tube, whilst the remaining supernatant was discarded. The 1ml supernatant was then returned to the original falcon tube containing the urine cell pellet (UCP). This was agitated and the remaining liquid was transferred to a sterile 2ml microfuge tube. This was spun at 13,300 rpm for 10 minutes and the supernatant was discarded leaving a dry UCP for storage at −80’C.

As well as pre-surgery plasma and urine, from a subset of patients we also collected post-surgery plasma and urine (**Fig. S11**). Furthermore, in addition to renal cancer patient samples, we obtained plasma (Sera Labs) and urine DNA (local collection) from healthy individuals to act as controls for mutation analysis.

DNA was extracted from the fluid samples, as well as matched buffy coat samples, using the QIAsymphony platform (QIAGEN). DNA was quantified using the Qubit assay on the PheraStar FSX plate reader and by digital PCR (dPCR) using probes targeting the *RPP30* gene. All patient and sample details are summarised in **Fig. 1B and table S1**.

### MonReC study sample collection

An independent cohort of patients was recruited to the Graz based MonReC (monitoring renal cancer) study (approved by Ethics Committee of the Medical University of Graz, Austria, approval number 27-210 ex 14/15 and by the Ethics Committee of the Military Institute of Medicine, Warsaw, Poland, approval number 33/WIM/2015. Written informed consent was obtained from all patients before blood draw.

For the MonReC study plasma was obtained at first diagnosis of metastases, during several lines of treatment, and/or at every further instance of progression/development of new metastases along with the introduction of a new line of treatment. Patient details are summarised in **Fig. 1C** and in **table S1**.

We obtained 49 blood samples from 18 patients (mean age 62.5 years; range 46-81) from the Department of Urology and from the Division of Oncology, Department of Internal Medicine, at the Medical University of Graz, Austria. In addition, 204 plasma samples were collected from 25 patients with metastatic disease (mean age 58.9 years; range 41-68), recruited from the Department of Oncology at Military Institute of Medicine, Poland.

For the Graz cohort 9 ml blood was drawn into EDTA-containing tubes containing 10% NBF (BD Biosciences) or Streck tubes. Blood drawn at the Medical University of Graz, Austria (18/43 patients) was immediately sent to the Institute of Human Genetics. Plasma was extracted as described previously^[35]^ and stored at −80°C prior to analysis. For samples collected at the Military Institute of Medicine, Poland (25/43 patients), plasma was extracted there and stored at −80°C before shipping to Graz. cfDNA was extracted from 2 ml of plasma using QIAamp Circulating Nucleic Acid Kit (QIAGEN) according to manufacturer’s protocol. DNA was quantified using Qubit dsDNA HS Assay Kit (Thermo Fisher Scientific).

### Library preparation and exome capture of tissue and germline samples from DIAMOND patients

In order to identify patient specific somatic mutations, we carried out whole exome sequencing (WES) of all tumour tissue and germline buffy coat DNA samples. Fifty nanograms of DNA was fragmented by acoustic shearing (Covaris) according to the manufacturer’s instructions. Libraries were prepared using the Thruplex DNA-Seq protocol (Rubicon Genomics) using 5x cycles of PCR. Exome capture was performed using the TruSeq Exome Capture protocol (Illumina) with the addition of i5 and i7 specific blockers (IDT) during the hybridisation steps to prevent adaptor ‘daisy chaining’. After capture, 8x cycles of PCR were performed. Libraries, before and after hybrid capture, were quantified using KAPA library quantification kits (KAPA) and fragment size distributions were determined using a Bioanalyzer or Tapestation (Agilent). Sequencing was performed on a HiSeq 4000 (Illumina).

### DIAMOND Shallow whole genome sequencing and tMAD analysis

Shallow whole genome sequencing (sWGS) was performed on all tissue, plasma and urine (USN and UCP) samples. For each sample type, libraries were pooled in an equimolar fashion and 150bp paired end sequencing was performed (to give an average 16.4 million reads per sample) using an Illumina HiSeq 4000.

For sWGS analysis, sequence data was analysed using an ‘in-house’ pipeline that consists of the following; paired-end sequence reads were aligned to the human reference genome (GRCh37) using BWA (version 0.7.13)^[36]^ after removing any contaminant adapter sequences. SAMtools (version 1.3.1)^[37]^ was used to convert files to BAM format. PCR and optical duplicates were marked using Picard-Tools’ (version 2.2.4) ‘MarkDuplicates’ feature and these were excluded from downstream analysis along with reads of low mapping quality and supplementary alignments.

CNA calling of tissue was performed in R using the QDNAseq pipeline^[38]^. Briefly, sequence reads were allocated into equally sized (here 1 Mb and 50 kb) non-overlapping bins throughout the length of the genome. Read counts in each bin were corrected to account for sequence GC content and mappability, and bins corresponding to previously ‘blacklisted’ (ENCODE) and manually blacklisted regions were excluded from downstream analysis.

For all sWGS data, we calculated the trimmed Median Absolute Deviation (tMAD) from the copy number neutral state. This method is described in detail in Mouliere et al, 2018^[14]^). Briefly this method compares normalised read counts across genomic bins in cases against those from a cohort of healthy control samples and the median absolute deviation from log_2_R=0 of segmented bins is calculated. To define the detection threshold, we measured the tMAD score for sWGS data from 46 healthy individuals and took the maximal value (median=0.01, range 0.004–0.015). The approach has a sensitivity of 0.3%^[14]^ and details can be found at its github page (https://github.com/sdchandra/tMAD). We downsampled all plasma bam files to 10 millions reads and carried out analysis using a bin size of 30kbp. We used two forms of normalisation – 1, normalisation to a plasma sample from a cohort of healthy controls and 2, normalisation to the samples’ own mean logR. All plasma and USN samples were analysed by method one. UCP samples were analysed by method 2 as no matched healthy control samples were available for that sample type. For these UCP samples, ctDNA was detected if we observed a signal that deviated from the copy number neutral state.

### MonReC Modified Fast Aneuploidy Screening Test-Sequencing System (mFAST-SeqS)

In order to estimate the tumour fraction prior to more expensive genome-wide and/or high-resolution approaches, all samples collected in Graz were analysed using the mFAST-SeqS method. Briefly, this approach is based on the selective amplification of LINE-1 (L1) sequences. LINE-1 amplicon libraries were generated from 0.5-1ng of plasma derived DNA according to our previously published protocol^[16]^. Libraries were pooled equimolarily and sequenced on the Illumina NextSeq or MiSeq platform generating a minimum of 100,000 single-end reads for each sample. *LINE1* read counts per chromosome arm and on a genomewide level were counted and compared to a set of healthy controls and deviations were reflected in z-scores^[16]^.

### MonReC Shallow whole genome sequencing and ichorCNA analysis

For a subset of MonReC samples, sWGS was performed. To this end shotgun libraries were generated from 5-10ng of plasma DNA using the TruSeq DNA Nano Sample Preparation Kit (Illumina, San Diego, CA, USA) as previously described^[35]^. Libraries were quantified by qPCR with the quality checked using a Bioanalyzer DNA 7500 Kit (Agilent Technologies). Pooled libraries were sequenced on the Illumina NextSeq or MiSeq platform using a 2 x75bp paired-end mode. Additionally, data were analysed with the previously published ichorCNA algorithm to calculate tumour fraction from ultra-low-pass whole-genome sequencing (ULP-WGS)^[18]^. Due to the low tumour fractions, we applied an updated version of the algorithm (https://github.com/broadinstitute/ichorCNA/wiki/Parameter-tuning-and-settings). Moreover, *in silico* size selection was performed to enrich for tumour-derived fragments. As the lower limit of detection of ichorCNA was previously determined as a tumor fraction of 0.03, samples with tumor fractions below that threshold were considered as ctDNA negative. Due to the lower number of reads after size selection, the number of total reads was also downsampled to the enable comparable profiles and to exclude false positive calls (see Supplementary material). For samples with less than 2 million reads after size selection, samples were only considered as positive when SCNA frequently found in RCC according to the ProGenetix database were observed.

### DIAMOND mutations calling of tissue WES data

Mutation Calling of WES data was performed as follows;

Sequence data were aligned with BWA MEM v0.7.15^[39]^ to the GRCh37/hg19 human reference genome assembly that includes unlocalized and unplaced contigs, the rCRS mitochondrial sequence (AC:NC_012920), Human herpesvirus 4 type 1 (AC:NC_007605) and the hs37d5 decoy sequence. Duplicate read pairs based on aligned positions of each end were marked using Picard v1.122 (http://broadinstitute.github.io/picard).

We focused our mutation calling efforts on regions of the genome that had depth >20x in the matched germline buffy coat sequencing data. For this, we used the CallableLoci tool from GATK^[37]^. Somatic mutations were called using Mutect2^[40]^. In addition to Mutect2’s filters we applied additional filters described in **table S8**^[41]^.

We excluded SNVs that represented likely SNPs by virtue of their having a population allele frequency of above 0.02 in the 1000 Genomes project global database^[42]^. Furthermore, we excluded variants that had an AF > 0 in any normal adjacent tissue samples.

Despite these filters, we observed a large excess of C:G>A:T mutations in sequencing data from FFPE samples. We explored features of the artefact that could be used to distinguish it from true somatic C:G>A:T calls and found that it had a distinctive sequence context, specifically it was often preceded by C or T and G or A respectively (i.e. [C/T] C>A and [G/A] G>T). As such we filtered out all C:G>A:T calls that had this sequence context. The resulting lists of patient-specific SNVs was used to guide TAPAS panel design. WES revealed an average 297.36 unique somatic single nucleotide variants (SNV) per patient that passed our quality thresholds.

### DIAMOND Tailored Panel Sequencing (TAPAS) - custom capture panel design based on WES data

A 2.077Mb (57306 probes) personalised capture panel was designed based upon the somatic SNVs identified by WES of patient FF and FFPE tissue samples. Significant filtering of SNVs was required, as outlined above. The panel was designed using Agilent’s interactive online design tool, Sure Design (Agilent). The probe tiling parameters used were as follows; Tiling density= 1x, Masking = Most stringent, Boosting = Maximise performance, Extension into repeats = 20, Strand = sense.

As well as targeting the patient specific mutations identified by WES of patient tumour samples, the open reading frames of oncogenes and tumour suppressor genes previously implicated in renal tumourigenesis were tiled. These included the genes *VHL, PBRM1,* and *SETD2*^[2]^. The promoter region of *TERT* was also tiled^[43]^. The complete list of genes tiled by the panel are shown **in table S5**.

### DIAMOND fluid library preparation and hybrid capture

Libraries were generated from 15-30ng plasma and USN derived DNA using Thruplex Tag-Seq kits (Rubicon Genomics). Tag-Seq libraries contain unique molecular indexes (UMIs) that make it possible to trace a sequence read back to the original DNA fragment that yielded it. After 8-11 PCR cycles (dependent on input), libraries were quantified using KAPA library quantification kits. UCP DNA was sheared using acoustic shearing (Covaris) and 15-30ng were used for library preparation as above.

333.3ng each of three libraries were pooled and the 1000ng mix was used for hybrid capture using the custom SureSelect XTHS panel described above (Agilent). Hybridisation was carried out with the addition of i5 and i7 specific blockers (IDT). Captured libraries were amplified using 13 cycles of PCR. Captured libraries were quantified using KAPA library quantification kits, and sequenced across multiple HiSeq4000 (paired end 150bp) lanes such that at least 30 million sequencing reads were obtained for each sample in order to allow sufficient duplication for the proper use of UMIs.

### DIAMOND INVAR-TAPAS ctDNA detection

Aligned sequence reads were ‘collapsed’ using UMI sequences incorporated during library preparation. The CONNOR tool (https://github.com/umich-brcf-bioinf/Connor/blob/master/doc/METHODS.rst) was used, with the following settings; minimum family size = 2, requires percent majority = 90%.

We applied our INtegration of VAriant Reads - TAilored PAnel Sequencing (INVAR-TAPAS) approach to the custom capture sequence data (**Fig. S6**). Briefly, the INVAR algorithm (Wan et al, under review) aggregates signal across hundreds to thousands of mutant loci identified by WES and targeted by the custom capture panel (described above). Error suppression, through the use of UMIs, and consideration of mutation sequence context, fragment length and tumour mutant allele fraction is used to diminish background noise levels and enrich for ctDNA signal. This generates a significance level for each of the patient specific loci, which are combined into an aggregate likelihood function. Sequencing data from DNA of patients using non-matched mutation lists are used as negative controls for receiver operating characteristic (ROC) curve analysis to select a likelihood threshold for ctDNA detection. A global ctDNA allele fraction is determined by taking a background-subtracted, depth-weighted mean allele fraction across the patient-specific loci in that sample.

The sensitivity of ctDNA detection depends on the total number of informative reads (IR, unique molecules aligning to patient specific mutant loci) covering tumour-mutated loci. Here, seven plasma and six urine samples had <20,000 IR **(Fig. S9)**, resulting in limited ability to detect ctDNA. Indeed, none showed evidence of detected ctDNA. This was often due to few patient specific mutations detected in tissue (<100 mutations, **Fig. S7A)**. Detection to levels below 1.0×10^−4^ mAF would require re-analysis using greater amounts of input DNA and/or re-design of the capture panel targeting a greater number of patient specific mutations. As such, these samples were excluded as technical failures.

### MonReC QIASeq Custom panel sequencing

For mutation profiling a customized QIAseq Targeted DNA Panel (CDHS-11685Z-538; QIAGEN) was used. The panel enriches 10 genes that are frequently mutated in RCC: *BAP1*, *KDM5C*, *MET*, *MTOR, PBRM1, PIK3CA, PTEN, SETD2, TP53, VHL*. Libraries were prepared from 10ng plasma DNA according to the QIAseq targeted DNA Panel Handbook (R2). Briefly, the DNA template was enzymatically fragmented, end-repaired and A-tailed. Each DNA molecule was then tagged using an Illumina-specific adapter containing a UMI. Target regions were enriched by one target-specific primer and one universal primer. Finally, library amplification and completion of Illumina adapters was done in a universal PCR with 18 cycles. Libraries were quantified using the QIAseq Library Quant Assay (QIAGEN) and the fragment size was checked using the Bioanalyzer High Sensitivity DNA Kit (Agilent Technologies). Libraries were pooled equimolarily and sequenced on the Illumina NextSeq platform. On average 3.47 million reads (range 1.33-5.7M) were obtained per sample. Raw sequencing data generated by the QIAseq Targeted DNA Panel was analysed using the QIAseq targeted DNA Panel Analysis pipeline, which processes the UMI information to distinguish between true variants and sequencing errors based on smCounter V1^[44]^. All variants that did not pass the predefined quality criteria from smCounter were dismissed. Moreover, we filtered synonymous variants and variants present with minor allele frequencies of >1% in population frequency databases (ExAC, gnomAS, 1000g, TOPME) were considered as polymorphisms. In order to increase variant calling stringency we analysed all samples in duplicate (median raw sequencing depth 9688.3, range 4176.7-17043.9), and only considered variants to be real if identified in both replicates. All detected variants were visually checked using Integrative Genomics Viewer (IGV) (version 2.3.58). Sensitivity assessment of two independent dilution series using SeraCare reference material revealed detection rates of 90% and 100% for 0.02 mAF, 70% for 0.01 and 25% for 5.0×10^−3^ based on the evaluation of 10 different mutations. Variants with an expected mAF of 2.5×10^−3^ and 1.3×10^−3^ could not be detected.

### ctDNA detection and comparison against patient tumour size and presence of renal vein or inferior vena cava tumour thrombus

For DIAMOND patients, we assessed whether there was a difference in the distribution of tumour sizes amongst patients with detected ctDNA as compared to those in whom ctDNA was not detected. Tumour size was determined as the maximum diameter of the primary tumour on abdominal CT scan. We determined whether there was a relationship through the use of the Mann-Whitney’s U test.

Similarly, we compared ctDNA detection in patients displaying evidence of extension of a tumour thrombus into the renal vein or inferior vena cava, as assessed on cross sectional imaging, with those that did not have a tumour thrombus. Fisher’s exact test was used to determine associations between detection of ctDNA (in individual and combined fluids, by INVAR-TAPAS +/− tMAD) and tumour thrombus extension, with a significance threshold of p<0.05 (**Fig. S11**).

In both cases these were exploratory analyses and to confirm the corresponding findings and hypotheses, further confirmatory tests are required.

### Ki-67 staining of DIAMOND tissue

We compared ctDNA detection with the Ki67 cellular proliferation rate in matched tumour cells, as levels of Ki67 have previously been found to correlate well with levels of ctDNA^[31]^. We carried out immunohistochemical (IHC) staining of FFPE tissue sections from a subset of DIAMOND patients (See **Fig. S1**), using an anti-Ki67 monclonal antibody (MIB-1 clone at 1:100 dilution; DAKO Agilent Technologies LDA). The immunohistochemistry was scored manually at x400 magnification. For each slide, 20 separate high-powered fields were assessed for positively stained tumour cell nuclei by a specialist uro-pathologist (AYW). Across the 20 regions, at least 6000 tumour cells were studied for each patient. An arbitrary proportions score was used to assess Ki67 levels with regions containing up to 0%, 1%, 10%, 30%, 75% and 100% of positive cells being assigned a score of 0, 1, 2, 3, 4 and 5 respectively. The sum of these scores, across all 20 regions, was used as an indicator of the level of proliferation.

These values were subsequently compared between patients with detected and no-detected ctDNA with T-test p<0.05 indicating a significant difference (**Fig. S11C-E**).

### RF model for ctDNA detection prediction

The model used here was based upon a classification model described in Mouliere et al, 2018^[14]^. Briefly, the model considers the following fragmentation features (outlined in detail in the referenced manuscript) which were calculated from sWGS data: t-MAD, amplitude_10bp (the amplitude of 10 bp oscillations), P(20-150) (the proportion of fragments between 20 and 150 bp), P(160-180), P(20-150)/P(160-180), P(100-150), P(100-150)/P(163-169), P(180-220), P(250-320), P(20-150)/P(180-220). The model was trained using sWGS data from a cohort of ‘high ctDNA’ cancer samples, and was validated on ‘low ctDNA’ cancer samples, including plasma from the sub-cohort of DIAMOND patients used in this study. Optimal classification of samples from cancer patients and controls, was observed using a random forest (RF) machine-learning algorithm.

Here, we used the RF disease classification model to triage patient samples, predicting which RCC patients were likely to have sufficient ctDNA (in plasma or urine) for targeted sequencing by other, more sensitive, methods. We used “50% probability of cancer classification” as a threshold, comparing ctDNA detection amongst patients that fell above and below this value, as output by the RF model. We compared the ability of the model to triage patients using the INVAR-TAPAS method with or without tMAD in plasma alone, or in either fluid.

### Assessment of tumour heterogeneity and representation in plasma and urine

We assessed the heterogeneity of tumour samples from patient 5842, and its representation in plasma and urine samples obtained prior to nephrectomy. Mutations in tissue were called as described above. The SAMtools (version 1.3.1)^[37]^ function mpileup was used to assess allelic content at mutant loci, with a mAF calculated for each site and this data was converted to a matrix. A heatmap was generated from this data using the R heatmap function. Hierarchical clustering was by mutations (columns) but not by sample (rows) according to Euclidean distance.

## Acknowledgments

We would like to thank our patients and their families. This work was carried out with financial support from the CRUK Cambridge Institute (core grant, C9545/A29580), an ERC project grant (CANCER EXOMES IN PLASMA awarded to NR), an Addenbrooke’s Charitable Trust (ACT) project grant (to CS and GDS), and Renal Cancer Research Fund (to GDS). TJM is funded by a Cancer Research UK/Royal College of Surgeons of England Clinician Scientist Fellowship, C63474/A27176. Infrastructure for the DIAMOND study was provided by the Cancer Research UK Cambridge Cancer Centre and NIHR Biomedical Research Centre. The Human Research Tissue Bank is supported by the NIHR Cambridge Biomedical Research Centre. We are grateful to Elizabeth Cromwell and Mark Evans for tissue sample preparation and the genomics and bioinformatics core facilities at the CRUK Cambridge Institute.

For the MonReC study, work carried out in Graz was supported by the Austrian Science Fund FWF [P28949-B28 to EH]; and Austrian Federal Ministry for Digital and Economic Affairs (Christian Doppler Research Fund for Liquid Biopsies for Early Detection of Cancer led by EH).

## Author contributions

CGS, TM, CEM, EH and GDS conceptualised and designed the study. CGS, TM, EH and GDS wrote the manuscript. CGS, ALR and TM performed experiments and collected data. FM, DC, KH, JCMW, JM and NR developed the INVAR algorithm. CGS, FM and DC conceptualised and generated the size selection approach and the tMAD algorithm. CGS, TM, FM, DC, KH, JCMW, JM, IH, WNC devised and optimised methods applied to DIAMOND samples. CGS, TM, ME, FM, DC, KH, JB, GDS, TJM, ACPR, TFA, JNA, SJW, AM, TE recruited, treated patients and collected DIAMOND samples. ES and SU performed radiological assessment of DIAMOND patients. MP, MS, GW recruited, treated patients and collected MonReC samples and provided clinical records. AYW carried out pathological analysis. CGS, CEM, NR, EH and GDS supervised the project. All authors reviewed and approved the manuscript.

## Data and materials availability

The data that support the findings of these studies are available from the corresponding author upon request. All sequencing raw data have been deposited at the European Genome-phenome Archive (EGA; http://www.ebi.ac.uk/ega/), which is hosted by the EBI, under the accession number EGAS00001003530.

## References

1. CRUKstatistics: https://www.cancerresearchuk.org/health-professional/cancer-statistics/statistics-by-cancer-type/kidney-cancer.

2. Gerlinger M, Rowan AJ, Horswell S, Math M, Larkin J, Endesfelder D, Gronroos E, Martinez P, Matthews N, Stewart A et al: Intratumor heterogeneity and branched evolution revealed by multiregion sequencing. The New England journal of medicine 2012, 366(10):883–892.

3. Stewart GD, O’Mahony FC, Laird A, Eory L, Lubbock AL, Mackay A, Nanda J, O’Donnell M, Mullen P, McNeill SA et al: Sunitinib Treatment Exacerbates Intratumoral Heterogeneity in Metastatic Renal Cancer. Clin Cancer Res 2015, 21(18):4212–4223.

4. Turajlic S, Xu H, Litchfield K, Rowan A, Chambers T, Lopez JI, Nicol D, O’Brien T, Larkin J, Horswell S et al: Tracking Cancer Evolution Reveals Constrained Routes to Metastases: TRACERx Renal. Cell 2018, 173(3):581–594 e512.

5. Yong E: Cancer biomarkers: Written in blood. Nature 2014, 511(7511):524–526.

6. Bettegowda C, Sausen M, Leary RJ, Kinde I, Wang Y, Agrawal N, Bartlett BR, Wang H, Luber B, Alani RM et al: Detection of circulating tumor DNA in early- and late-stage human malignancies. Sci Transl Med 2014, 6(224):224ra224.

7. Heitzer E, Haque IS, Roberts CES, Speicher MR: Current and future perspectives of liquid biopsies in genomics-driven oncology. Nat Rev Genet 2019, 20(2):71–88.

8. Wan JCM, Massie C, Garcia-Corbacho J, Mouliere F, Brenton JD, Caldas C, Pacey S, Baird R, Rosenfeld N: Liquid biopsies come of age: towards implementation of circulating tumour DNA. Nature reviews Cancer 2017, 17(4):223–238.

9. Ball MW, Nathan R, Gerayli F: Long-Term Response After Surgery and Adjuvant Chemoradiation for T4 Mucinous Adenocarcinoma of the Bladder: A Case Report and Review of the Literature. Clin Genitourin Cancer 2016, 14(2):e225–227.

10. Corro C, Hejhal T, Poyet C, Sulser T, Hermanns T, Winder T, Prager G, Wild PJ, Frew I, Moch H et al: Detecting circulating tumor DNA in renal cancer: An open challenge. Exp Mol Pathol 2017, 102(2):255–261.

11. Yamamoto Y, Uemura M, Fujita M, Maejima K, Koh Y, Matsushita M, Nakano K, Hayashi Y, Wang C, Ishizuya Y et al: Clinical significance of the mutational landscape and fragmentation of circulating tumor DNA in renal cell carcinoma. Cancer Sci 2019, 110(2):617–628.

12. Dudley JC, Schroers-Martin J, Lazzareschi DV, Shi WY, Chen SB, Esfahani MS, Trivedi D, Chabon JJ, Chaudhuri AA, Stehr H et al: Detection and surveillance of bladder cancer using urine tumor DNA. Cancer Discov 2018.

13. Patel KM, van der Vos KE, Smith CG, Mouliere F, Tsui D, Morris J, Chandrananda D, Marass F, van den Broek D, Neal DE et al: Association Of Plasma And Urinary Mutant DNA With Clinical Outcomes In Muscle Invasive Bladder Cancer. Sci Rep 2017, 7(1):5554.

14. Mouliere F, Chandrananda D, Piskorz AM, Moore EK, Morris J, Ahlborn LB, Mair R, Goranova T, Marass F, Heider K et al: Enhanced detection of circulating tumor DNA by fragment size analysis. Sci Transl Med 2018, 10(466).

15. Hellwig S, Nix DA, Gligorich KM, O’Shea JM, Thomas A, Fuertes CL, Bhetariya PJ, Marth GT, Bronner MP, Underhill HR: Automated size selection for short cell-free DNA fragments enriches for circulating tumor DNA and improves error correction during next generation sequencing. PLoS One 2018, 13(7):e0197333.

16. Belic J, Koch M, Ulz P, Auer M, Gerhalter T, Mohan S, Fischereder K, Petru E, Bauernhofer T, Geigl JB et al: Rapid Identification of Plasma DNA Samples with Increased ctDNA Levels by a Modified FAST-SeqS Approach. Clinical chemistry 2015, 61(6):838–849.

17. Suppan C, Brcic I, Tiran V, Mueller HD, Posch F, Auer M, Ercan E, Ulz P, Cote RJ, Datar RH et al: Untargeted Assessment of Tumor Fractions in Plasma for Monitoring and Prognostication from Metastatic Breast Cancer Patients Undergoing Systemic Treatment. Cancers 2019, 11(8).

18. Adalsteinsson VA, Ha G, Freeman SS, Choudhury AD, Stover DG, Parsons HA, Gydush G, Reed SC, Rotem D, Rhoades J et al: Scalable whole-exome sequencing of cell-free DNA reveals high concordance with metastatic tumors. Nature communications 2017, 8(1):1324.

19. Mouliere F, Mair R, Chandrananda D, Marass F, Smith CG, Su J, Morris J, Watts C, Brindle KM, Rosenfeld N: Detection of cell-free DNA fragmentation and copy number alterations in cerebrospinal fluid from glioma patients. EMBO Mol Med 2018, 10(12).

20. Sugiyama M, Woodman A, Sugino T, Crowley S, Ho K, Smith J, Matsumura Y, Tarin D: Non-invasive detection of bladder cancer by identification of abnormal CD44 proteins in exfoliated cancer cells in urine. Clin Mol Pathol 1995, 48(3):M142–147.

21. Togneri FS, Ward DG, Foster JM, Devall AJ, Wojtowicz P, Alyas S, Vasques FR, Oumie A, James ND, Cheng KK et al: Genomic complexity of urothelial bladder cancer revealed in urinary cfDNA. Eur J Hum Genet 2016, 24(8):1167–1174.

22. Wan JCM, Heider, K., Gale, D., Murphy, S., Fisher, E., Morris, J., Mouliere, F., Chandrananda, D., Marshall, A., Gill, A.B., Chan, P.Y., Barker, E., Young, G., Cooper, W.N., Hudecova, I., Marass, F., Bignell, G.R., Alifrangis, C., Middleton, M.R., Gallagher, F. A., Parkinson, C., Durrani, A., McDermott, U., Smith, C.G., Massie, C., Corrie, P.G., Rosenfeld, N.: ctDNA monitoring to parts per million using patient-specific sequencing and integration of variant reads. Preprint at bioRxiv https://doi.org/101101/692269 2019.

23. Cancer Genome Atlas Research N: Comprehensive molecular characterization of clear cell renal cell carcinoma. Nature 2013, 499(7456):43–49.

24. Sato Y, Yoshizato T, Shiraishi Y, Maekawa S, Okuno Y, Kamura T, Shimamura T, Sato-Otsubo A, Nagae G, Suzuki H et al: Integrated molecular analysis of clear-cell renal cell carcinoma. Nature genetics 2013, 45(8):860–867.

25. Belic J, Koch M, Ulz P, Auer M, Gerhalter T, Mohan S, Fischereder K, Petru E, Bauernhofer T, Geigl JB et al: mFast-SeqS as a Monitoring and Pre-screening Tool for Tumor-Specific Aneuploidy in Plasma DNA. Advances in experimental medicine and biology 2016, 924:147–155.

26. Murtaza M, Dawson SJ, Pogrebniak K, Rueda OM, Provenzano E, Grant J, Chin SF, Tsui DWY, Marass F, Gale D et al: Multifocal clonal evolution characterized using circulating tumour DNA in a case of metastatic breast cancer. Nature communications 2015, 6.

27. Dawson SJ, Tsui DW, Murtaza M, Biggs H, Rueda OM, Chin SF, Dunning MJ, Gale D, Forshew T, Mahler-Araujo B et al: Analysis of circulating tumor DNA to monitor metastatic breast cancer. The New England journal of medicine 2013, 368(13):1199–1209.

28. Mohan S, Heitzer E, Ulz P, Lafer I, Lax S, Auer M, Pichler M, Gerger A, Eisner F, Hoefler G et al: Changes in colorectal carcinoma genomes under anti-EGFR therapy identified by whole-genome plasma DNA sequencing. PLoS genetics 2014, 10(3):e1004271.

29. Ulz P, Belic J, Graf R, Auer M, Lafer I, Fischereder K, Webersinke G, Pummer K, Augustin H, Pichler M et al: Whole-genome plasma sequencing reveals focal amplifications as a driving force in metastatic prostate cancer. Nature communications 2016, 7:12008.

30. Mair R, Mouliere F, Smith CG, Chandrananda D, Gale D, Marass F, Tsui DWY, Massie CE, Wright AJ, Watts C et al: Measurement of Plasma Cell-Free Mitochondrial Tumor DNA Improves Detection of Glioblastoma in Patient-Derived Orthotopic Xenograft Models. Cancer research 2019, 79(1):220–230.

31. Abbosh C, Birkbak NJ, Wilson GA, Jamal-Hanjani M, Constantin T, Salari R, Le Quesne J, Moore DA, Veeriah S, Rosenthal R et al: Phylogenetic ctDNA analysis depicts early-stage lung cancer evolution. Nature 2017, 545(7655):446–451.

32. Pal SK, Sonpavde G, Agarwal N, Vogelzang NJ, Srinivas S, Haas NB, Signoretti S, McGregor BA, Jones J, Lanman RB et al: Evolution of Circulating Tumor DNA Profile from First-line to Subsequent Therapy in Metastatic Renal Cell Carcinoma. Eur Urol 2017, 72(4):557–564.

33. Maia MC, Bergerot PG, Dizman N, Hsu J, Jones J, Lanman RB, Banks KC, Pal SK: Association of Circulating Tumor DNA (ctDNA) Detection in Metastatic Renal Cell Carcinoma (mRCC) with Tumor Burden. Kidney Cancer 2017, 1(1):65–70.

34. Shen SY, Singhania R, Fehringer G, Chakravarthy A, Roehrl MHA, Chadwick D, Zuzarte PC, Borgida A, Wang TT, Li T et al: Sensitive tumour detection and classification using plasma cell-free DNA methylomes. Nature 2018, 563(7732):579–583.

35. Heitzer E, Ulz P, Belic J, Gutschi S, Quehenberger F, Fischereder K, Benezeder T, Auer M, Pischler C, Mannweiler S et al: Tumor-associated copy number changes in the circulation of patients with prostate cancer identified through whole-genome sequencing. Genome medicine 2013, 5(4):30.

36. Li H, Durbin R: Fast and accurate short read alignment with Burrows-Wheeler transform. Bioinformatics 2009, 25(14):1754–1760.

37. Li H, Handsaker B, Wysoker A, Fennell T, Ruan J, Homer N, Marth G, Abecasis G, Durbin R, Genome Project Data Processing S: The Sequence Alignment/Map format and SAMtools. Bioinformatics 2009, 25(16):2078–2079.

38. Scheinin I, Sie D, Bengtsson H, van de Wiel MA, Olshen AB, van Thuijl HF, van Essen HF, Eijk PP, Rustenburg F, Meijer GA et al: DNA copy number analysis of fresh and formalin-fixed specimens by shallow whole-genome sequencing with identification and exclusion of problematic regions in the genome assembly. Genome Res 2014, 24(12):2022–2032.

39. Li H: Aligning sequence reads, clone sequences and assembly contigs with BWA-MEM. arXiv 2013, :1303.3997.

40. Cibulskis K, Lawrence MS, Carter SL, Sivachenko A, Jaffe D, Sougnez C, Gabriel S, Meyerson M, Lander ES, Getz G: Sensitive detection of somatic point mutations in impure and heterogeneous cancer samples. Nature biotechnology 2013, 31(3):213–219.

41. Alioto TS, Buchhalter I, Derdak S, Hutter B, Eldridge MD, Hovig E, Heisler LE, Beck TA, Simpson JT, Tonon L et al: A comprehensive assessment of somatic mutation detection in cancer using whole-genome sequencing. Nature communications 2015, 6:10001.

42. Genomes Project C, Auton A, Brooks LD, Durbin RM, Garrison EP, Kang HM, Korbel JO, Marchini JL, McCarthy S, McVean GA et al: A global reference for human genetic variation. Nature 2015, 526(7571):68–74.

43. Wang K, Liu T, Liu L, Liu J, Liu C, Wang C, Ge N, Ren H, Yan K, Hu S et al: TERT promoter mutations in renal cell carcinomas and upper tract urothelial carcinomas. Oncotarget 2014, 5(7):1829–1836.

44. Xu C, Ranjbar MRN, Wu Z, DiCarlo J, Wang YX: Detecting very low allele fraction variants using targeted DNA sequencing and a novel molecular barcode-aware variant caller. Bmc Genomics 2017, 18.

