## Supplementary Material for "Comprehensive characterisation of cell-free tumour DNA in plasma and urine of patients with renal tumours"

Smith et al,

Supplementary Material

**Tables and figures**

**Table S1.** Summary of patient characteristics of the DIAMOND cohort

**Table S2.** Summary of patient characteristics of the MonReC cohort

**Table S3.** Summary of samples with detected ctDNA at baseline of the MonReC cohort.

**Table S4.** Mutations identified in tissue of in VHL, PTEN, TP53, SETD2, PBRM1, and BAP1 of patients subsequently analysed with INVAR-TAPAS (n=29).

**Table S5.** Genes targeted for open reading frame sequencing**.**

**Table S6.** Mutations identified at baseline in the MonReC cohort using a QIASeq custom panel targeting 10 frequently mutated genes in RCC.

**Table S7.** Summary of all mutation data of the MonReC cohort

**Table S8.** Summary of filters applied to somatic single nucleotide variants (SNV) calls from Mutect2, for mutation calling of DIAMOND tissue samples.

**Fig. S1.** Summary of SCNA observed in matched tumour tissue from select DIAMOND patients.

**Fig. S2.** tMAD - interrogation of plasma and USN data.

**Fig. S3.** Assessment of SCNA landscape of matched tumour tissue supports their classification as oncocytoma.

**Fig. S4.** Improved detection of SCNA by *in silico* size selection.

**Fig. S5.** Comparison of the distribution of ichorCNA tumour fraction and tMAD score at baseline, and assessment of ichorCNA scores between cancer types

**Fig. S6.** Schematic explaining the INVAR-TAPAS approach.

**Fig. S7.** Summary of tumour tissue WES of the DIAMOND patients

**Fig. S8.** Tumour heterogeneity in RCC.

**Fig. S9.** Summary of global ctDNA levels relative to informative reads in INVAR-TAPAS data from patient plasma and urine

**Fig. S10.** Correlation of tumour size and ctDNA detection.

**Fig. S11.** Correlation of venous tumour thrombus and cell proliferation rates with ctDNA detection.

**Fig. S12.** Confirmation of pathological classification as oncocytoma.

**Fig. S13.** Summary of ctDNA detection in all patients and all biofluids.

**Fig. S14.** Random Forest (RF) model for plasma fragment size distributions.

**Fig. S15.** Comparison of z-score distribution at baseline, during treatment and when progression occurred.

**Fig. S16.** Longitudinal monitoring of mutations detected with the QIAseq custom panel and ichorCNA tumour fractions.

**Fig. S17.** Clinical records of select DIAMOND patients.

**Fig. S18.** Comparison of imaging data and ctDNA levels (as predicted by INVAR.TAPAS and tMAD).

**Fig. S19.** Comparison of SCNA landscape of matched tumour tissue, normal adjacent tissue, and longitudinal plasma samples from DIAMOND patient 5634.

**Fig. S20.** Comparison of imaging data and ctDNA levels (as predicted by INVAR.TAPAS and tMAD).

**Fig. S21.** Tumour map of patient 5842.

**Fig. S22.** Comparison between mAF and mutation representation in tumour lesions.

**Fig. S23.** Representation of private mutations in the baseline plasma and USN samples of patient 5842.

**Fig. S24.** Representation of mutations private to tumour regions in plasma and urine.

**Fig. S25.** Distribution of mAF of patient specific mutations in plasma and urine from patient 5842.

**Fig. S26.** Assessment of tumour heterogeneity representation in plasma from patient 5634.

**Table S1. Summary of patient characteristics of the DIAMOND cohort**

| **Patient** | **Gender** | **Surgery type** | **Surgery date** | **Days between plasma and surgery** | **Days between urine and surgery** | **Age at diagnosis** | **Disease subtype** | **Tumour size (cm)** | **Tumour stage** | **Fuhrman grade** | **Leibovich score** | **Metastatic at sampling** |
| --- | --- | --- | --- | --- | --- | --- | --- | --- | --- | --- | --- | --- |
| 5001 | F | RN | 1.10.2014 | 0 | - | 67 | ccRCC | 8 | pT3a | 2 | 4 | No |
| 5014 | F | PN | 9.10.2014 | 0 | - | 62 | chRCC | 3 | pT1a | - | - | No |
| 5030 | F | RN | 21.01.2015 | 0 | - | 76 | oncoC | 4 | - | - | - | No |
| 5047 | F | PN | 23.01.2015 | 0 | - | 59 | chRCC | 2 | pT3a | - | - | No |
| 5401 | F | RN | 14.04.2015 | 0 | -33 | 77 | ccRCC | 11 | pT3a | 2 | 5 | No |
| 5410 | M | PN | 27.04.2015 | 0 | - | 57 | chRCC | 3 | pT1a | - | - | No |
| 5532 | M | RN | 23.10.2015 | -29 | - | 62 | ccRCC | 6 | pT3a | 2 | 4 | Yes |
| 5597 | M | PN | 14.10.2015 | -1 | - | 69 | oncoC | 2.2 | - | - | - | No |
| 5603 | M | PN | 12.11.2015 | -13 | -13 | 31 | ccRCC | 2.2 | pT1a | 2 | - | No |
| 5626 | M | RN | 14.12.2015 | -18 | -18 | 76 | ccRCC | 4.5 | pT3a | 2 | 4 | Yes |
| 5627 | F | RNU | 27.11.2015 | 0 | - | 60 | chRCC | 2.6 | pT3a | - | - | No |
| 5634 | F | RN | 10.12.2015 | 0 | - | 59 | ccRCC | 11.5 | pT3a | 4 | 11 | Yes |
| 5641 | M | RN | 8.01.2016 | -22 | -22 | 71 | oncoC | 4 | - | - | - | No |
| 5644 | M | RN | 5.02.2016 | -15 | -15 | 62 | ccRCC | 7.5 | pT2a | 3 | 5 | No |
| 5790 | F | RN | 3.02.2016 | 0 | -6 | 51 | ccRCC | 6.7 | pT3a | 3 | 6 | No |
| 5799 | M | RN | 1.03.2016 | 0 | 0 | 54 | ccRCC | 6.5 | pT1b | 2 | 2 | No |
| 5801 | M | RN | 14.03.2016 | -32 | -32 | 74 | ccRCC | 10.8 | pT3a | 4 | 8 | No |
| 5802 | F | RN | 11.03.2016 | -29 | -29 | 75 | ccRCC | 2.8 | pT3a | 3 | 6 | No |
| 5813 | M | RN | 31.03.2016 | 0 | -21 | 42 | ccRCC | 8.7 | pT3a | 2 | 4 | No |
| 5818 | F | RN | 11.03.2016 | 0 | 0 | 64 | ccRCC | 7.4 | pT3a | 3 | 6 | Yes |
| 5820 | M | RN | 11.04.2016 | 0 | 0 | 76 | chRCC | 5.1 | pT3a | - | - | No |
| 5826 | F | RN | 18.04.2016 | 0 | -25 | 77 | ccRCC | 6.1 | pT1b | 3 | 4 | No |
| 5827 | F | PN | 18.04.2016 | 0 | -18 | 60 | ccRCC | 2.5 | pT1a | 2 | - | No |
| 5829 | F | RN | 22.04.2016 | 0 | -22 | 56 | oncoC | 4.1 | - | - | - | No |
| 5830 | M | RN | 11.04.2016 | 0 | 0 | 46 | chRCC | 9 | pT2a | - | - | No |
| 5834 | F | RN | 21.04.2016 | -7 | -7 | 85 | oncoC | 3.9 | - | - | - | No |
| 5842 | M | RN | 25.05.2016 | -27 | -27 | 68 | ccRCC | 13.5 | pT3a | 4 | 9 | No |
| 5845 | M | RN | 29.04.2016 | 0 | - | 62 | ccRCC | 5 | pT3a | 3 | 5 | No |
| 5846 | F | RN | 9.06.2016 | -35 | -35 | 62 | ccRCC | 23 | pT3a | 3 | 6 | No |
| 5848 | F | RN | 6.06.2016 | 0 | - | 66 | ccRCC | 5.4 | pT1b | 3 | 3 | No |
| 5998 | M | RN | 24.06.2016 | 0 | 0 | 65 | ccRCC | 5.2 | pT1b | 2 | 2 | No |
| 7035 | F | PN | 27.07.2017 | 0 | - | 54 | ccRCC | 4 | pT1a | 2 | - | No |
| 7036 | M | RN | 28.07.2017 | -11 | - | 65 | oncoC | 9 | - | - | - | No |
| 7037 | M | PN | 21.11.2017 | 0 | - | 33 | ccRCC | 4.2 | pT1b | 2 | 2 | No |
| 7046 | F | RN | 12.09.2017 | 0 | - | 65 | oncoC | 11 | - | - | - | No |
| 7051 | M | RN | 19.09.2017 | 0 | - | 63 | ccRCC | 6 | pT3a | 4 | 4 | No |
| 7056 | M | PN | 28.09.2017 | 0 | - | 50 | ORN | 2.4 | - | - | - | No |
| 7059 | M | RN | 09.11.2017 | 0 | - | 73 | ccRCC | 2.8 | pT1a | 4 | 3 | No |
| 7060 | F | RN | 3.10.2017 | 0 | - | 76 | ccRCC | 2.6 | pT1a | 3 | 1 | No |
| 7066 | F | RN | 17.10.2017 | 0 | - | 51 | ORN | 4.7 | pT3a | - | - | Yes |
| 7076 | M | RN | 24.11.2017 | 0 | - | 62 | ccRCC | 7.5 | pT1b | 3 | 3 | No |
| 7078 | M | RN | 16.11.2017 | 0 | - | 74 | ccRCC | 3.5 | pT3a | 3 | 5 | No |
| 7087 | M | PN | 15.12.2017 | 0 | - | 63 | oncoC | 3 | - | - | - | No |
| 7090 | F | RN | 06.12.2017 | 0 | - | 41 | chRCC | 12 | pT2b | - | - | No |
| 7091 | F | RN | 06.12.2017 | 0 | - | 56 | ccRCC | 8.5 | pT3a | 3 | 6 | No |
| 7092 | F | RN | 05.02.2018 | 0 | - | 30 | MiT-FTRCC | 13 | pT2b | 2 | - | No |
| 7102 | M | RN | 16.01.2018 | 0 | - | 70 | pRCC | 14 | pT3a | - | - | Yes |

PN = Partial Nephrectomy, RN = Radical Nephrectomy, RNU = Radical Nephroureterectomy, ccRCC = clear cell Renal Cell Carcinoma, chRCC = chromophone Renal Cell Carcinoma, OncoC = Oncocytoma, pRCC = papillary Renal Cell Carcinoma, MiT-FTRCC = MiT Family Translocation Renal Cell Carcinoma, ORN = Oncocytic Renal Neoplasm. Days between plasma/urine and surgery indicate the time between fluid collection and surgery, e.g. -13 indicates that the samples was collected 13 days before treatment.

**Table S2. Summary of patient characteristics of the MonReC cohort**

| **MonReC No** | **Gender** | **Age at Diagnosis** | **Surgery type** | **Date of diagnosis/ Surgery** | **Days between plasma and surgery** | **Tumour Type** | **Tumour Stage** | **Tumour Size (cm)** | **Fuhrman Grade** | **Diagnosed with metastases** | **Status at the time of first blood draw** |
| --- | --- | --- | --- | --- | --- | --- | --- | --- | --- | --- | --- |
| K01 | M | 60 | RN | 23.01.2014 | 427 | ccRCC | pT3aN1M1 | 8 | G2 | Yes | Metastases in bone and LN |
| K02 | F | 69 | NN | 09.04.2015 | 12 | ccRCC | NA | NA | NA | Yes | No nephrecotmy; metastases in lung and LN |
| K03 | F | 78 | NN | 05.03.2015 | NA | ccRCC | pT3aN0M1 | 7 | G2 | Yes | No nephrectomy; bone metastases |
| K04 | F | 74 | NN | 15.07.2015 | 1 | ccRCC | pT4N0M0 | 8.5 | G3 | Yes | No nephectomy; metastases in lungs, liver, bone and LN |
| K05 | M | 53 | RN | 29.07.2015 | -6 | NA | pT4N0M1 | 15 | G4 | Yes | Prior nephrectomy; lung metastases |
| K06 | M | 68 | RN | 07.09.2015 | -32 | pRCC | pT3aN0M1 | 11 | G4 | Yes | Prior nephrecromy; lung metastases |
| K07 | M | 54 | RN | 15.09.2015 | -12 | chRCC | pT3aN0M1 | 6.5 | G2 | Yes | Prior nephrecromy; lung metastases |
| K08 | M | 50 | RN | 14.05.1998 | 6341 | ccRCC | pT1aN0M0 | 7.5 | NA | No | Metastases in lung, LN and bone |
| K10 | M | 58 | RN | 09.11.2015 | 2 | chRCC | pT3aN0M0 | 8 | G4 | No | No metastases at first blood draw |
| K11 | M | 46 | RN | 29.12.2015 | -1 | ccRCC | pT3aN1M1 | 10.5 | G4 | Yes | Prior nephrectomy; metastases in lungs, bone and LNs |
| K12 | F | 81 | NN | 05.03.2016 | NA | NA | NA | NA | NA | No | No metastases at first blood draw, surgery denied |
| K13 | M | 67 | NN | 23.05.2016 | NA | pRCC | NA | 8 | G2 | Yes | CT punctured biopsy, metastases in lung, bone and LN |
| K14 | F | 46 | RN | 20.04.2004 | 4441 | ccRCC | pT1aN0M0 | NA | G1 | No | Metastases in lung |
| K15 | F | 57 | RN | 07.09.2015 | 71 | ccRCC | pT1bNxMx | NA | G2 | NA | Metastases in the lungs and target metastases in the liver |
| K16 | F | 58 | RN | 30.06.2004 | 4167 | ccRCC | NA | NA | G2 | NA | Relapsed tumours in the right kidney |
| K17 | M | 41 | RN | 06.06.1993 | 8230 | ccRCC | NA | NA | NA | NA | Metastases in the lungs |
| K18 | M | 68 | RN | 24.09.2015 | 111 | ccRCC | pT2bNxMx | NA | G4 | NA | Metastases in lungs and liver |
| K19 | F | 57 | RN | 16.08.2010 | 1977 | ccRCC | NA | NA | G2 | NA | Metastases in lungs, bones and sternum |
| K20 | M | 63 | RN | 16.12.2014 | 399 | ccRCC | pT3aN0Mx | NA | G3 | NA | Metastases in lungs |
| K21 | M | 65 | RN | 20.01.2016 | 36 | ccRCC | pT3aNxM1 | NA | G4 | Yes | Metastasis in the brain, thoracic lymph nodes and retroperitonum |
| K22 | M | 64 | RN | 22.05.2013 | 1080 | ccRCC | pT3bN1M1 | NA | G2 | Yes | Thoracic lymphadenopathy |
| K23 | M | 68 | RN | 14.01.2011 | 1943 | ccRCC | pT1bNxMx | NA | NA | NA | Lymph nodes |
| K24 | M | 59 | RN | 15.12.2014 | 520 | ccRCC | pT3bNxMx | NA | G3 | NA | Metastases in lung and brain |
| K25 | M | 62 | RN | 31.12.2 | 5617 | ccRCC | pT1NxMx | NA | G1 | NA | Metastases in the lungs and pelvic bones |
| K26 | M | 58 | RN | 23.09.2013 | 1008 | ccRCC | pT3aNxMx | NA | G3 | NA | Second tumour in left kidney (ccRCC, G2, pT1aNxMx), lung metastases |
| K27 | F | 61 | RN | 19.05.2016 | 50 | ccRCC | pT1aNxM1 | NA | G4 | Yes | Bone metastases |
| K28 | M | 44 | RN | 03.06.2014 | 834 | ccRCC | pT3aNxMx | NA | G3 | NA | Cutaneous metastasis |
| K29 | M | 68 | RN | 14.09.2015 | 367 | ccRCC | pT3aNxMx | NA | G4 | NA | Metastases in lungs and LN |
| K30 | M | 65 | RN | 29.12.2014 | 659 | ccRCC | pT3bNxMx | NA | G4 | NA | Metastases in lungs, local relapse of the kidney |
| K31 | M | 57 | RN | 26.07.2016 | 86 | ccRCC | pT3bN0Mx | NA | G2 | NA | Metastases in lungs and new metastases in the spine |
| K32 | M | 54 | RN | 17.07.2007 | 3383 | ccRCC | pT1bNxMx | NA | G2 | NA | Metastases in pancreas and liver |
| K33 | M | NA | NA | NA | NA | NA | NA | NA | NA | NA | NA |
| K34 | M | NA | NA | NA | NA | ccRCC | NA | NA | NA | NA | Metastases in LN and bone |
| K35 | M | 51 | RN | 09.11.1994 | 8023 | ccRCC | NA | 5.5 | G1 | No | Several metastases in bone, LN, lung, suprarenal gland, and recurrence in right kidney |
| K36 | F | NA | RN | 31.03.2016 | 229 | ccRCC | pT3N0M1 | NA | G4 | NA | NA |
| K37 | M | 58 | RN | 01.01.2016 | 333 | ccRCC | pT1 | NA | G1 | NA | NA |
| K38 | M | 61 | RN | 12.06.2008 | 3109 | ccRCC | pT1 | NA | G1 | NA | NA |
| K39 | M | 65 | RN | 24.10.2016 | 56 | ccRCC | pT3a | 7 | G3 | Yes | NA |
| K40 | F | NA | RN | 21.09.2016 | 120 | ccRCC | pT2a | NA | G4 | NA | Metastases in lung and omentum |
| K41 | M | 50 | RN | 17.04.2015 | 768 | ccRCC | pT3a | NA | G2 | NA | NA |
| K42 | M | 71 | NA | 03.11.2017 | NA | ccRCC | pT3M1 | NA | G2 | Yes | Metastases in bone and lung, secondary tumour in prostate |
| K43 | M | 73 | RN | 02.10.2017 | 129 | ccRCC | pT3bN1M1 | 9 | G3 | Yes | Metastases in liver and lung |
| [K44](http://10.200.114.102/cgi-bin/plasmadb/patient_summary.py?patientID=K44) | M | 54 | RN | 28.07.2003 | 5342 | pRCC | pT1aN0M0 | 1.3 | G3 | No | Metastases in lungs and LN |

RN = Radical Nephrectomy, ccRCC = clear cell Renal Cell Carcinoma, chRCC = chromophobe Renal Cell Carcinoma, pRCC = papillary Renal Cell Carcinoma, LN = lymph nodes, NA = Not available, NN= no nephrectomy. Days between plasma and surgery indicate the time between fluid collection and nephrectomy, e.g. 36 indicates that the samples was collected 36 days after treatment.
**Table S3. Summary of samples with detected ctDNA at baseline of the MonReC cohort.**

| **ID** | **Mutation [%]** | **ichorCNA tumour fraction [%]** | | | **tMAD** | | | **mFAST-SeqS** |  | **Mutation detected** | **SCNA detected** | **tMAD detected** | **mFAST-SeqS z-score >3** |
| --- | --- | --- | --- | --- | --- | --- | --- | --- | --- | --- | --- | --- | --- |
|  | **Mean mAF** | **all fragments** | **subsampled** | **size selection 90-150bp** | **10M 30kbp vs Control** | **2M 500kbp vs Control** | **size selection 90-150bp** | **z-score** |  |  |  |  |  |
| K02_1 | **ND** | **0,0** | **2,2** | **4,3** | **0,005** | **0,012** | **0,015** | **1,6** |  | **FALSE** | **FALSE** | **TRUE** | **FALSE** |
| K05_1 | **7,1** | **5,5** | **5,9** | **8,7** | **0,010** | **0,015** | **0,030** | **0,3** |  | **TRUE** | **TRUE** | **TRUE** | **FALSE** |
| K08_1 | **ND** | **0,0** | **2,1** | **4,7** | **0,006** | **0,011** | **NA** | **0,1** |  | **FALSE** | **TRUE** | **FALSE** | **FALSE** |
| K11_1 | **ND** | **0,0** | **1,9** | **4,4** | **0,009** | **0,011** | **0,013** | **2,4** |  | **FALSE** | **TRUE** | **FALSE** | **FALSE** |
| K12_1 | **15,1** | **12,5** | **2,6** | **10,8** | **0,019** | **0,021** | **0,028** | **3,7** |  | **TRUE** | **TRUE** | **TRUE** | **TRUE** |
| K13_1 | **ND** | **0,0** | **1,7** | **5,3** | **0,006** | **0,012** | **NA** | **1,8** |  | **FALSE** | **TRUE** | **FALSE** | **FALSE** |
| K18_1 | **5,1** | **11,5** | **11,1** | **16,1** | **0,031** | **0,033** | **0,054** | **1,5** |  | **TRUE** | **TRUE** | **TRUE** | **FALSE** |
| K19_1 | **ND** | **0,0** | **0,0** | **3,9** | **0,006** | **0,011** | **0,010** | **0,7** |  | **FALSE** | **TRUE** | **FALSE** | **FALSE** |
| K20_1 | **8,7** | **5,1** | **5,0** | **9,9** | **0,017** | **0,021** | **0,036** | **-0,6** |  | **TRUE** | **TRUE** | **TRUE** | **FALSE** |
| K21_1 | **ND** | **5,4** | **5,9** | **21,6** | **0,016** | **0,017** | **0,062** | **-0,4** |  | **FALSE** | **TRUE** | **TRUE** | **FALSE** |
| K23_1 | **5,1** | **0,0** | **0,0** | **5,3** | **0,006** | **0,011** | **NA** | **-0,4** |  | **TRUE** | **TRUE** | **FALSE** | **FALSE** |
| K27_1 | **4,9** | **8,1** | **8,3** | **11,5** | **0,025** | **NA** | **0,037** | **1,0** |  | **TRUE** | **TRUE** | **TRUE** | **FALSE** |
| K39_1 | **14,2** | **17,2** | **15,5** | **23,1** | **0,090** | **0,078** | **0,147** | **3,3** |  | **TRUE** | **TRUE** | **TRUE** | **TRUE** |
| K40_1 | **ND** | **4,0** | **4,5** | **5,6** | **0,016** | **0,016** | **NA** | **1,0** |  | **FALSE** | **TRUE** | **TRUE** | **FALSE** |
| K42_1 | **3,4** | **0,0** | **0,0** | **2,4** | **0,007** | **0,007** | **NA** | **1,5** |  | **TRUE** | **FALSE** | **FALSE** | **FALSE** |
| K44_1 | **ND** | **1,2** | **3,3** | **6,8** | **0,011** | **0,018** | **NA** | **0,1** |  | **FALSE** | **TRUE** | **FALSE** | **FALSE** |

**Table S4. Mutations identified in tissue of in *VHL, PTEN, TP53, SETD2, PBRM1, and BAP1* of patients subsequently analysed with INVAR-TAPAS (n=29).**

| **Patient** | **Disease subtype** | **Gene^1^** | **Mutation according to HGVS (RefSeq No. : cds : protein)** | **Consequence** | **Predicted impact** | **Observed in x/y tumour regions** |
| --- | --- | --- | --- | --- | --- | --- |
| 5001 | ccRCC | *VHL* | NM_000551.2:c.481C>T: p.Arg161* | stop gained | HIGH | 1/2 |
| 5047 | chRCC | *PTEN* | NM_000314.4:c.633C>A: p.Cys211* | stop gained, splice region variant | HIGH | 3/5 |
| 5047 | chRCC | *TP53* | NM_000546.4:c.1024C>T:p.Arg342* | stop gained | HIGH | 3/5 |
| 5401 | ccRCC | *PBRM1* | NM_018313.4:c.1142_1143insC:p.Val382Cysfs*10 | frameshift variant | HIGH | 1/3 |
| 5532 | ccRCC | *VHL* | NM_000551.2:c.404T>A:p.Leu135* | stop gained | HIGH | 4/4 |
| 5532 | ccRCC | *PBRM1* | NM_018313.4:c.1708_1721del:p.Ile570Hisfs*4 | frameshift variant | HIGH | 1/4 |
| 5532 | ccRCC | *PBRM1* | NM_018313.4:c.1708_1721del:p.Ile570Hisfs*4 | frameshift variant | HIGH | 3/4 |
| 5626 | ccRCC | *VHL* | NM_000551.2:c.517del:p.Glu173Argfs*29 | frameshift variant | HIGH | 1/2 |
| 5626 | ccRCC | *PBRM1* | NM_018165.4:c.637_645+6del:p.? | splice donor variant, coding sequence variant | HIGH | 2/2 |
| 5627 | chRCC | *VHL* | NM_000551.2:c.239G>A:p.Ser80Asn | missense variant | MODERATE | 2/3 |
| 5627 | chRCC | *PBRM1* | NM_018313.4:c.1600C>G: p.Arg534Gly | missense variant | MODERATE | 2/3 |
| 5634 | ccRCC | *VHL* | NM_000551.2:c.539_540del:p.Ile180Serfs*75 | frameshift variant | HIGH | 4/4 |
| 5634 | ccRCC | *PBRM1* | NM_018313.4:c.2421del: p.Pro808Leufs*17 | frameshift variant | HIGH | 2/4 |
| 5644 | ccRCC | *VHL* | NM_000551.2:c.231C>A: p.Cys77* | stop gained | HIGH | 4/6 |
| 5644 | ccRCC | *PBRM1* | NM_018313.4:c.3981del:p.Ile1329Serfs*3 | frameshift variant | HIGH | 3/6 |
| 5790 | ccRCC | *VHL* | NM_000551.2:c.169del: p.Arg58Glyfs*9 | frameshift variant | HIGH | 2/4 |
| 5790 | ccRCC | *BAP1* | NM_004656.3:c.421C>T: p.His141Tyr | missense variant | MODERATE | 2/4 |
| 5799 | ccRCC | *VHL* | NM_000551.2:c.464-1G>A:p.? | splice acceptor variant | HIGH | 5/9 |
| 5801 | ccRCC | *VHL* | NM_000551.2:c.511A>T: p.Lys171* | stop gained | HIGH | 2/6 |
| 5801 | ccRCC | *BAP1* | chr3_52440354_ACTGCCAT_A | frameshift variant | HIGH | 4/6 |
| 5802 | ccRCC | *VHL* | NM_000551.2:c.239G>A:p.Ser80Asn | missense variant | MODERATE | 2/4 |
| 5802 | ccRCC | *PBRM1* | NM_018313.4:c.1345G>T: p.Glu449* | stop gained | HIGH | 2/4 |
| 5813 | ccRCC | *VHL* | NM_000551.2:c.394_397del: p.Gln132Leufs*26 | frameshift variant | HIGH | 5/8 |
| 5813 | ccRCC | *SETD2* | NM_014159.6: c.5421_5436del: p.Pro1808Ilefs*25 | inframe deletion | MODERATE | 7/8 |
| 5813 | ccRCC | *PBRM1* | NM_018165.4:c.236+1G>T:p.? | splice donor variant | HIGH | 7/8 |
| 5818 | ccRCC | *TP53* | NM_001126115.1:c.448C>T:p.Arg150Trp | missense variant | MODERATE | 1/4 |
| 5818 | ccRCC | *VHL* | NM_000551.2:c.583C>T:p.Gln195* | stop gained | HIGH | 2/4 |
| 5818 | ccRCC | *SETD2* | NM_014159.6:c.4584A>T:p.Glu1528Asp | missense variant, splice region variant | MODERATE | 1/4 |
| 5826 | ccRCC | *VHL* | NM_000551.2:c.517del:p.Glu173Argfs*29 | frameshift variant | HIGH | 1/5 |
| 5826 | ccRCC | *SETD2* | NM_014159.6:c.1468A>G: p.Lys490Glu | missense variant | MODERATE | 4/5 |
| 5826 | ccRCC | *PBRM1* | NM_018313.4:c.3800+1G>T:p.? | splice donor variant | HIGH | 1/5 |
| 5826 | ccRCC | *PBRM1* | NM_018165.4:c.637_645+6del:p.? | splice donor variant, coding sequence variant, intron variant | HIGH | 1/5 |
| 5827 | ccRCC | *VHL* | NM_000551.2:c.287A>C:p.Gln96Pro | missense variant | MODERATE | 2/4 |
| 5842 | ccRCC | *TP53* | NM_001126115.1:c.178C>T:p.Gln60* | stop gained | HIGH | 6/10 |
| 5842 | ccRCC | *VHL* | NM_000551.2:c.332_340+1del:p.? | inframe deletion, splice region variant | MODERATE | 9/10 |
| 5846 | ccRCC | *VHL* | NM_000551.2:c.195del:p.Val66* | frameshift variant | HIGH | 1/2 |
| 5848 | ccRCC | *VHL* | NM_000551.2:c.595G>T:p.Glu199* | stop gained | HIGH | 3/3 |
| 5848 | ccRCC | *BAP1* | NM_004656.3:c.877_880del:p.Pro293Trpfs*41 | frameshift variant | HIGH | 3/3 |
| 5998 | ccRCC | *VHL* | NM_000551.2:c.263_264del:p.Trp88Serfs*43 | frameshift variant | HIGH | 3/4 |

Mutations in these genes have previously been implicated in driving renal tumourigenesis and, as such, the listed mutations likely represent driver events^1^. Heterogeneity is observed across most mutant loci – with few mutations present in all tumour regions sequenced.

**Table S5. Genes targeted for open reading frame sequencing.**

| **Gene** | **Mutation rate** | **Gene** | **Mutation rate** | **Gene** | **Mutation rate** | **Gene** | **Mutation rate** | **Gene** | **Mutation rate** |
| --- | --- | --- | --- | --- | --- | --- | --- | --- | --- |
| VHL | 40.22 | APC | 1.02 | ALK | 0.43 | RHEB | 0.19 | MAP2K2 | 0.08 |
| PBRM1 | 23.73 | NOTCH1 | 1.02 | MLH1 | 0.43 | GLYAT | 0.18 | RHOA | 0.08 |
| SETD2 | 9.28 | ROS1 | 0.94 | BRCA1 | 0.40 | HOOK2 | 0.18 | SDHD | 0.07 |
| BAP1 | 8.84 | NFE2L2 | 0.92 | PRKAG2 | 0.38 | CCND2 | 0.18 | GNAQ | 0.06 |
| TP53 | 8.37 | ERBB4 | 0.87 | FAAH2 | 0.37 | GATA3 | 0.18 | PDZK1IP1 | 0.05 |
| CTNNB1 | 6.97 | KRAS | 0.82 | SLC5A3 | 0.37 | ASB1 | 0.18 | MITF | 0.04 |
| KDM5C | 5.04 | KIT | 0.80 | FH | 0.37 | SMAD4 | 0.17 | CDKN2B | 0.04 |
| MTOR | 4.65 | BRCA2 | 0.72 | EZH2 | 0.36 | CDK4 | 0.15 | CDK6 | 0.04 |
| PTEN | 3.18 | EGFR | 0.70 | GNAS | 0.33 | RAF1 | 0.15 | GNA11 | 0.04 |
| LRP1B | 2.73 | FGFR1 | 0.65 | DDR2 | 0.32 | ESR1 | 0.15 | CCND1 | - |
| MET | 2.68 | ERBB2 | 0.60 | ZNF765 | 0.32 | PDHB | 0.14 | RIT1 | - |
| ARID1A | 2.56 | NTRK1 | 0.60 | STK11 | 0.32 | SPINT1 | 0.14 | SDCBP2 | - |
| BRAF | 2.37 | PDXDC1 | 0.60 | SMO | 0.30 | ARAF | 0.12 | SDHC | - |
| ATM | 2.24 | RET | 0.57 | PTPN11 | 0.28 | FLCN | 0.11 | TERT* | 1.29286116 |
| FAT1 | 2.01 | JAK2 | 0.56 | NRAS | 0.28 | MAP2K1 | 0.11 |  |  |
| PIK3CA | 1.76 | PDGFRA | 0.53 | FGFR2 | 0.25 | CCNE1 | 0.11 |  |  |
| TSC2 | 1.75 | HIF1A | 0.50 | BAIAP3 | 0.23 | MYC | 0.11 |  |  |
| AKAP9 | 1.75 | PLCG2 | 0.49 | SRC | 0.23 | SDHB | 0.11 |  |  |
| SMARCA4 | 1.59 | RB1 | 0.47 | HNF1A | 0.23 | MPL | 0.11 |  |  |
| RANBP2 | 1.55 | NTRK3 | 0.47 | AR | 0.22 | NPM1 | 0.11 |  |  |
| TSC1 | 1.45 | JAK3 | 0.46 | IDH2 | 0.21 | IDH1 | 0.11 |  |  |
| CDKN2A | 1.40 | TCEB1 | 0.46 | CDH1 | 0.21 | ADAP1 | 0.09 |  |  |
| SPEN | 1.38 | FGFR3 | 0.44 | AKT1 | 0.21 | MAPK1 | 0.08 |  |  |
| NF1 | 1.34 | FBXW7 | 0.43 | HRAS | 0.20 | MAPK3 | 0.08 |  |  |

The open reading frame of 109 genes were tiled for untargeted sequencing. Genes are frequently mutated in RCC (based on Catalogue of Somatic Mutations in Cancer database), have been demonstrated to be differentially expressed in a form RCC, have a published role in RCC tumourigenesis and/or have been targeted in previous studies assessing ctDNA in renal cancer^2^. *the promoter region of TERT was tiled.

**Table S6. Mutations identified at baseline in the MonReC cohort using a QIASeq custom panel targeting 10 frequently mutated genes in RCC.**

| **Patient** | **Disease subtype** | **Gene^1^** | **Mutation according to HGVS (RefSeq No. : cds : protein)** | **mAF [%]** | **Consequence** | **Impact** |
| --- | --- | --- | --- | --- | --- | --- |
| K5 | NA | *TP53* | NM_000546.5:c.701A>G:p.Tyr234Cys | **7,1** | Missense | MODERATE |
| K12 | NA | *SETD2* | NM_014159.6:c.6391C>T:p.Gln2131* | **15,1** | Stop Gain | HIGH |
| K18 | ccRCC | *VHL* | NM_551.3:c.473T>C:p.Leu158Pro | **6,3** | Missense | MODERATE |
|  |  | *SETD2* | NM_014159.6:c.4652delA:p.Asp1551Valfs*14 | **5,9** | Frameshift | HIGH |
|  |  | *PBRM1* | NM_018313.4:c.1879C>T:p.Pro627Ser | **4,4** | Missense | MODERATE |
|  |  | *PBRM1* | NM_018313.4:c.1846_1876del:p.Lys616Alafs*16 | **4,3** | Frameshift | HIGH |
| K20 | ccRCC | *KDM5C* | NM_004187.3:c.4336C>A:p.His1446Asn | **8,7** | Missense | MODERATE |
| K23 | ccRCC | *KDM5C* | NM_004187.3:c.137delT:p.Ile46Thrfs*27 | **5,1** | Frameshift | HIGH |
| K27 | ccRCC | *SETD2* | NM_014159.6:c.7537A>C:p.Thr2513Pro | **5,3** | Missense | MODERATE |
|  |  | *PBRM1* | NM_018313.4:c.127dupA:p.Thr43Asnfs*10 | **4,4** | Frameshift | HIGH |
| K39 | ccRCC | *VHL* | NM_551.3: c.606_607insT:p.Gln203Serfs*53 | **13,3** | Frameshift | HIGH |
|  |  | *SETD2* | NM_014159.6:c.4562T>G:p.Leu1521Arg | **17,5** | Missense | MODERATE |
|  |  | *BAP1* | NM_004656.3:c.947_948insTC:p.Ala317Argfs*19 | **11,9** | Frameshift | HIGH |
| K42 | ccRCC | *MTOR* | NM_004958.3:c.3239G>A:p.Arg1080His | **3,5** | Missense | MODERATE |

**Table S7. Summary of all mutation data of the MonReC cohort (please see separate Excel file)**

**Table S8. Summary of filters applied to somatic single nucleotide variants (SNV) calls from Mutect2, for mutation calling of DIAMOND tissue samples.**

| **Name** | **Filter** | **Description** |
| --- | --- | --- |
| VariantAlleleCount | VariantAlleleCount < 4 | The number of reads supporting the variant allele in the tumour sample. |
| VariantAlleleCountControl | VariantAlleleCountControl > 1 | The number of reads supporting the variant allele in the control sample. |
| DepthControl | ReadCountControl < 20 | The number of reads covering the variant position in the control sample excluding duplicates, supplementary records and reads that fall below minimum base quality and mapping quality thresholds of 10 and 1 respectively. |
| VariantBaseQualMedian | VariantBaseQualMedian < 30 | The median base quality at the variant  position of variant reads. |
| VariantMapQualMedian | VariantMapQualMedian < 40 | The median mapping quality of variant reads. |
| MapQualDiffMedian | MapQualDiffMedian > 5  or  MapQualDiffMedian < -5 | The difference in the median mapping  quality of variant and reference reads. |
| LowMapQual | LowMapQual > 0.05 | The proportion of all reads from all  samples at the variant position that have low mapping quality (less than 1). |
| StrandBias | VariantAlleleCount >= 7  and  VariantStrandBias < 0.05  and  ReferenceStrandBias >= 0.2 | VariantAlleleCount - the variant allele count, i.e. the number of reads supporting the variant allele.  The strand bias for reads covering the  variant position is the fraction of reads aligning to one strand (smaller of 2 values).  VariantStrandBias - The strand bias for variant-supporting reads.  ReferenceStrandBias - The strand bias for reads with the reference allele. |

Filters were created and thresholds chosen by assessing kernel density plots of true and false positive SNV calls for a medulloblastoma International Cancer Genome Consortium (ICGC) benchmark dataset^3^.

**Fig. S1. Summary of SCNA observed in matched tumour tissue from select DIAMOND patients**

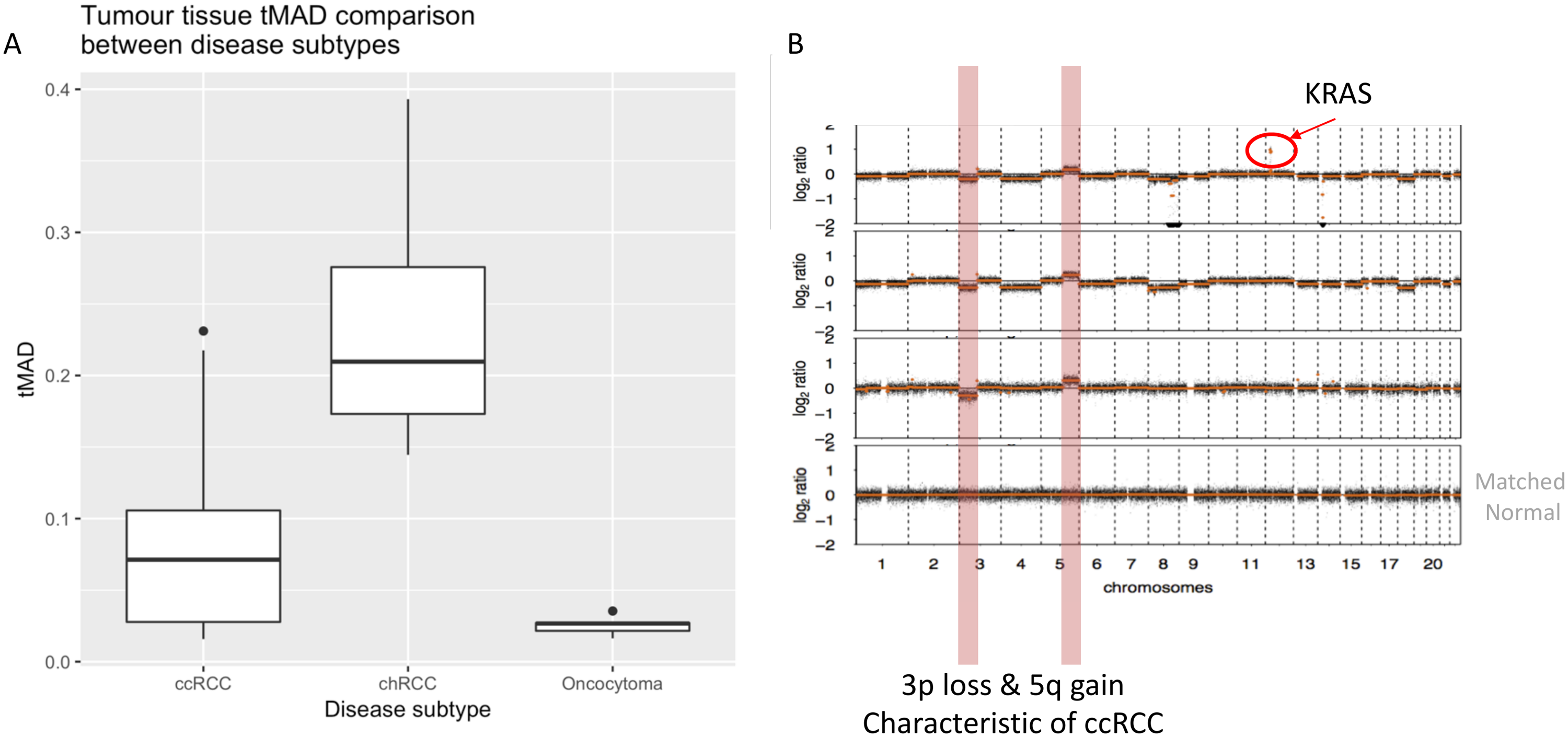

**(A)** tMAD as measured in tumour tissue samples across the different disease sub-types studied as part of DIAMOND. On average 4 (range 2-9) lesions were analysed across 19 ccRCCs, 5 chRCCs and 5 oncocytomas. chRCCs showed the greatest level of SCNA.

**(B)** Example of an SCNA profile from a patient with ccRCC. Here multiple tumour lesions were studied, along with matched normal tissue. For the majority of ccRCC tumours, including that shown here, we observed 3p loss and 5q gain that has been observed previously^1^. In addition, our analysis showed evidence of tumour heterogeneity^4^. One of the three regions studied showed evidence of a focal amplification of the KRAS oncogene. chRCC tumours showed numerous SCNA affecting chromosomes throughout the length of the genome while oncocytomas showed few SCNA, with 1p loss the most frequent change. Such SCNA patterns have previously been observed for chRCC^5,6^ and oncocytoma^7^.

**Fig. S2. tMAD - interrogation of plasma and USN data.**

**A**

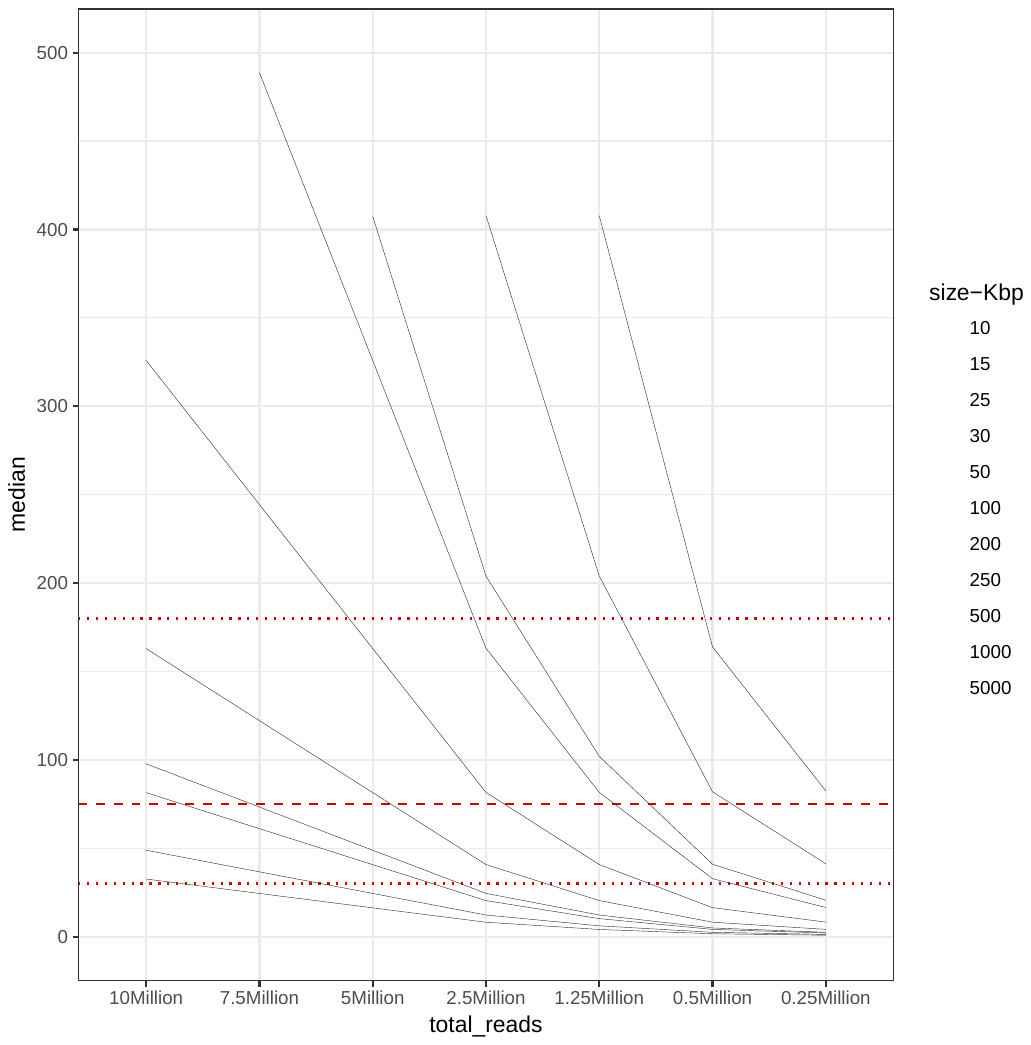

**B**

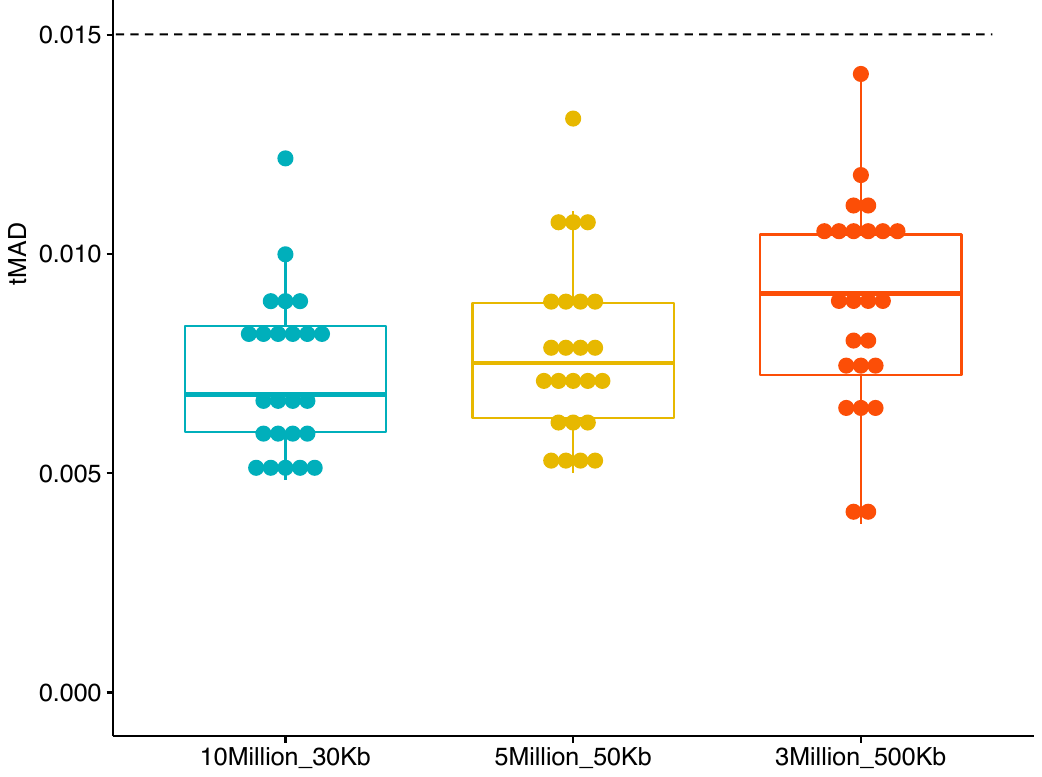

**(A)** We assessed the ability of size selection to enrich for ctDNA signal. Bam files from samples pre size selection were downsampled to 10 million reads and read counts were assessed in 30kbp bins. After size selection the number of reads was reduced and all bam files were downsampled to 2 millions reads. In order to account for this reduction in reads, we increased the bin size to 500kbp, as supported by *in silico* tests which highlights a sliding window principle for tMAD analysis of sWGS data^8^.

**(B)** For the first time we applied tMAD analysis to urine cell-free DNA data. We initially assessed the tMAD scores amongst healthy control samples, comparing values generated using different read counts and bin sizes, in order to determine a detection threshold. This threshold was set at 0.015, the same as used for plasma analysis.

**Fig. S3. Assessment of SCNA landscape of matched tumour tissue supports their classification as oncocytoma**

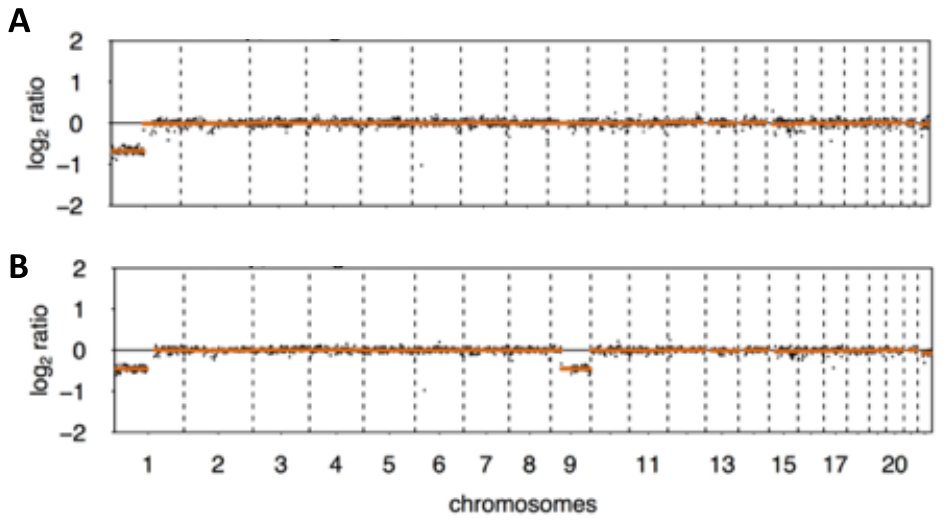

Two patients with apparent oncocytoma had ctDNA detected, by tMAD, in plasma and urine respectively. We aimed to confirm that these patients did indeed have oncocytoma by checking the SCNA landscape of matched tumour tissue from each (**A** and **B** respectively). In both instances, SCNA were observed that were characteristic of oncocytoma (loss of chromosome 1p)^7^. Shown are SCNA traces from one of multiple tissue samples available for each patient – all other available tissue samples shared the same copy number landscape.

**Fig. S4. Improved detection of SCNA by *in silico* size selection of ichorCNA data.**

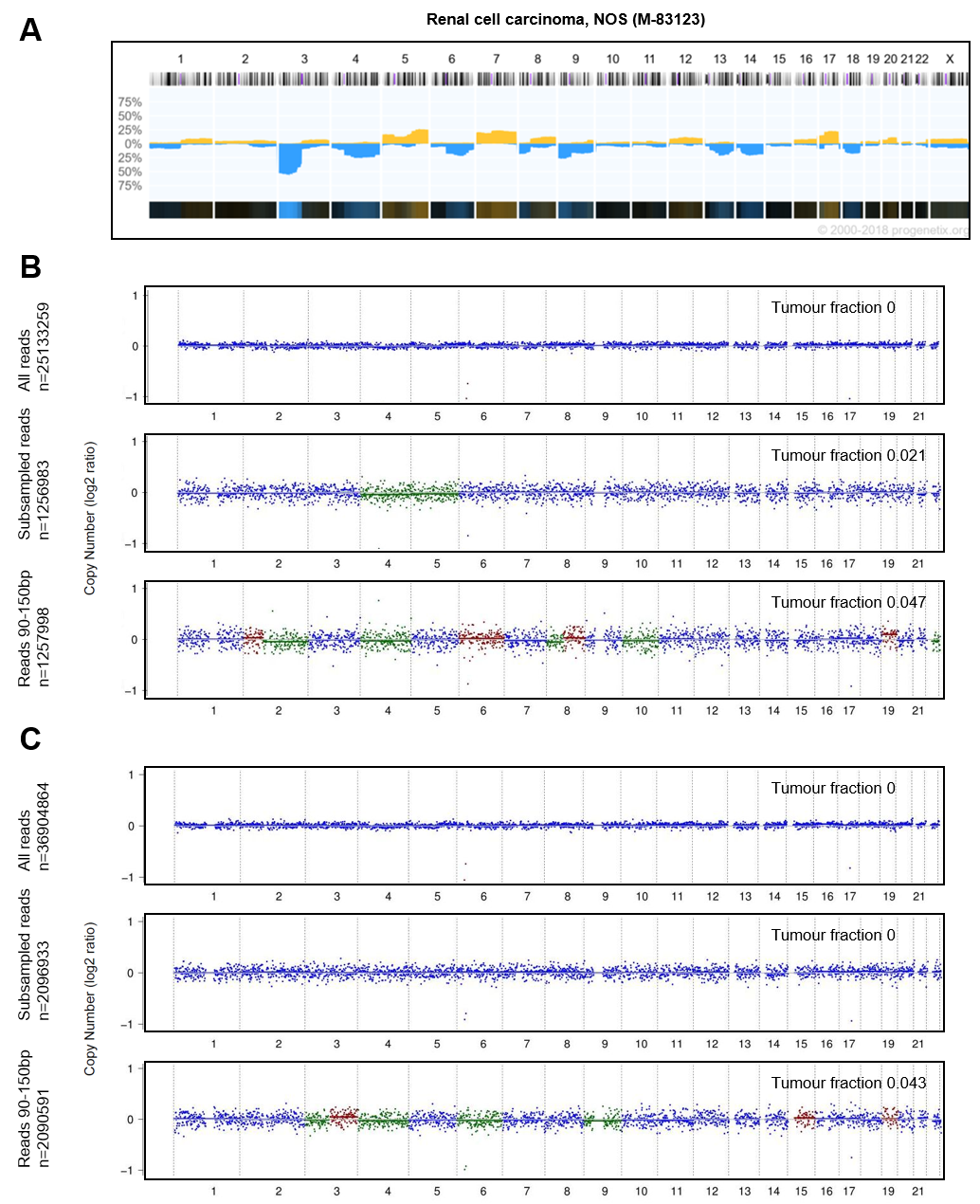

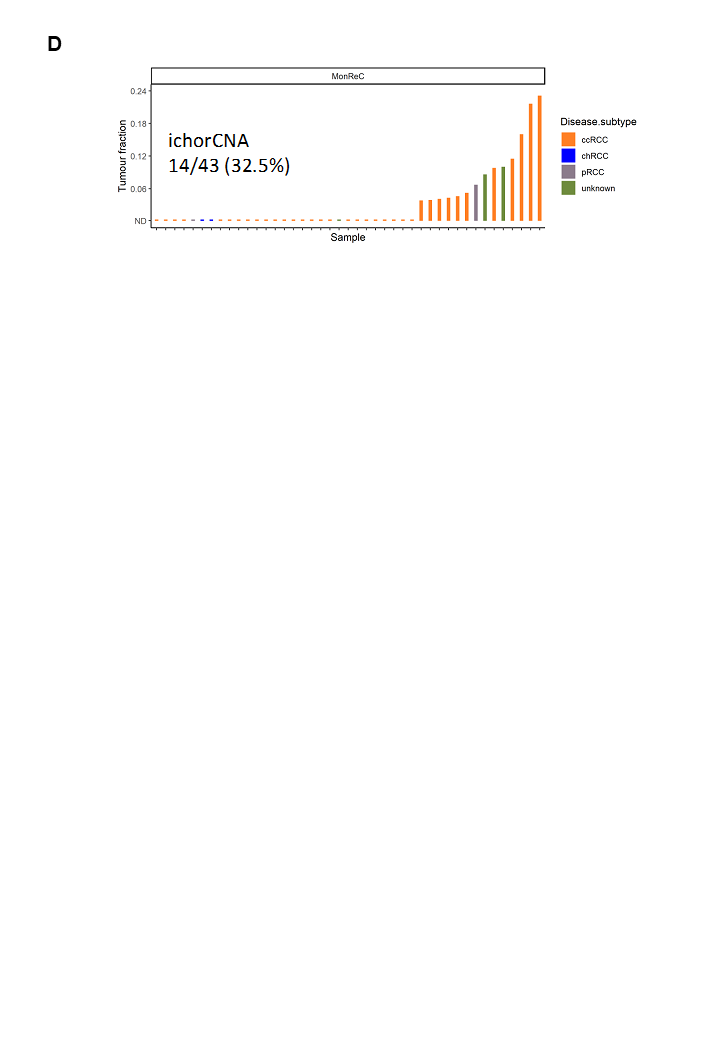

**(A)** Frequently observed SCNAs in renal cell carcinoma (n=314) from the Progenetix database. Blue regions indicate areas with copy number loss. Yellow regions indicate copy number gain.

**(B)** Data from two exemplary case are shown (**B**: K08_1 and **C**: K13_1). No SCNAs were detected at baseline using all available reads (upper panel). *In silico* size selection of reads with fragment sizes of 90-150bp led to an increase in tumour fraction, and detected SNCAs commonly observed in RCC (lower panel). We confirmed that these SCNA did not simply arise as a result of noise introduced by the lower read count by randomly downsampling the initial sequence data to a read count equivalent to that attained after size selection (middle panel). In silico selection of random reads and fragments 90-150bp from healthy control samples did not lead to an increase in the proportion of false-positives (data not shown). **(C)** In silico size selection of random reads (middle panel) and fragments 90-150bp (lower panel) from one healthy control sample did not lead to an increase in the proportion of false-positives. (D) Summary of ichorCNA analysis of patient plasma from MonReC.

**Fig. S5. Comparison of the distribution of ichorCNA tumour fraction and tMAD score at baseline, and assessment of ichorCNA scores between cancer types**

A

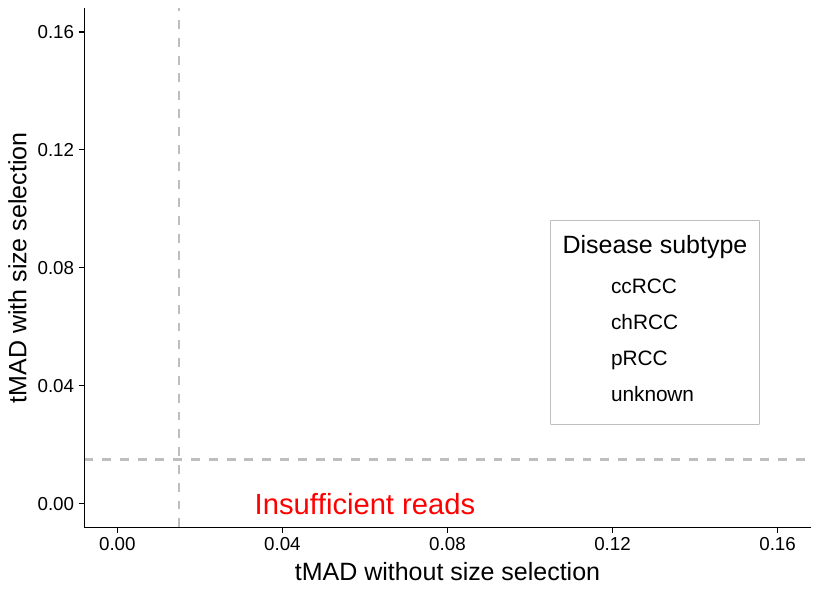

B

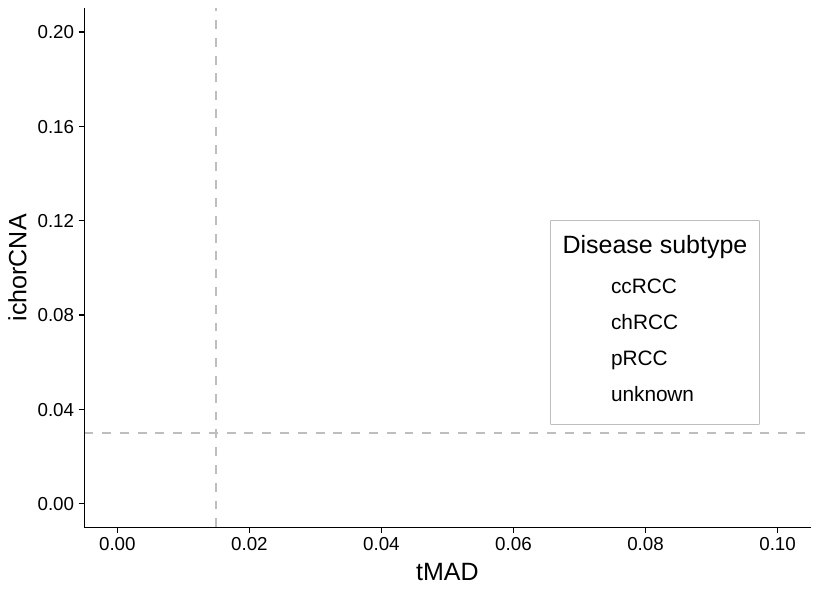

**C**

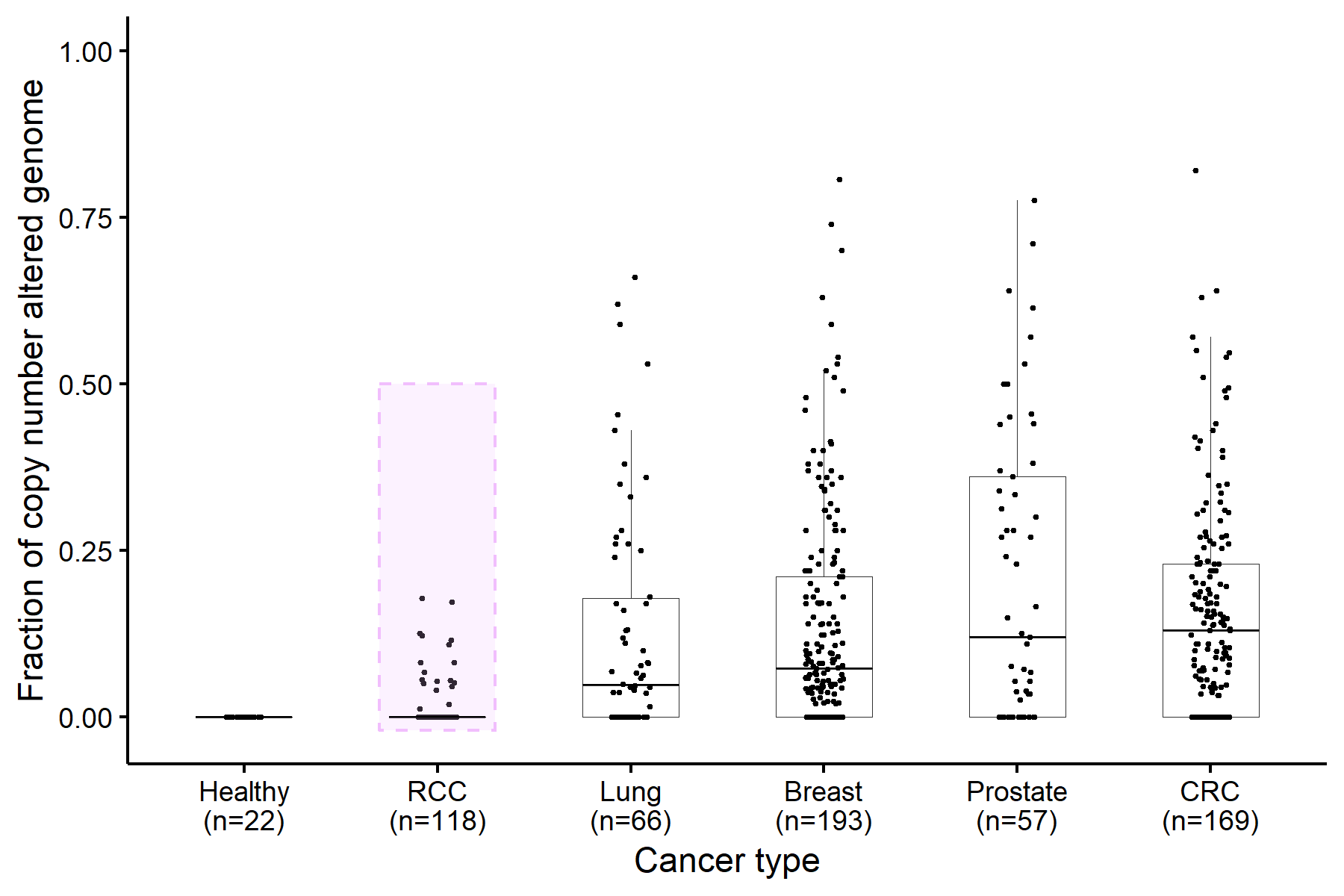

**(A)** *In silico* size selection of reads with fragment length of 90-150 bp led to an increased tMAD score and 2 additional sample surpassed the threshold of 0.015. tMAD values before (x-axis) and after (y-axis) size selection are shown.

**(B)** Comparison of tMAD and ichorCNA tumour fraction distribution without size selection.

**(C)** In addition to comparing the comparing tMAD and z-score distribution between RCCs and other cancer types (**Fig 2 D and E** respectively), we also compared the tumour fraction distribution of RCC samples, as inferred by ichorCNA, against that of other cancer types collected at the Medical University of Graz.

**Fig. S6. Schematic explaining the INVAR-TAPAS approach.**

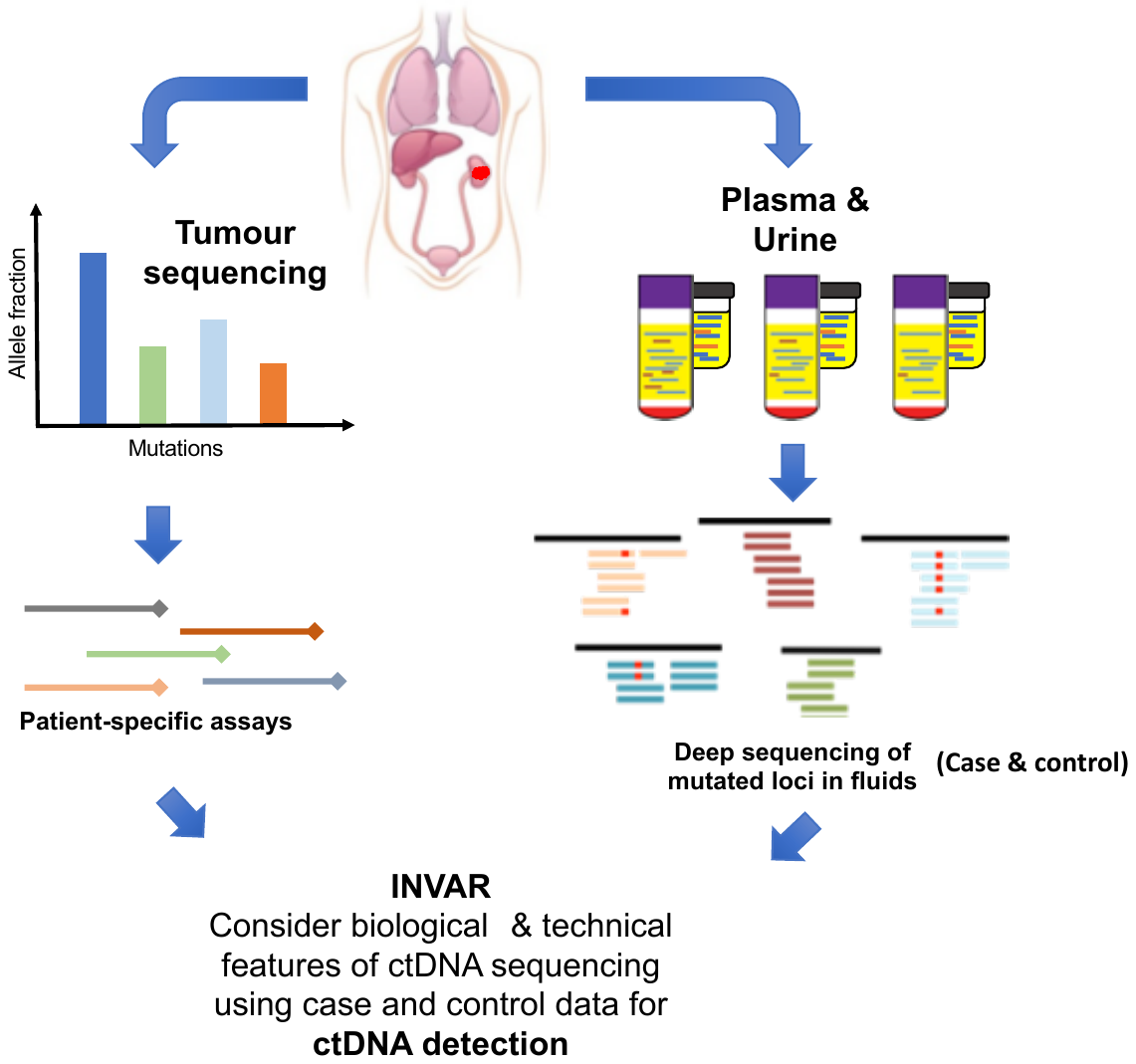

Patient specific mutations were identified through sequencing of matched tumour tissue and buffy coat. These mutations were targeted through the use of custom capture panels. By targeting very many somatic mutations, the probability of detecting a mutation in the fluid of interest is greatly increased. The custom capture panel was applied to fluid samples (here plasma and urine) and deep sequencing of each locus was carried out. We used a case-control set up to determine background error rates of sequencing and used a variety of technical features to reduce it. These included the use of unique molecular indexes (UMIs), locus noise filters, outlier suppression and the requirement for mutations to have been observed on both strands. We used probability weighting of signal based on fragment size, as well as the mutant allele fraction of mutations in the tumour tissue. For each patient specific locus we generated a significance score that were combined into an aggregate likelihood function. This was ultimately used for determining ctDNA detection and levels.

**Fig. S7. Summary of mutations targeted by INVAR-TAPAS**

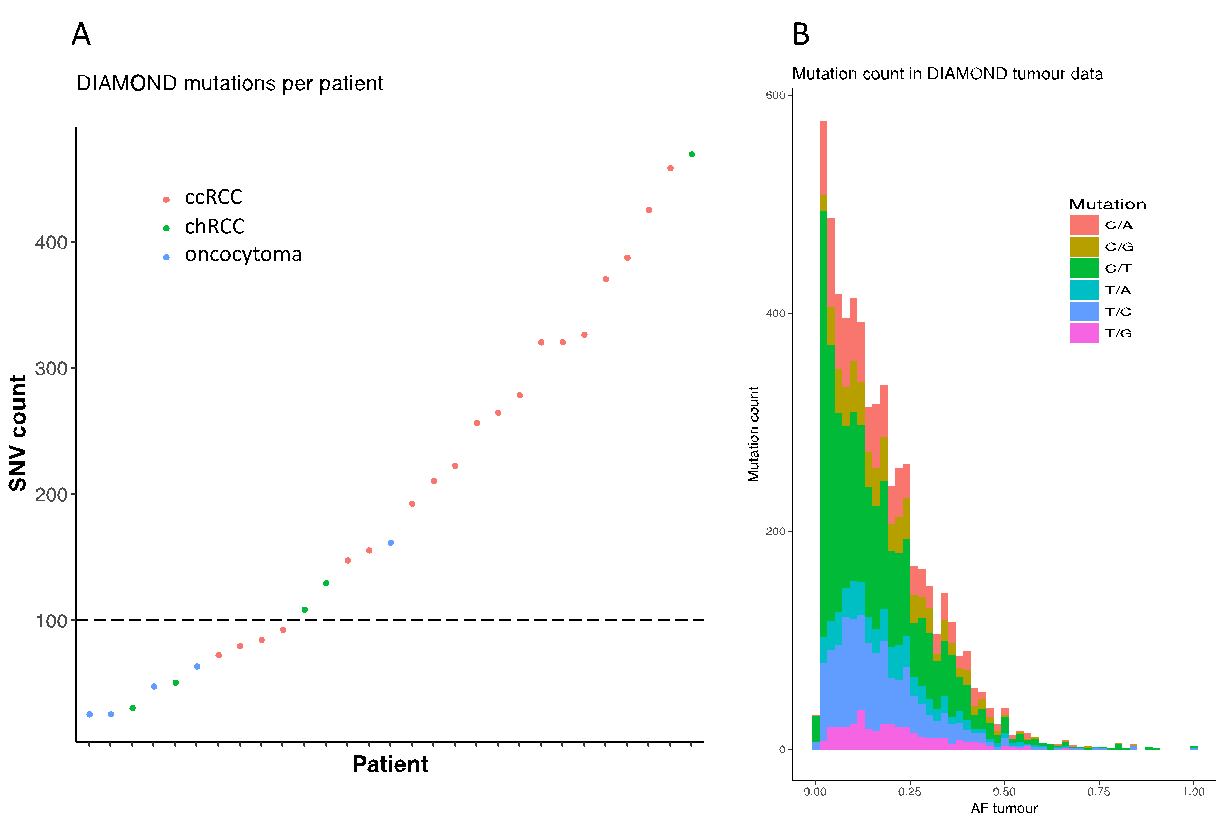

**C	D	E**

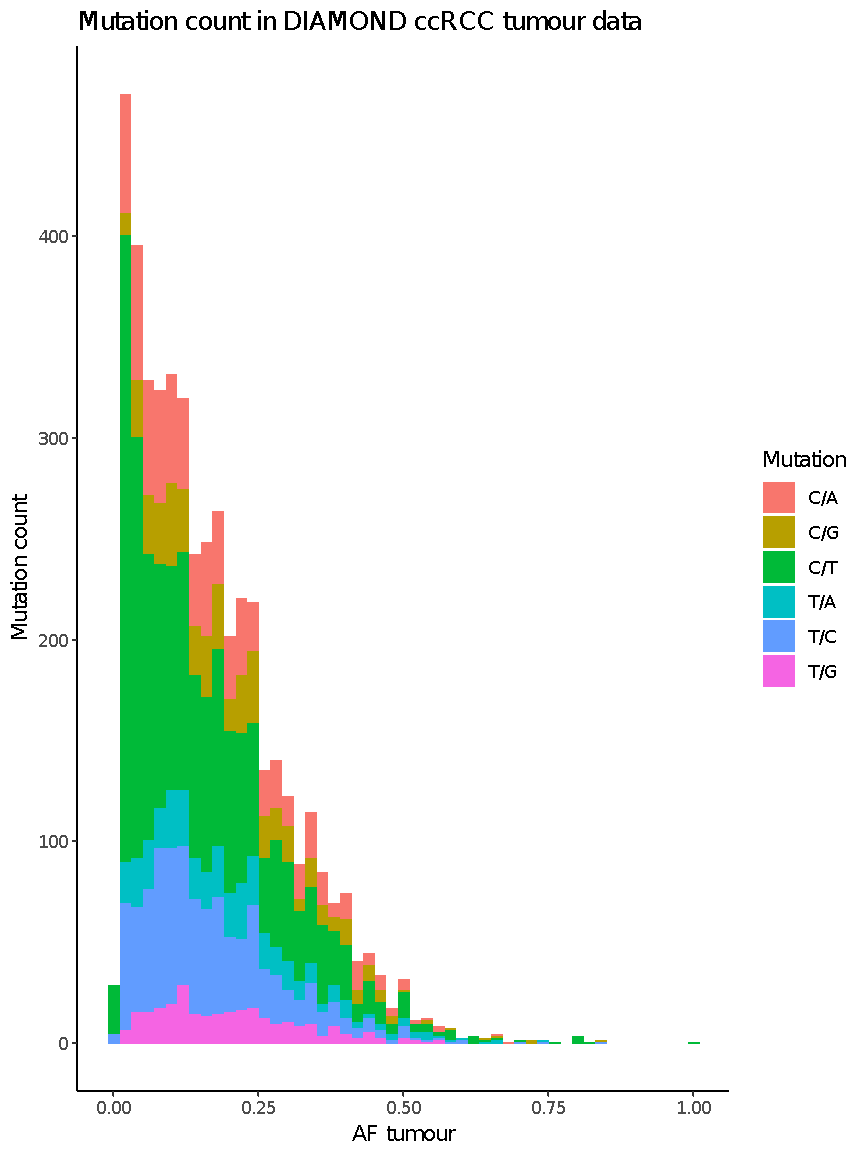

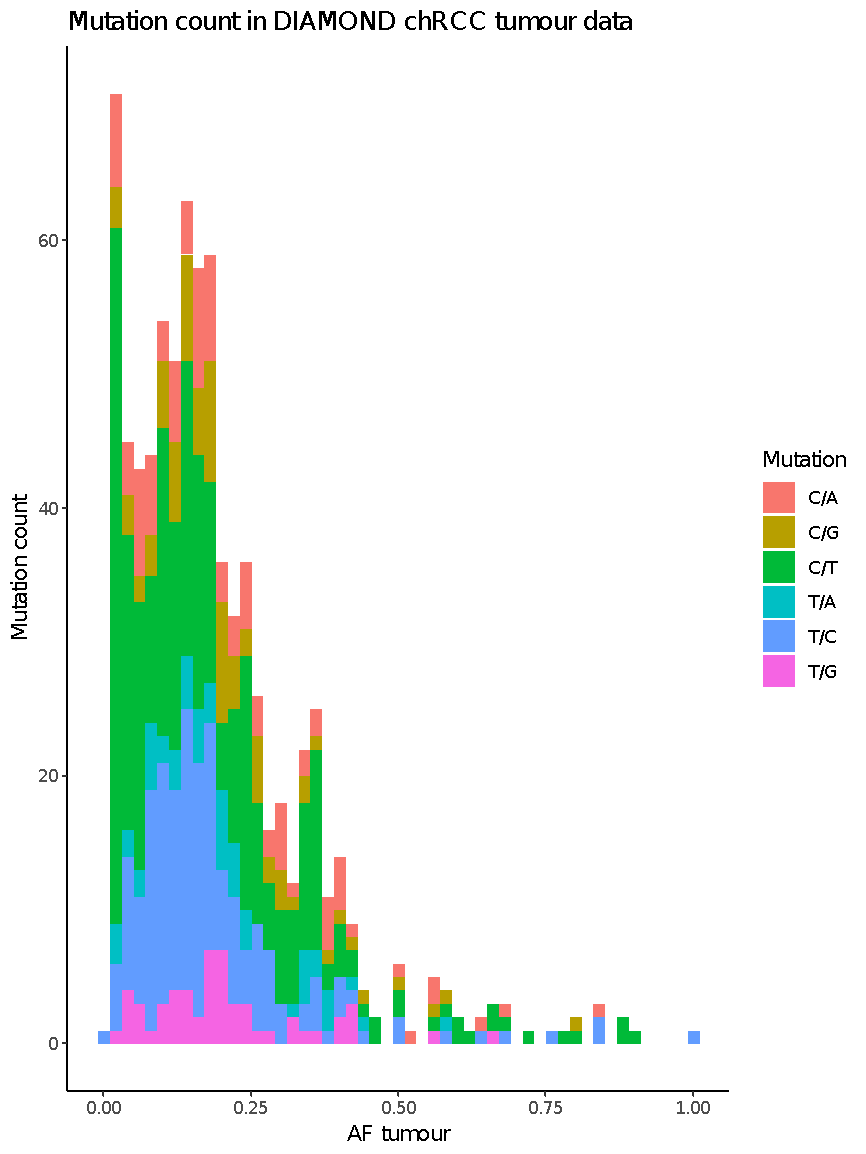

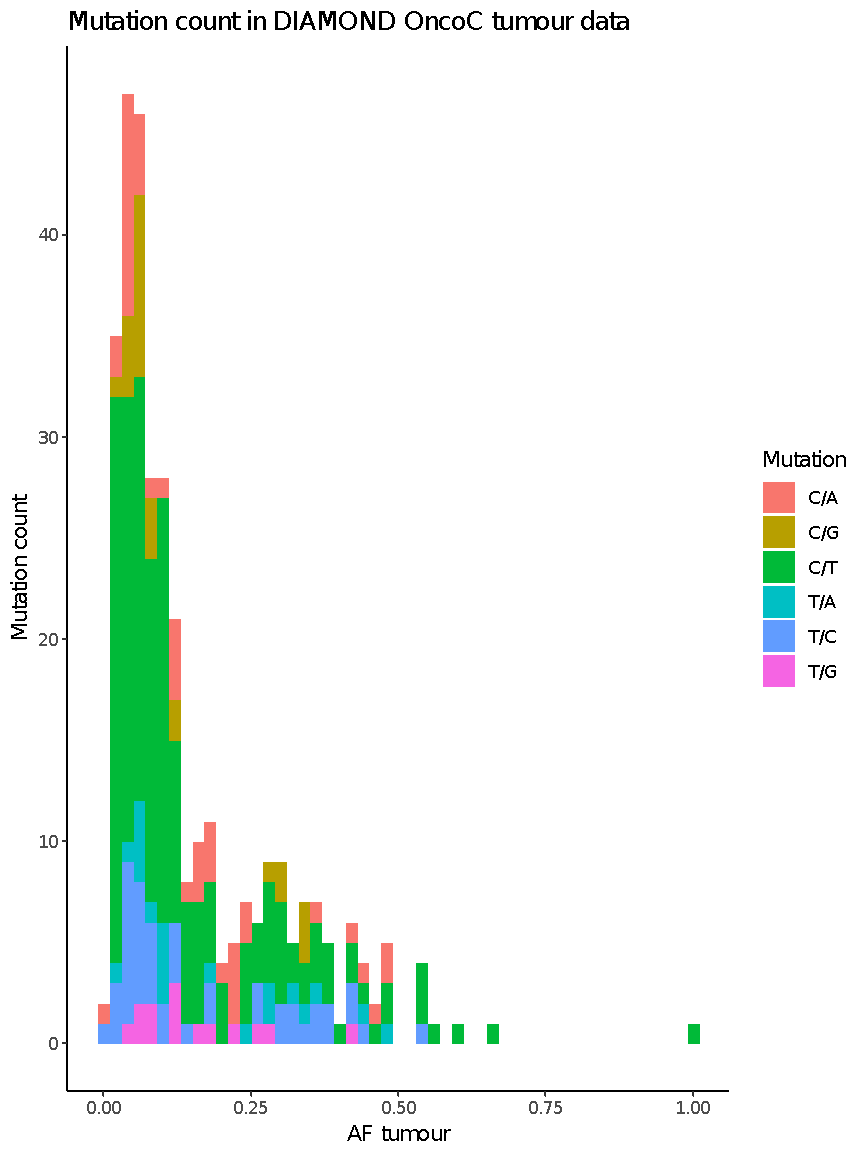

**(A)** Summary of the number of somatic SNVs identified in each of the 29 DIAMOND patients studied. On average 199.8 SNVs were targeted for each patient. Considering disease subtypes, 358.1, 158.2 and 65.2 SNVs were targeted for patients with ccRCC, chRCC and oncocytoma respectively.

**(B)** Breakdown of the SNV classes observed across the 29 patients studied. The predominant SNV class observed was C>T mutations. **(C-E)** As above but split between ccRCC, chRCC and oncocytomas respectively.

**Fig. S8. Tumour heterogeneity in RCC – example data from one patient**

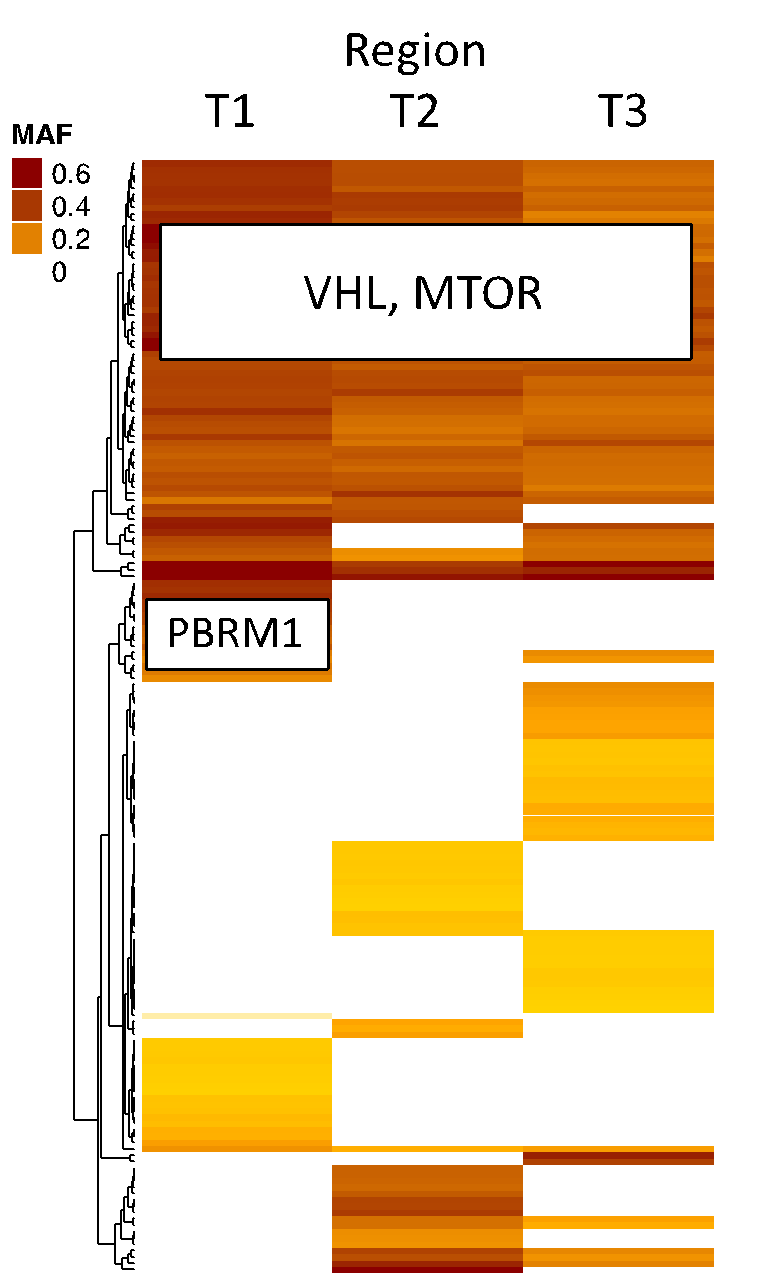

Data demonstrating the heterogeneity observed in SNV calls between tumour biopsies taken from different regions of the same tumour – here, three separate regions (labelled T1-3) were biopsied. Some SNVs were observed in all three lesions, including likely driver mutations of *VHL* and *MTOR*. However, SNV private to 1 or 2 lesions were also called. Of note, these ‘phylogenetic branches’ included mutations of key renal cancer genes including *PBRM1*.

**Fig. S9.** **Summary of global ctDNA levels relative to informative reads in INVAR-TAPAS data from patient plasma and urine**

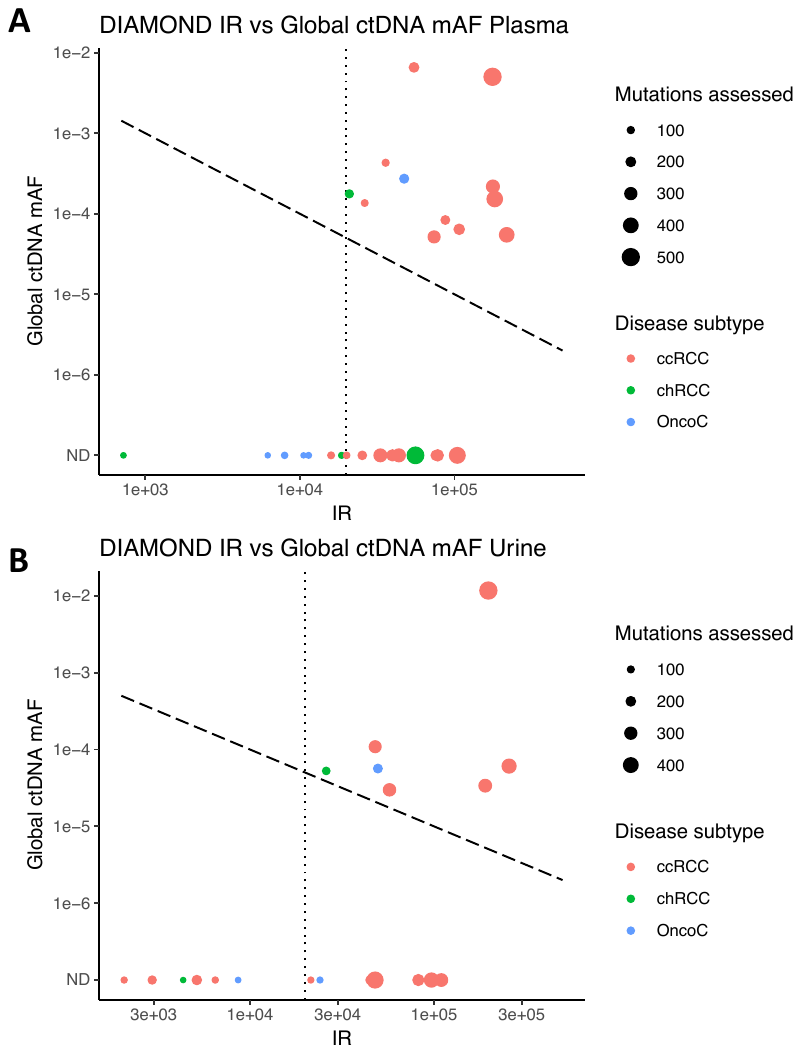

Two-dimensional representation of global ctDNA mAF detected plotted against the total number of informative reads (IR; number of unique molecules across patient specific loci for each sample. ctDNA could be detected if its global mAF was higher than 2/ number of unique molecules (falling above the dashed line, here plotted at 1/number of unique molecules). In some samples, few IR were available. In this study a threshold of 20,000 IR (vertical dotted line) was selected for both plasma **(A)** and urine **(B)**, and samples with undetected ctDNA with fewer than 20,000 IR were excluded as technical failures (7 out of 29 samples for plasma and 6 out of 20 for urine). Samples outside this region had detected ctDNA, or had estimated ctDNA levels below 0.01% (confidence ranges for this value vary for each sample depending on total IR)**.** The size of each point corresponds to the number of mutations assessed for each sample by the INVAR algorithm. The colour of the point corresponds to disease subtype.

**Fig. S10. Correlation of tumour size and ctDNA detection.**

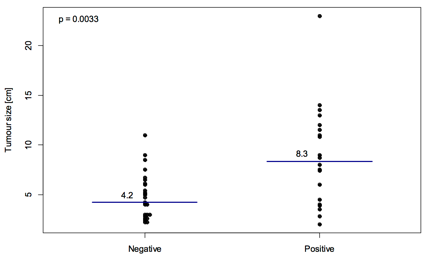

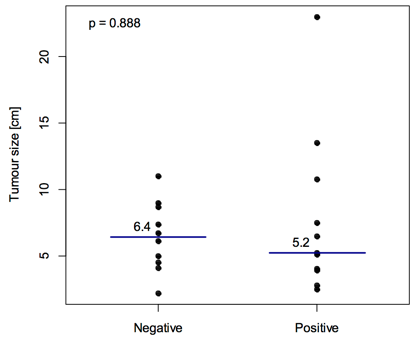
**A	B**

**(A)** Summary of tumour size as measured on CT scanning of patients with any renal tumour, with (right) and without (left) detected plasma ctDNA. Larger tumours were significantly more likely to have detected ctDNA.

**(B)** Equivalent plot but for patients with and without detected ctDNA in urine – there does not appear to be any correlation between tumour size and ctDNA detection.

In both cases, we determined whether there was a relationship through the use of the Mann-Whitney’s U test

**Fig. S11. Correlation of venous tumour thrombus and cell proliferation rates with ctDNA detection.**

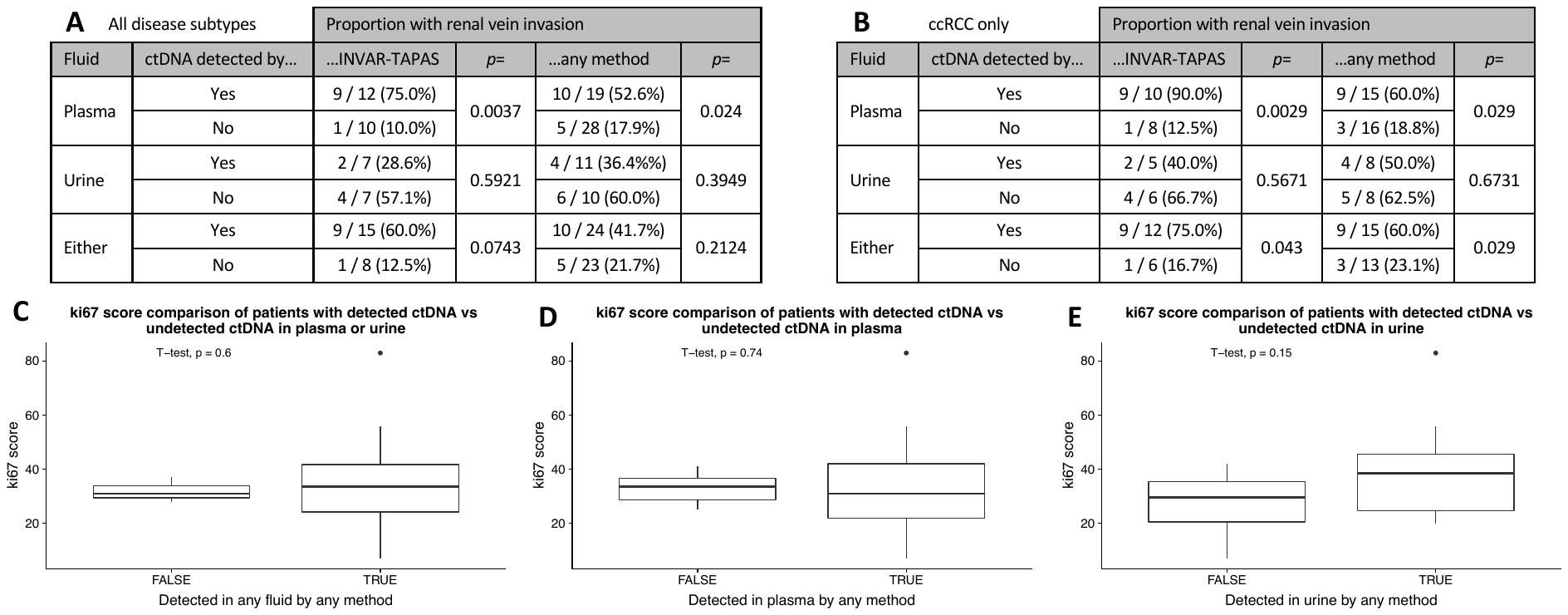

Assessment of the relationship between local renal vein invasion and cellular proliferation rates with ctDNA detection in plasma and/or urine. **(A)** Considering all DIAMOND patients, there was a significant increase in the proportion of patients with renal vein invasion amongst patients with ctDNA detected by INVAR-TAPAS, as compared to those in which ctDNA was not detected. This was not the case in urine, nor when considering both fluids combined. Similarly, the relationship was not evident when considering detection by INVAR-TAPAS and/or tMAD [25 samples were analysed by tMAD but not INVAR-TAPAS]. **(B)** As A but considering only ccRCC patients. The small number of chRCC and oncocytoma patients precluded a similar analysis.

**(C-E)** Comparison of cancer cell proliferation rates, as inferred by ki67 IHC staining of matched tumour tissue, in samples from patients with detected ctDNA in plasma and/or urine **(C)**, in only plasma **(D)** and in only urine **(E)** by any method. There was no evidence of a relationship between ctDNA detection and proliferation rate.

**Fig. S12. Confirmation of pathological classification as oncocytoma**

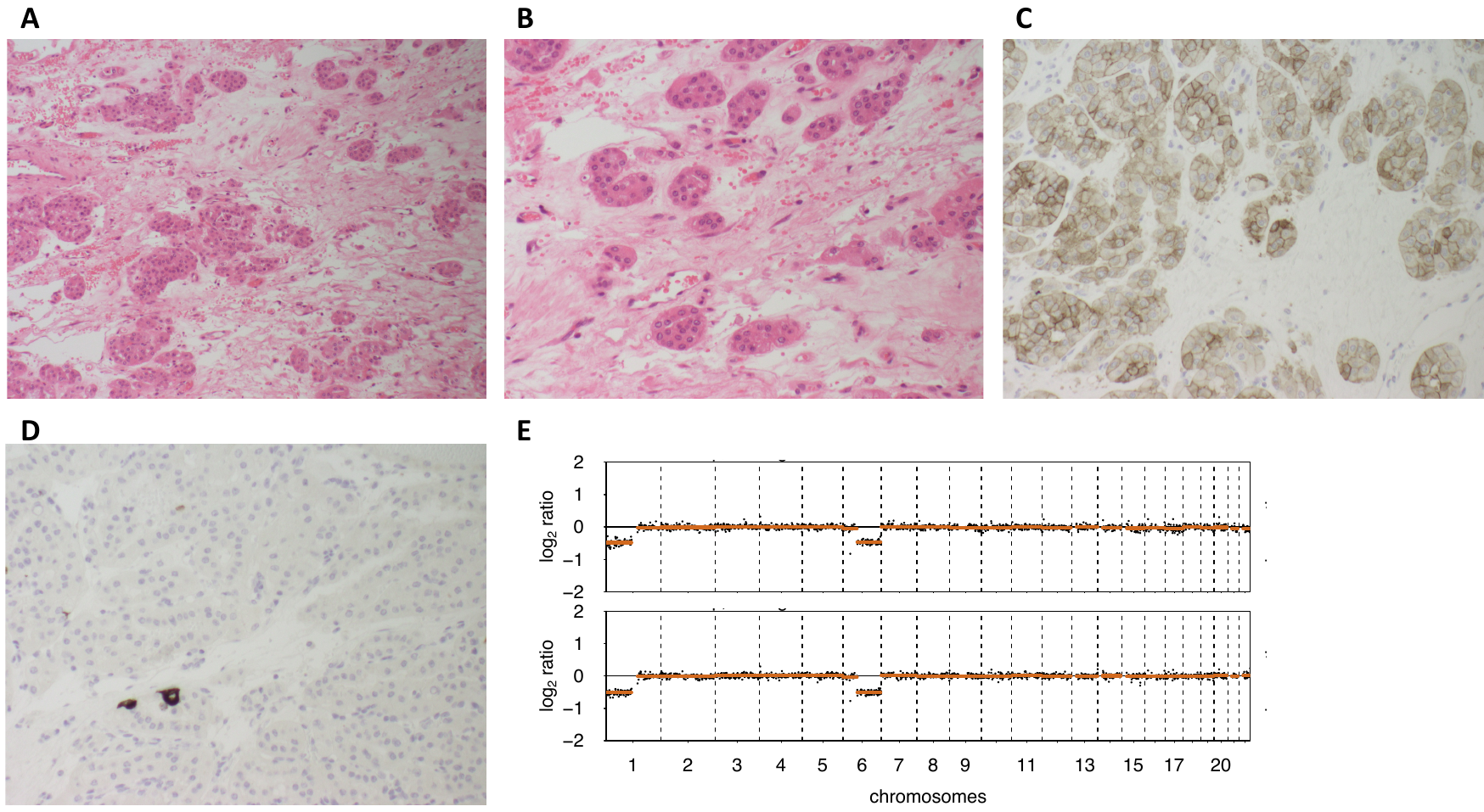

ctDNA was detected in the plasma and urine of a patient with an oncocytoma by INVAR-TAPAS. We assessed pathological and genetic information from matched tissue to confirm this diagnosis.

**(A)** Low magnification H&E stained sections showed the typical architecture of an oncocytoma, with rounded nests of tumour cells within loose hypocellular fibrous stroma. **(B)** High magnification H&E stained section showed the regular round nuclei and densely eosinophilic cytoplasm of tumour cells that are characteristic of an oncocytoma. We also carried out immunohistochemical staining and found that the oncocytic tumour cells were positive for **(C)** CD117, but very few cells were positive for **(D)** CK7. Again, this immunoprofile is consistent with the diagnosis of an oncocytoma. **(E)** sWGS of tumour tissue from this patient revealed a copy number profile consistent with that expected of an oncocytoma (e.g. copy number loss of chromosome 1p)^7^. Shown are profiles from two of four spatially distinct tumour tissue samples. All had identical copy number profiles.

**Fig. S13. Summary of ctDNA detection in all patients and all biofluids.**

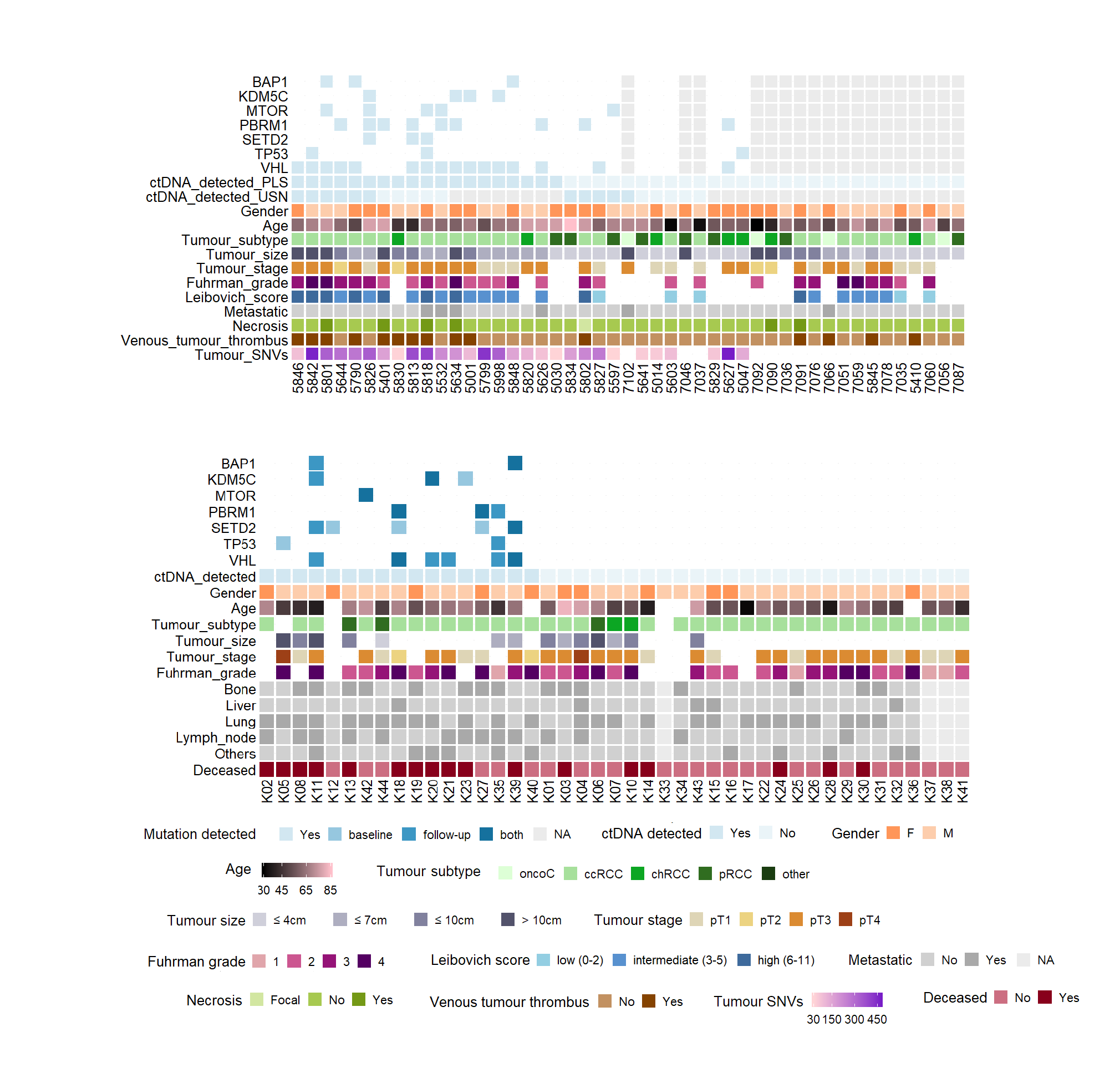

Summary of patient characteristics and ctDNA detection in DIAMOND (top) and MonReC (bottom) cohorts. Both are expanded versions of plots shown in **Fig. 1B** and **1C**. For DIAMOND, ctDNA detection status in plasma (PLS) and urine supernatant (USN) is indicated by rows labelled ‘ctDNA_detected_PLS’ and ‘ctDNA_detected_USN’ respectively. For MonReC, ctDNA detection in plasma is indicated by the row labelled’ctDNA_detected’.

**Fig. S14. Random Forest (RF) model for plasma fragment size distributions.**

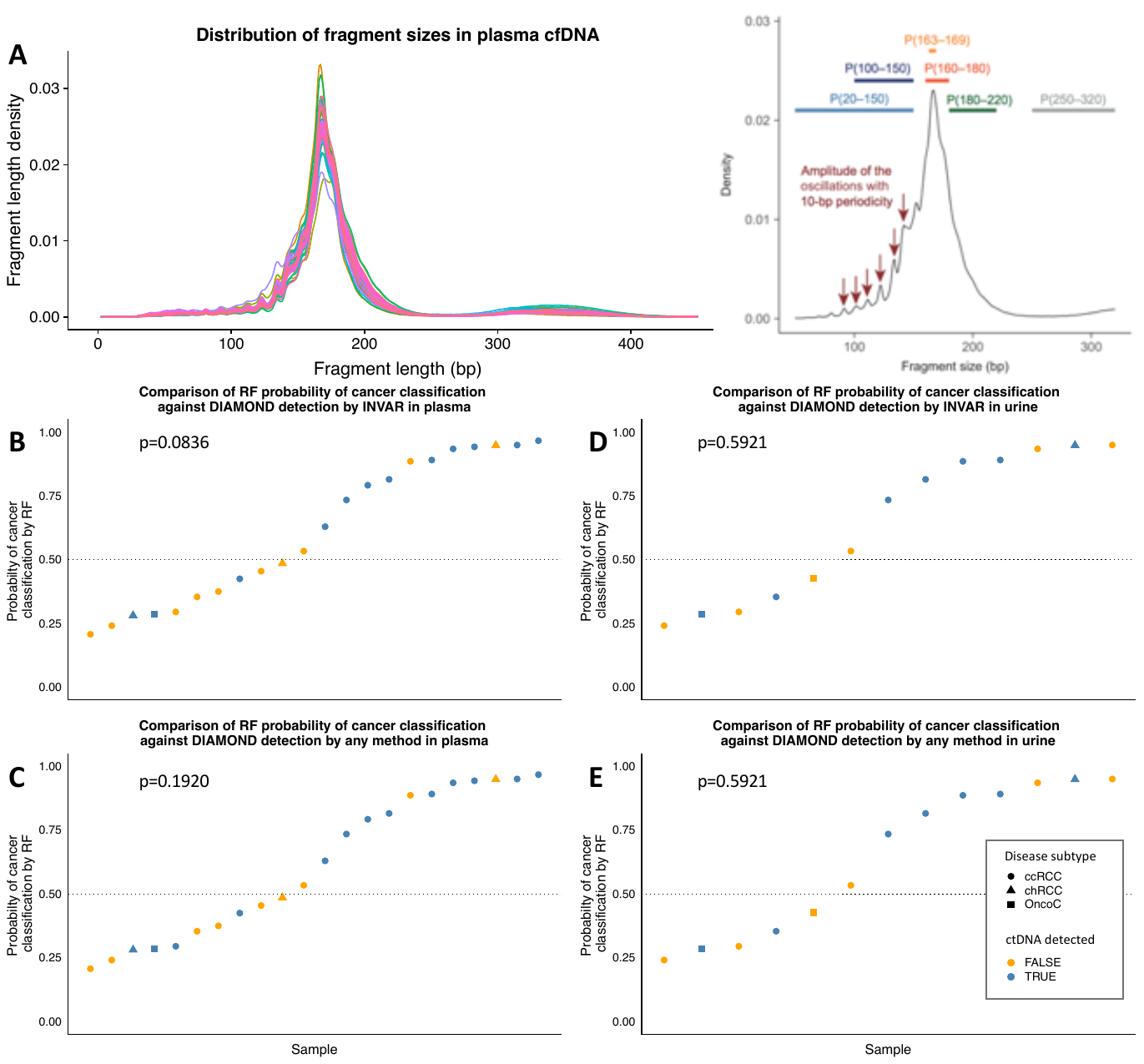

We utilised a Random Forest (RF) cancer classification model that we have demonstrated to be able to differentiate plasma DNA from healthy individuals vs cancer patients based on their fragmentation features (**A,** right)^8^. Here we fed plasma fragment size distributions for each patient (**A,** left) into the RF model and assessed its ability to triage patient plasma samples into those for which higher sensitivity methods are likely to detect ctDNA, and those in which more sensitive methods are unlikely to detect ctDNA. Indeed, the model was able to differentiate patients for whom sensitive methods detected ctDNA in plasma and/or urine, and those in which ctDNA was not detected in either fluid (**Fig. 5**). We further explored the predictive ability of the model by assessing its ability to predict ctDNA detection in **(B)** plasma by INVAR-TAPAS, **(C)** plasma by INVAR-TAPAS or tMAD, **(D)** urine by INVAR-TAPAS and **(E)**, urine by INVAR-TAPAS or tMAD. A blue point indicates patients with detected ctDNA while orange points indicate patients in which ctDNA was not detected. Disease subtype is indicated by data point shape with circle=ccRCC, triangle=chRCC and square=oncocytoma. P values are based on Fisher’s exact test

**Fig. S15.** **Comparison of z-score distribution at baseline, during treatment and when progression occurred.**

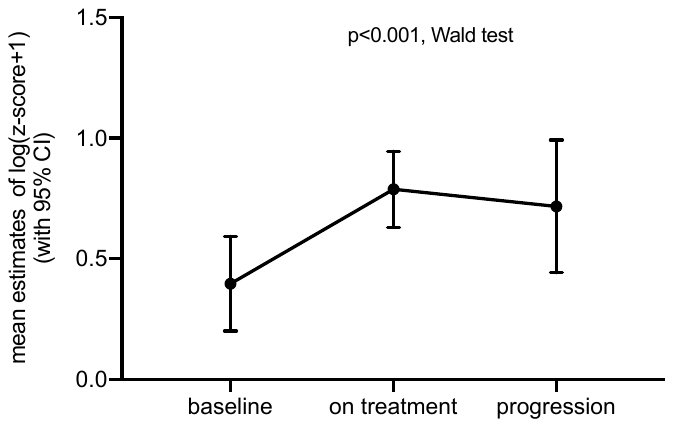

The median time between first and last z-score measurement was 3.6 months (25th-75th percentile: 0.95-7.60). Plotted are overall (joint) difference between mean estimates [baseline 0.40 (95%CI 0.20-0.60), treatment 0.79 (95%CI 0.63-0.95), progression 0.72 (95%CI 0.44-0.99)] of log(zscore+1) (p<0.001). Pairwise difference only significant between baseline as compared to treatment (p<0.001) or progression (p=0.0294), not significant between treatment and progression (p=0.5826).

**Fig. S16. Longitudinal monitoring of mutations detected with the QIAseq custom panel and ichorCNA tumour fractions.**

**A**

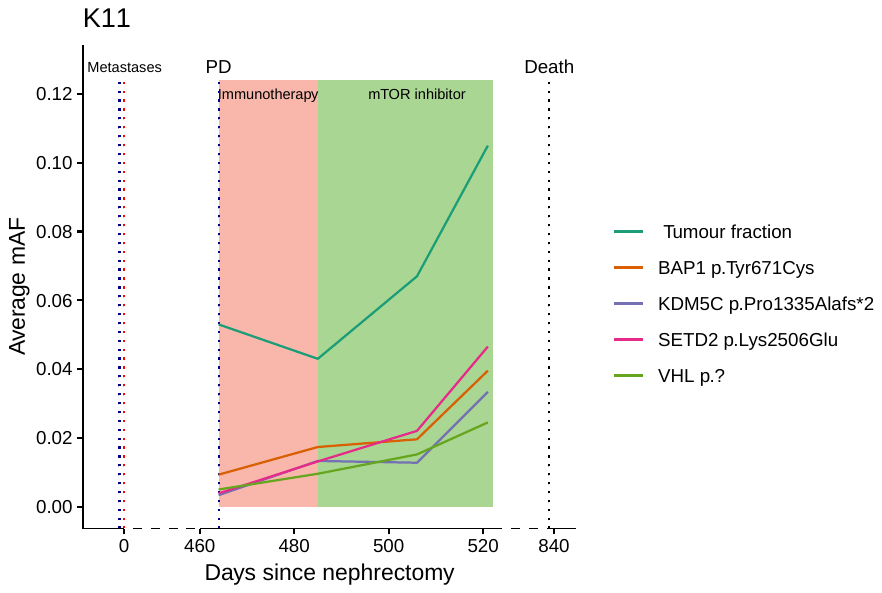

**B**

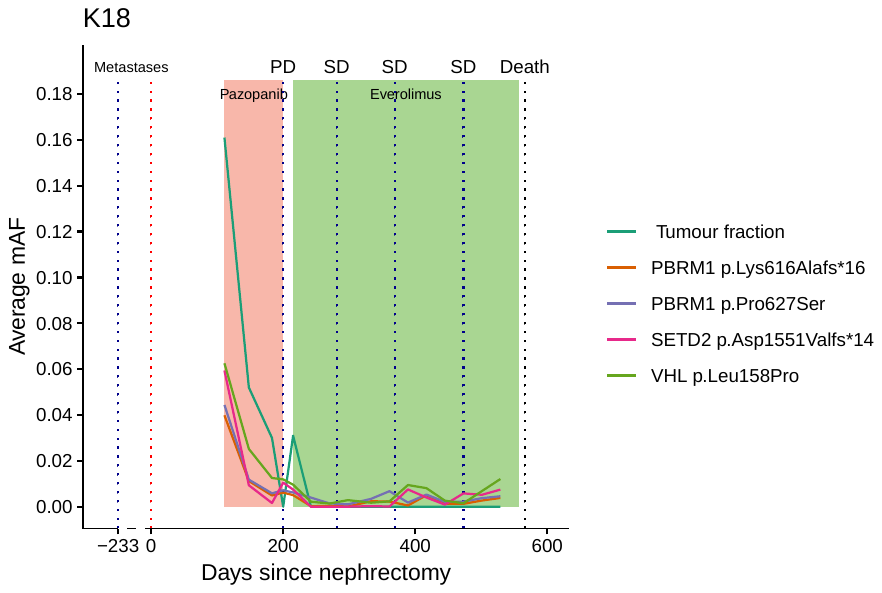

**C**

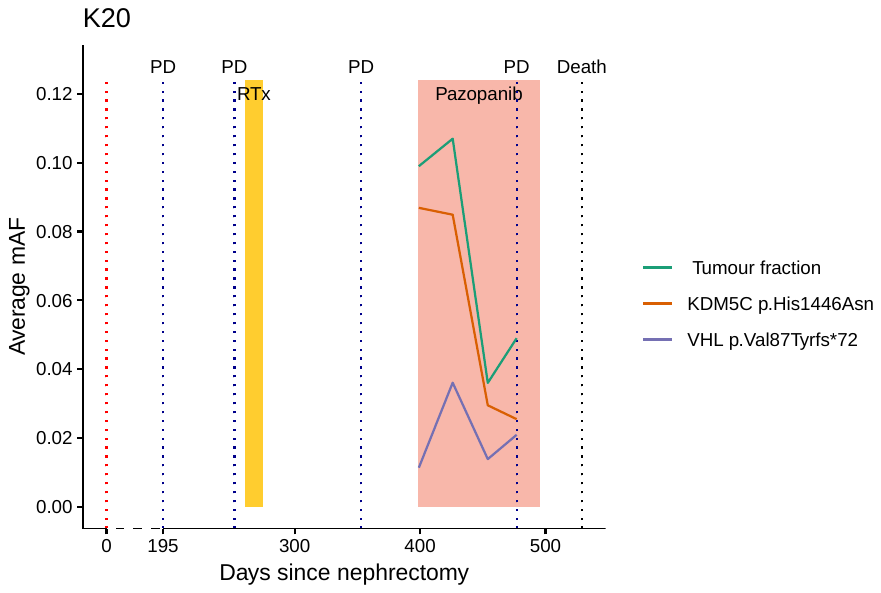

**D**

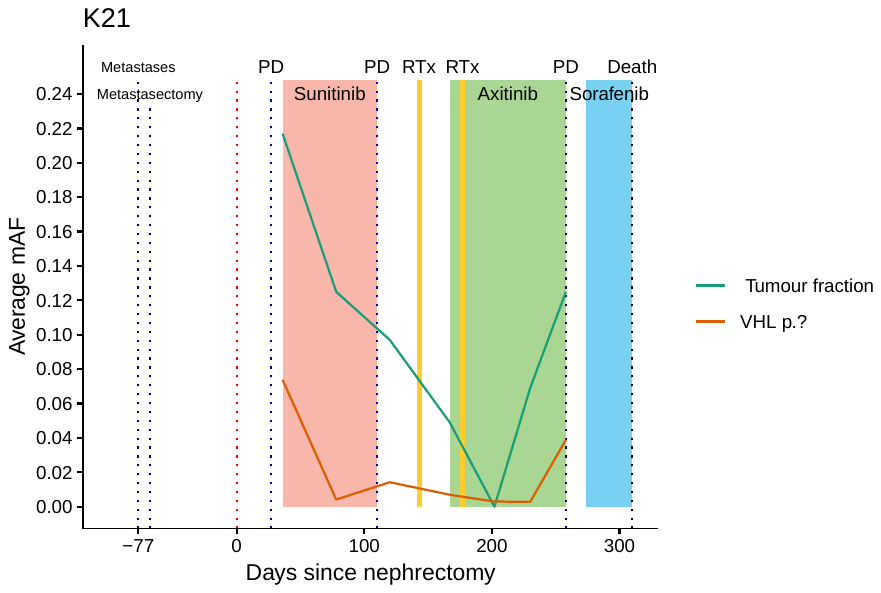

**E**

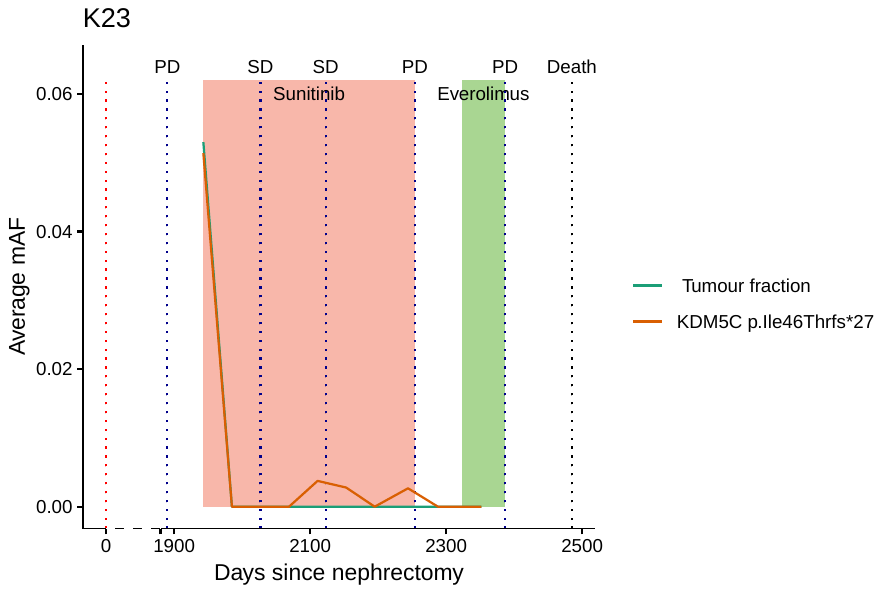

**F**

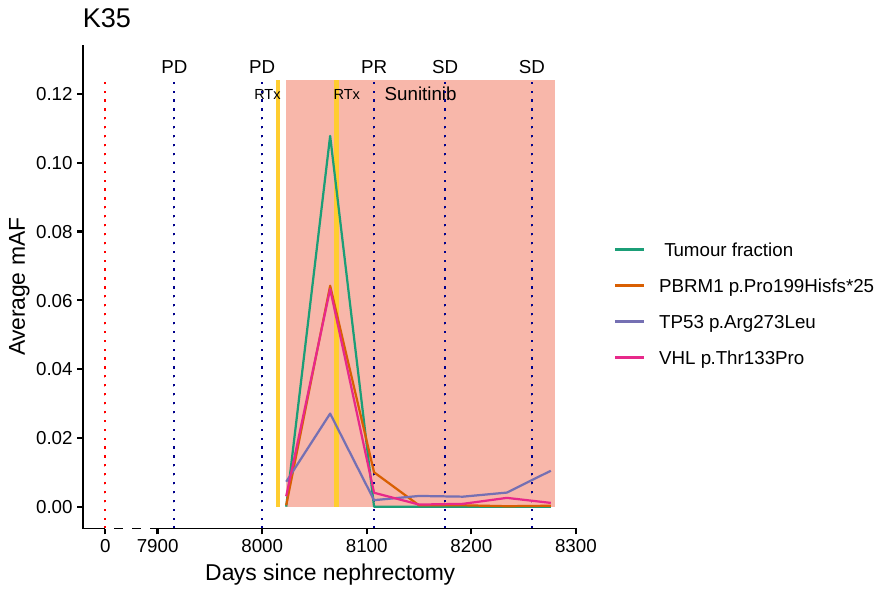

**G**

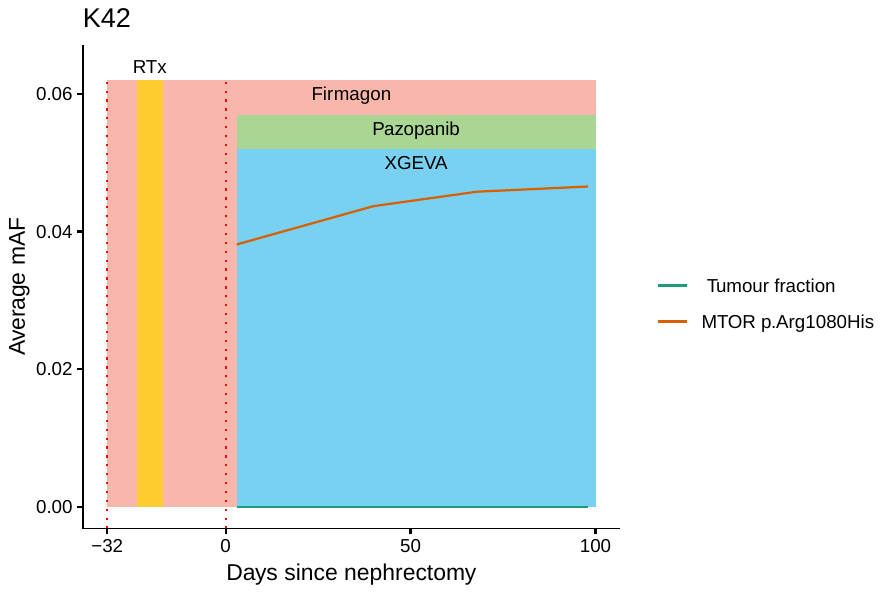

**A-G**, Shown are multiple measurements of ctDNA over time. Therapy regimens are indicated as colours. Disease status at various times (ascertained by computed tomography or MRI), time of diagnosis and if applicable time of death are shown as vertical dotted lines. SD denotes stable disease, PD progressive disease, PR partial response, and MR minor response. ‘Tumour fraction’ was calculated by ichorCNA analysis while individual mutations were observed by QIASeq. Plots for patients K27 and K39 are provided in **Fig. 6B and C** respectively, with detailed notes regarding clinical course provided below;

**Fig. 6B. K27** - The patient was diagnosed with metastases in lymph nodes and bones. At baseline when 1^st^ line treatment with Sunitinib was initiated the patient presented a high tumour load in plasma. Due to progressive disease, treatment was switched to the MTOR inhibitor everolimus. Although a decrease of ctDNA was detected after treatment initiation, the patient progressed again which was accompanied by a rise of ctDNA. After third line initiation, the patient showed a ctDNA response. Time of nephrectomy (red lines), start of radiotherapy (yellow) and time of radiological progression (PD) (black) are shown as vertical dotted lines.

**Fig. 6C. K39** - Nephrectomy and resection of lymph node metastases was carried out fifty-six days before the first blood draw. At baseline when treatment with Sunitinib was initiated the patient presented a high tumor load in plasma, which remained elevated during treatment. After 12 weeks the patient presented clinically progressive disease and died 8 weeks later. Time of nephrectomy (red lines), time of radiological progression (PD) (black)

and time of death (Blue) are shown as vertical dotted lines.

**Fig. S17. Clinical records of select DIAMOND patients.**

**A**

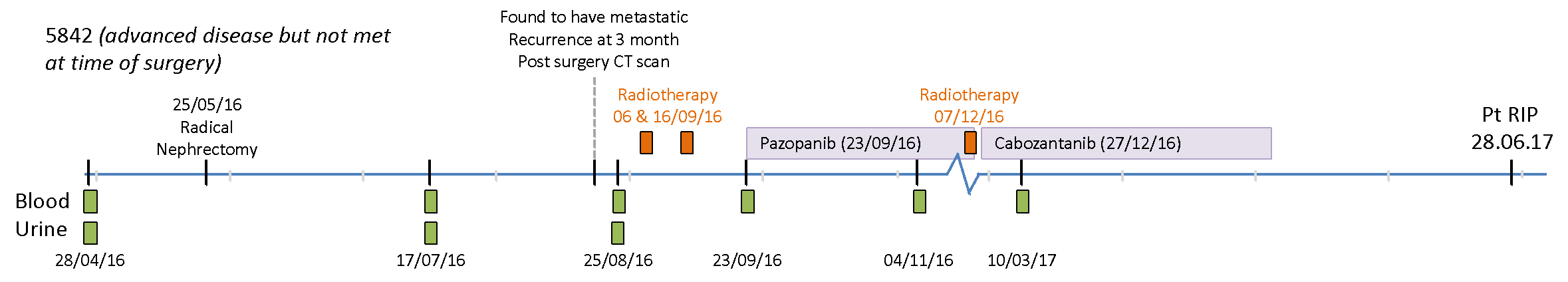

**B**

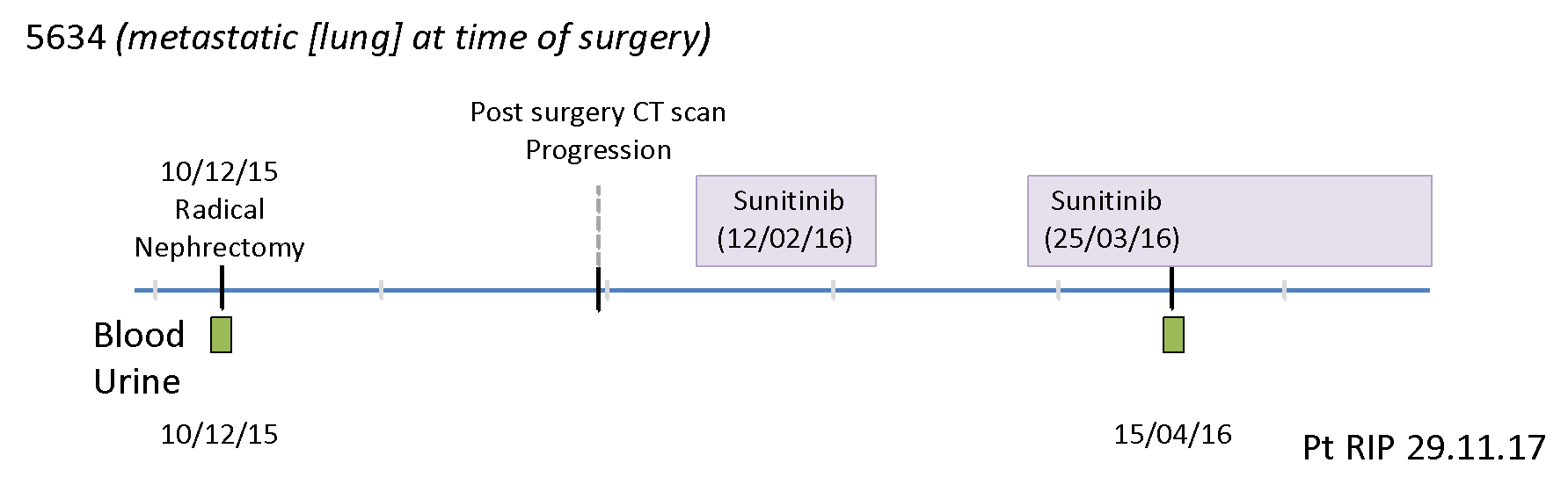

Diagrammatic representation of clinical course of patients **(A)** 5842 and **(B)** 5634.

**Fig. S18. Comparison of imaging data and ctDNA levels (as predicted by INVAR-TAPAS and tMAD).**

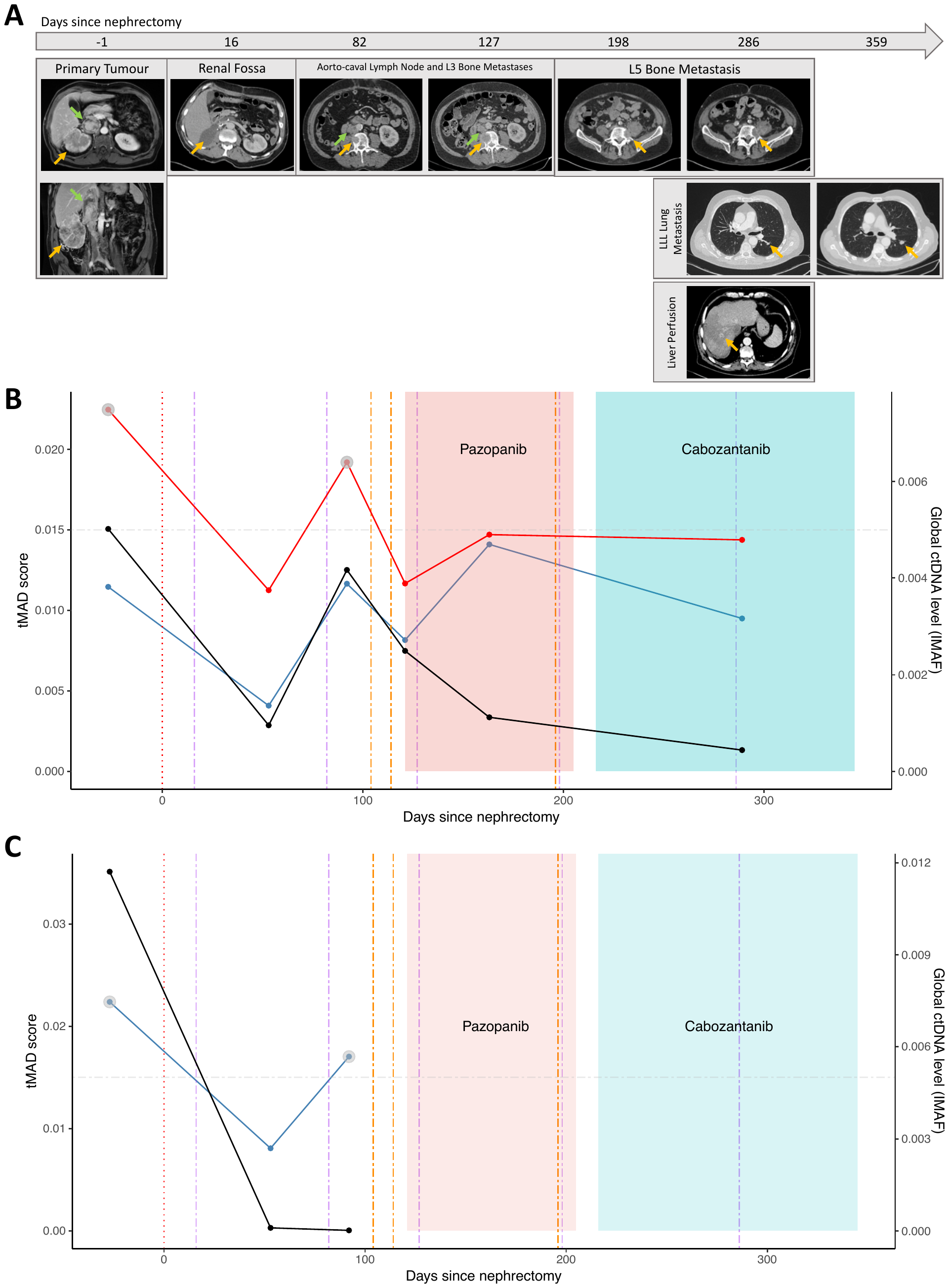

**(A)** CT scan of patient 5842.

Day -1 - Right renal tumour (orange arrow) with tumour thrombus extending into the intrahepatic inferior vena cava (green arrow).

Day 16 - After radical nephrectomy and thrombectomy, fluid collection is visualised in the otherwise normal right renal fossa (orange arrow).

Day 82 - Newly visualised bone metastasis in L3 (orange arrow) and enlarged aorto-caval lymph node (green arrow) on follow-up imaging with interval increase in size (Day 127)

Day 198 - New bone metastasis in L5 with no interval growth (Day 286) under 600 mg of pazopanib. However, new lung metastasis in the left lower lobe (Day 286) with progression on follow-up imaging (Day 359) despite progression to second line therapy with 80 mg cabozantinib. Visualisation of tumour thrombus in the left hepatic vein with surrounding perfusion abnormality (Day 286).

**(B)** Analysis of longitudinal plasma from patient 5842. We applied tMAD (left y-axis) and INVAR-TAPAS (right y-axis) analysis to plasma samples taken throughout their clinical course. Following nephrectomy, whilst global ctDNA levels, as inferred by INVAR-TAPAS (black line), drop dramatically, it is detected (9.5x10-4) at day 53, indicating residual disease post nephrectomy. Conversely, imaging did not detect any residual disease at day 16. ctDNA levels rise (to 4.2x10-3), likely as a result of disease spread to the bone (bone metastasis in L3, orange arrow) and enlarged aorto-caval lymph node (green arrow). Two rounds of radiotherapy (orange dashed lines) and commencement of pazopanib appear to correlate with decreased disease burden, as indicated by a drop in ctDNA levels (2.5x10-3 and 1.1x10-3). This contrasts with imaging that suggests an increase in lesion size. ctDNA levels, with mAF 4.4x10-4, are low at 289 days post nephrectomy, despite clinical progression of the disease that includes new lung metastasis and tumour thrombus in the left hepatic vein. tMAD values pre- (blue line) and post- (red line) in silco size selection are also shown. Whilst tMAD values show a similar trend to INVAR-TAPAS, ctDNA is only detected in size selected samples at the pre surgery and day 92 time points, as indicated by grey circles.

**(C)** as B but longitudinal analysis of USN from patient 5842. Size selection of tMAD data was not carried out.

**Fig. S19. Comparison of SCNA landscape of matched tumour tissue, normal adjacent tissue, and longitudinal plasma samples from DIAMOND patient 5634.**

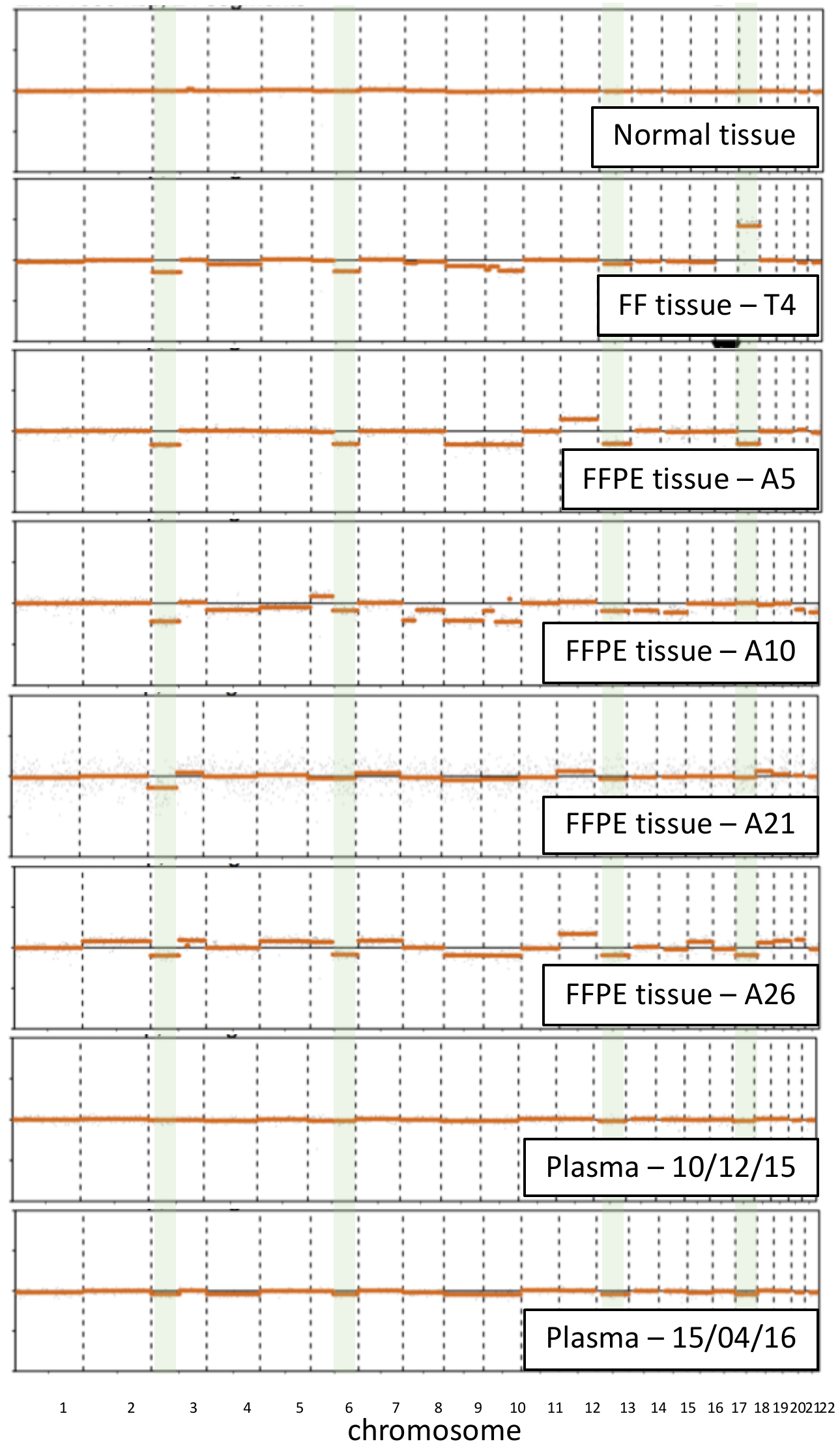

Comparison of the SCNA landscape of matched tumour tissue (FF = fresh frozen, FFPE=formalin fixed paraffin embedded), normal adjacent tissue, and longitudinal plasma samples from ccRCC patient 5634. Chromosome 3p and 5q loss, common events in ccRCC, were observed in tissue all tumour tissue samples. Whilst SCNA were not visible in the baseline plasma sample, low level events can be seen in the follow-up plasma sample that resemble those observed in tissue. The tMAD score reflected this increase in apparent ctDNA levels (**Supplementary Fig. 20**).

**Fig. S20. Comparison of imaging data and ctDNA levels (as predicted by INVAR-TAPAS and tMAD).**

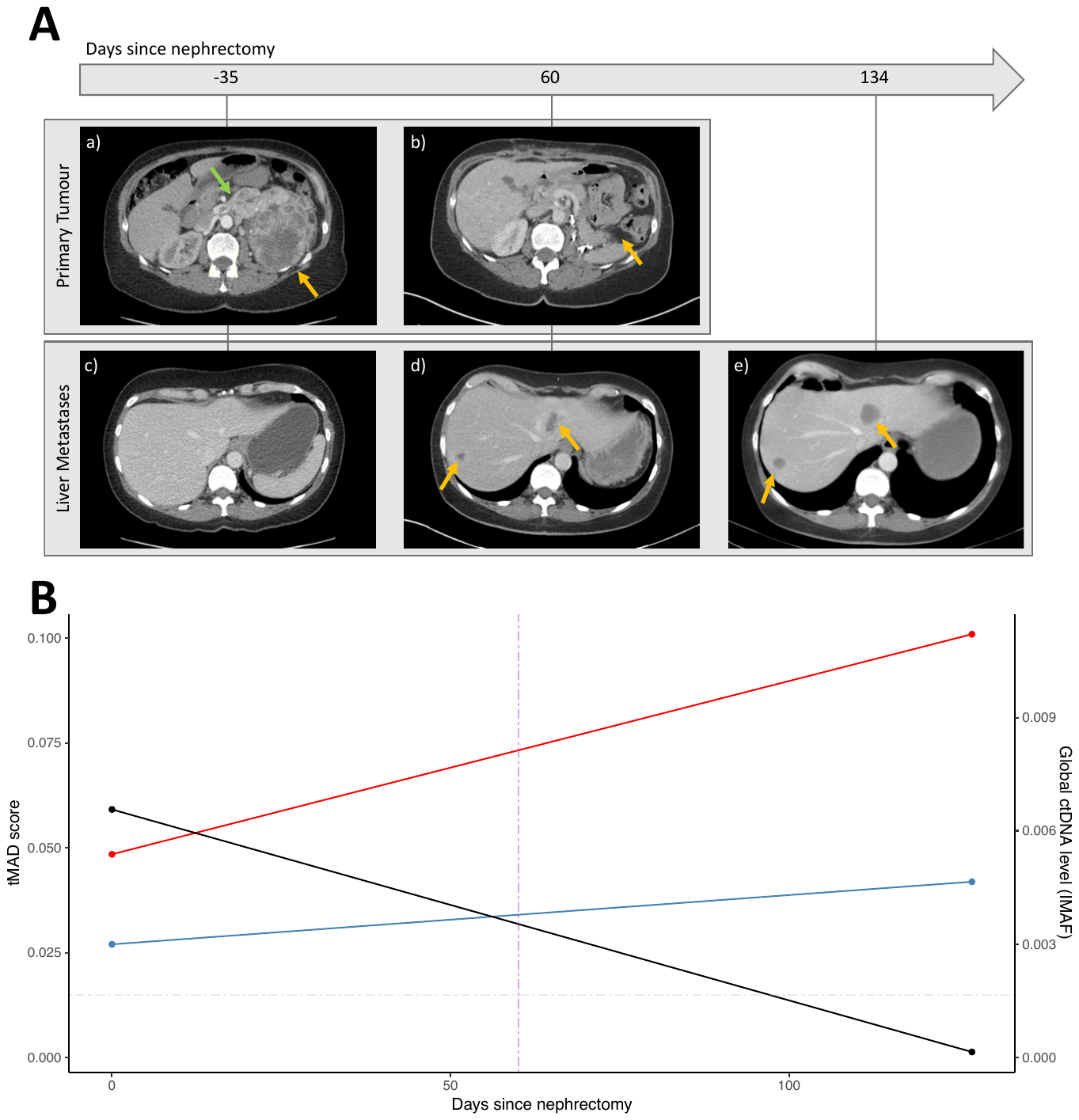

Comparison of imaging data and ctDNA levels (as predicted by INVAR-TAPAS) from patient 5634.

**(A)**

Day -35 - Left renal tumour (orange arrow) with tumour thrombus extending into the left renal vein (green arrow) and patent inferior vena cava (a).

Day 60 - Clear left renal fossa after nephrectomy (b). At baseline, no liver metastases were observed (c) but newly diagnosed liver metastases (orange arrows) at Day 60 follow-up (d) which decreased in size and became increasingly necrotic/cystic at day 134 weeks of sunitinib (e).

**(B)** Analysis of longitudinal plasma from patient 5634. We applied tMAD (left y-axis) and INVAR-TAPAS (right y-axis) analysis to plasma samples taken before and after nephrectomy. Following nephrectomy, global ctDNA levels, as inferred by INVAR-TAPAS (black line), drop from 6.6x10^-3^ to 1.5x10^-4^. tMAD values pre- (blue line) and post- (red line) in silco size selection are also shown. In both cases, and in contrast to the INVAR-TAPAS data, the apparent levels of ctDNA rise.

**Fig. S21. Tumour map of patient 5842.**

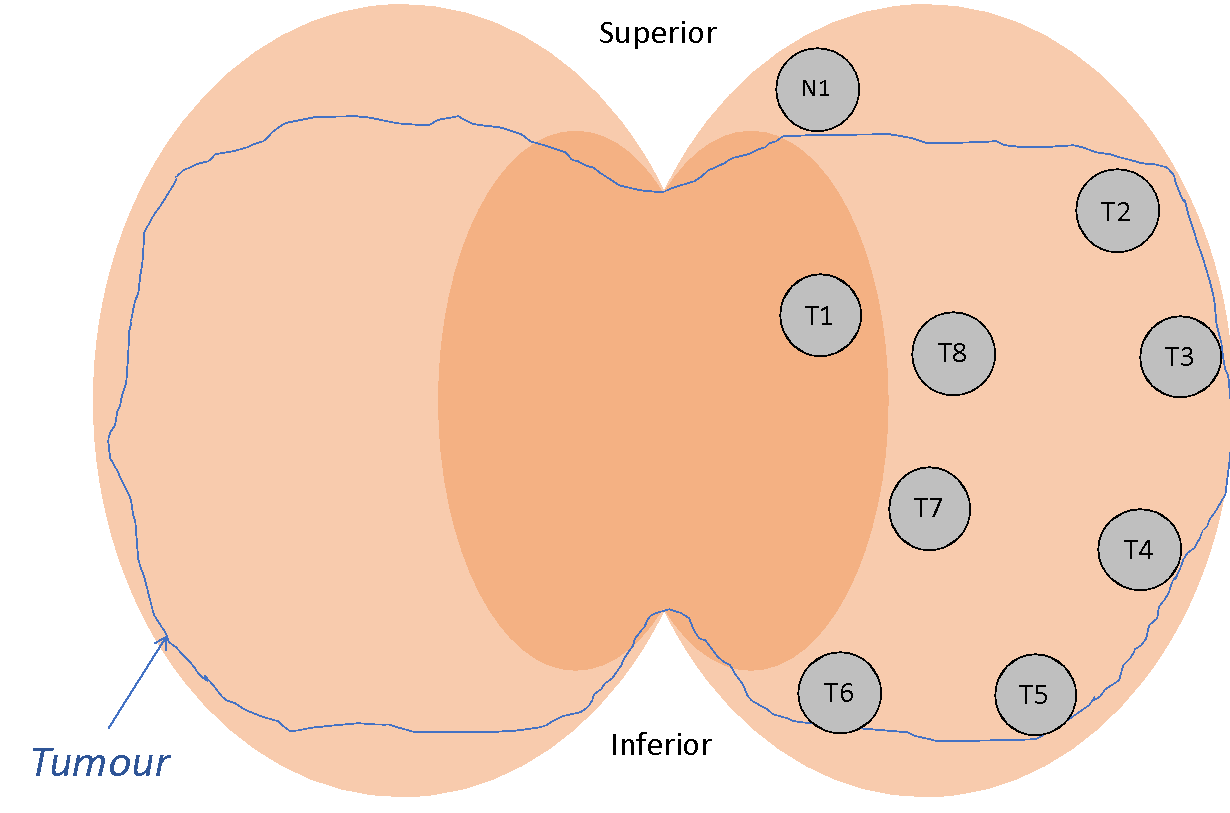

Schematic of a bivalved surgically removed tumour bearing kidney centred on the renal hilum, as is the appearance of the kidney at the time of research sample acquisition by pathologist. Grey circles indicate areas where fresh frozen tissue biopsies were taken. ‘T’ indicates tumour biopsies while ‘N’ indicates matched normal adjacent tissue. Blue line delimits the entire tumour region/area. The locations of T9 (fresh frozen) and A7 (FFPE) biopsies are not known.

**Fig. S22. Comparison between mAF and mutation representation in tumour lesions.**

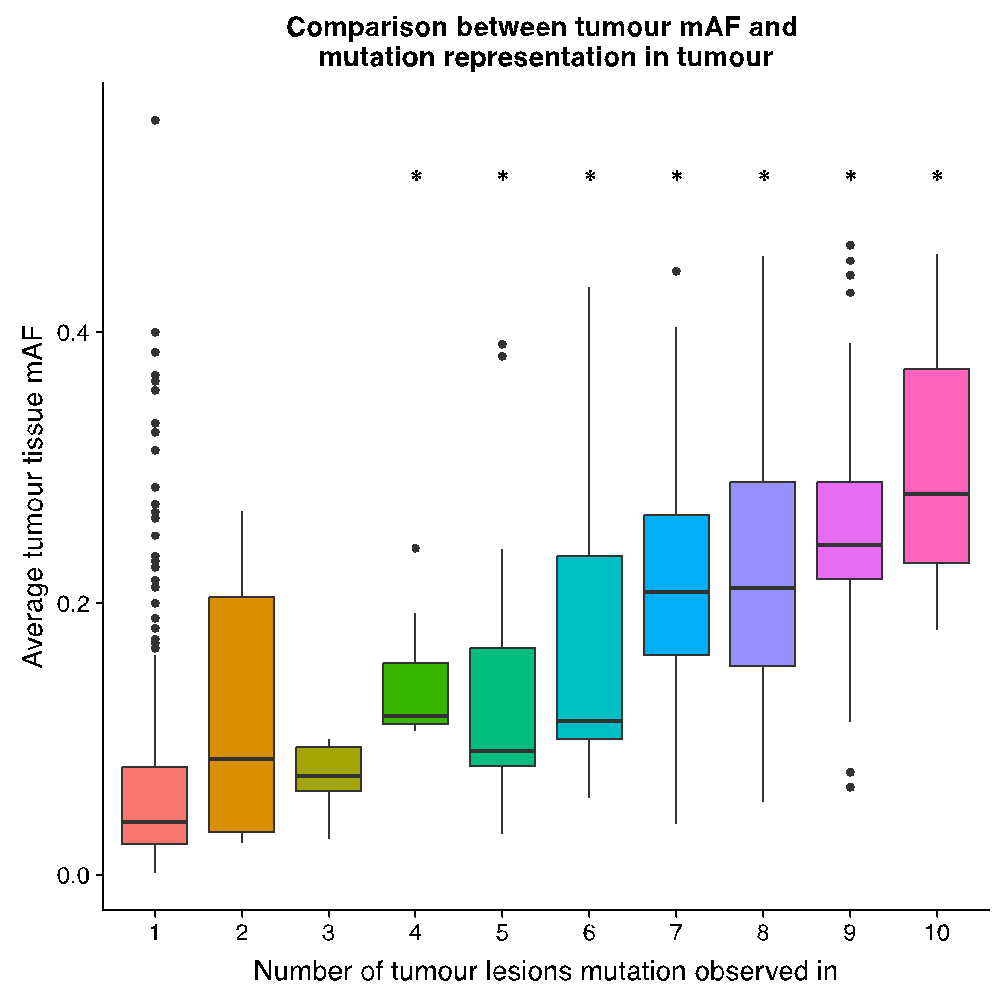

The average observed tumour tissue mAF was found to increase as the number of tumour lesions that mutation was observed in increased. Mutations called in >3 regions had significantly higher mAF as compared to mutations private to one tumour region. This was echoed in plasma and USN ctDNA (**Fig. 6C**).

**Fig. S23. Representation of private mutations in the baseline plasma and USN samples of patient 5842**.

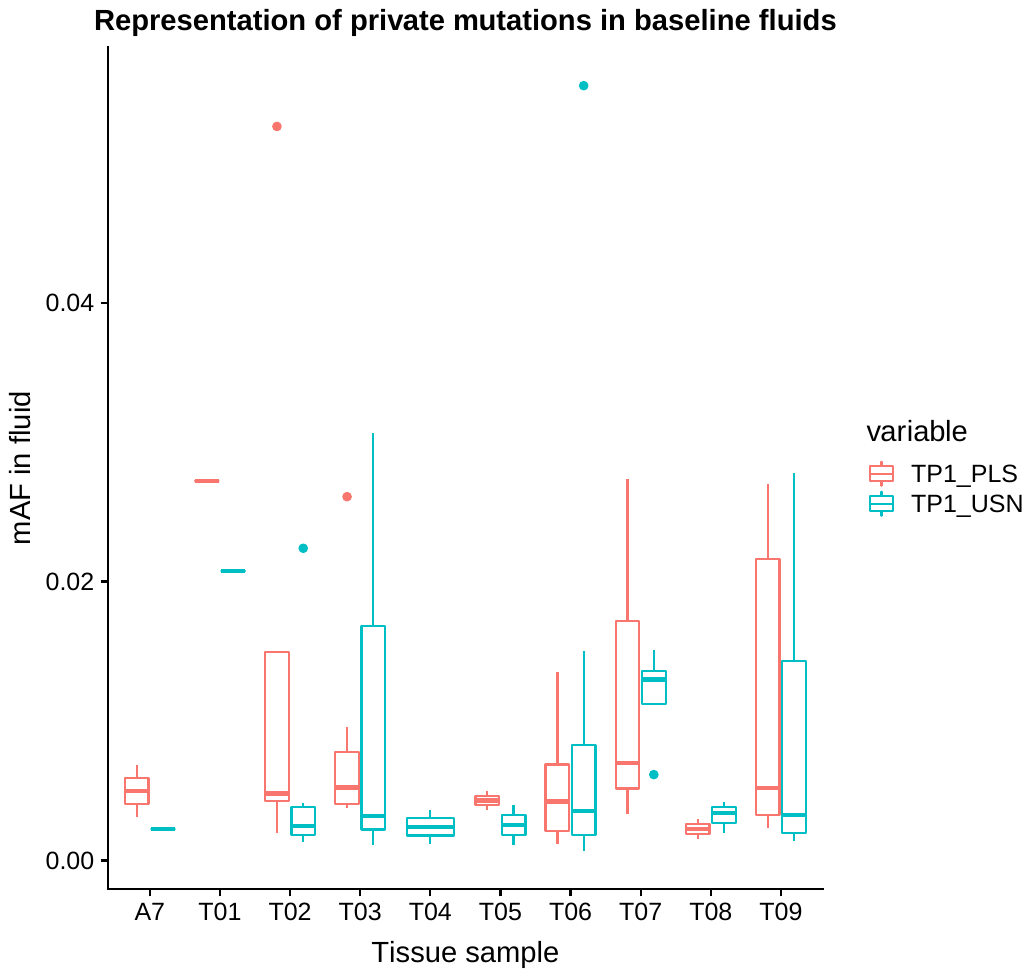

9/10 tumour regions were represented by at least one mutation in plasma (PLS) while 10/10 regions were represented in urine (USN). Amongst the detected mutations, mAF varied to different extents. TP1 = baseline sample from patient 5842.

**Fig. S24. Representation of mutations private to tumour regions in plasma and urine.**

**A**

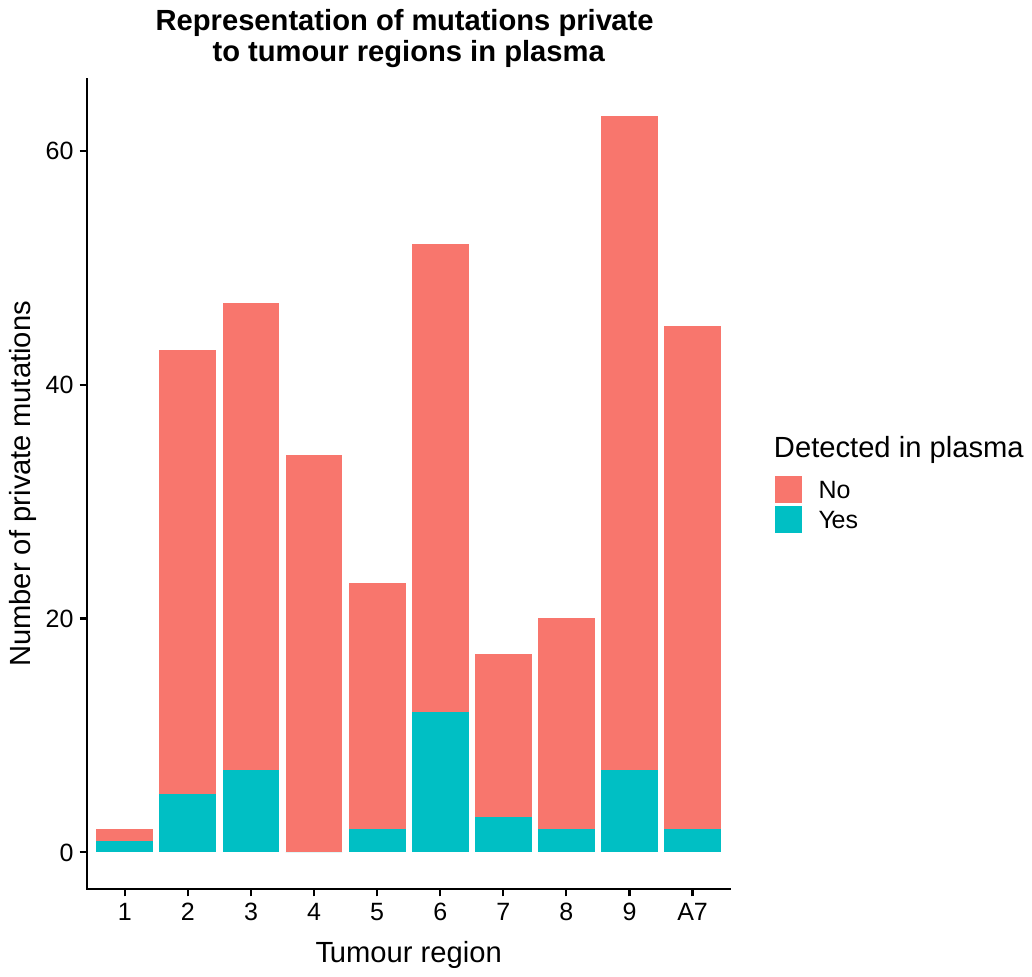

**B**

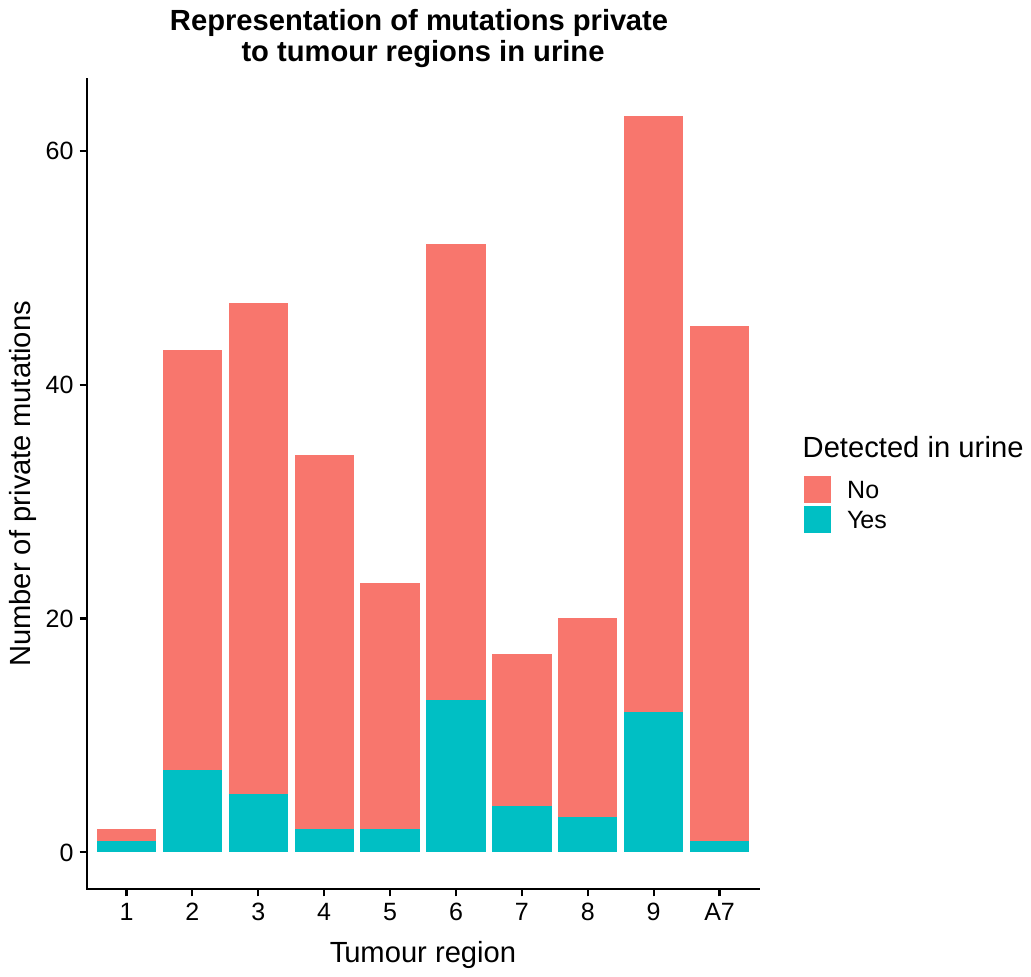

**(A)** Comparison of the number of mutations private to different tumour regions, that were detected in patient plasma. All but one tumour region had at least one private mutation called in plasma. **(B)** As above but for urine.

**Fig. S25. Distribution of mAF of patient specific mutations in plasma and urine from patient 5842.**

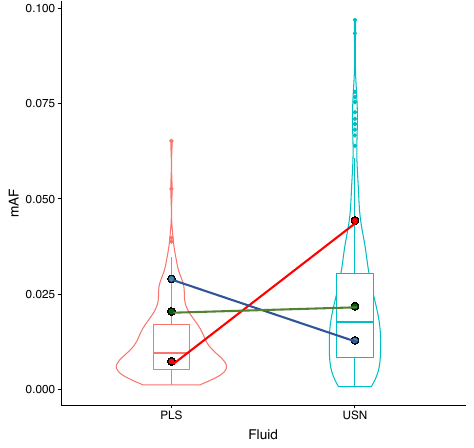

Distribution of mAF of patient specific mutations in pre-surgery plasma (PLS) and urine supernatant (USN). Likely driver mutations (VHL, green; TP53, red; MAPK1 blue; details below) show variable representation in each fluid as compared to other mutations (summarised in violin plots) called in that same sample.

VHL = ENST00000256474.2:c.333_340+1delCTACCGAGG

TP53 = ENST00000269305.4:c.574C>T:p.Gln192*

MAPK1 = ENST00000215832.6:c.493-1G>T

**Fig. S26. Assessment of tumour heterogeneity representation in plasma from patient 5634.**

**A**

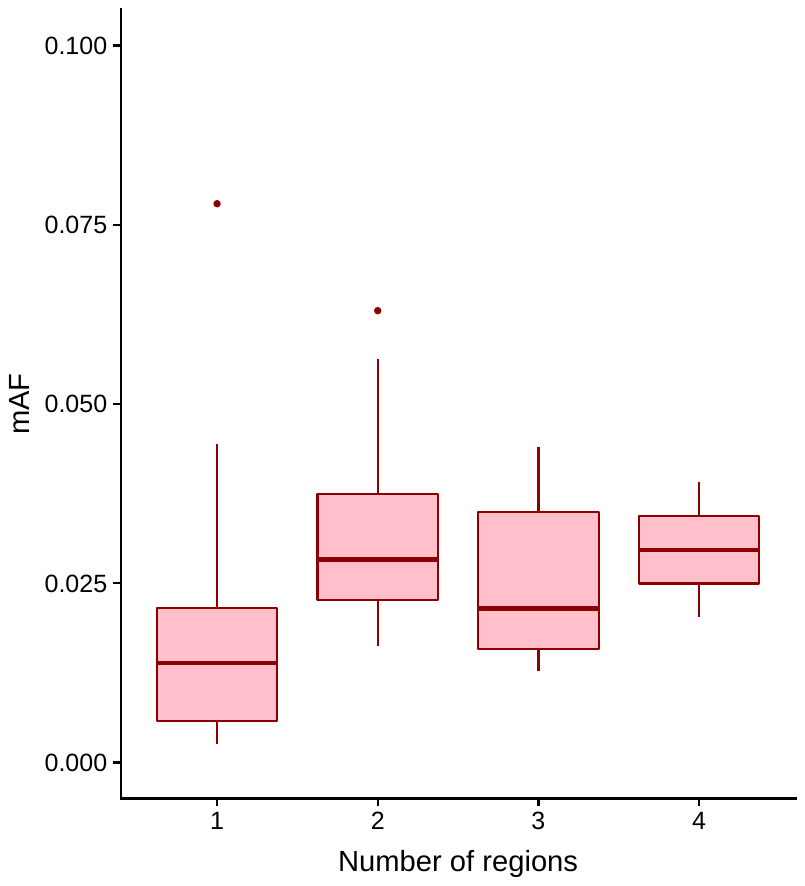

**B**

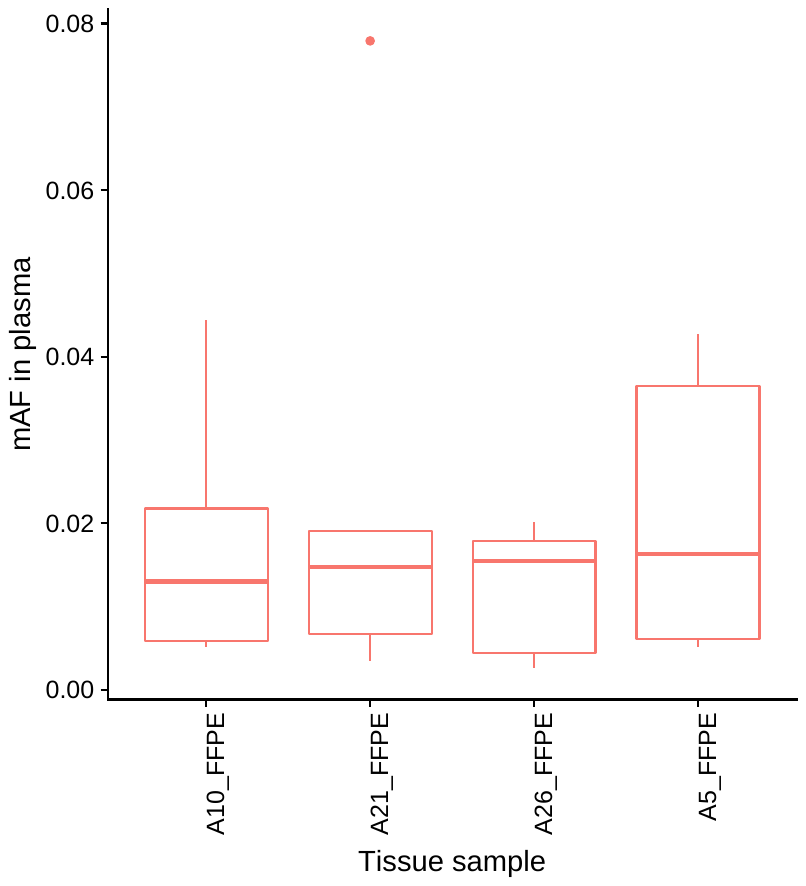

**C**

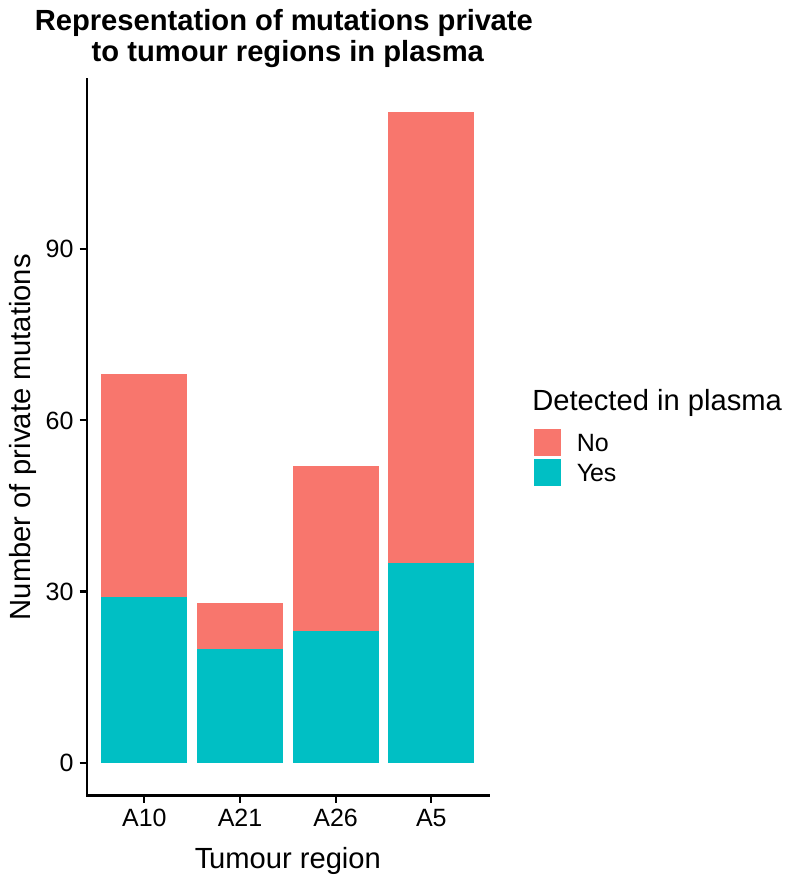

**D**

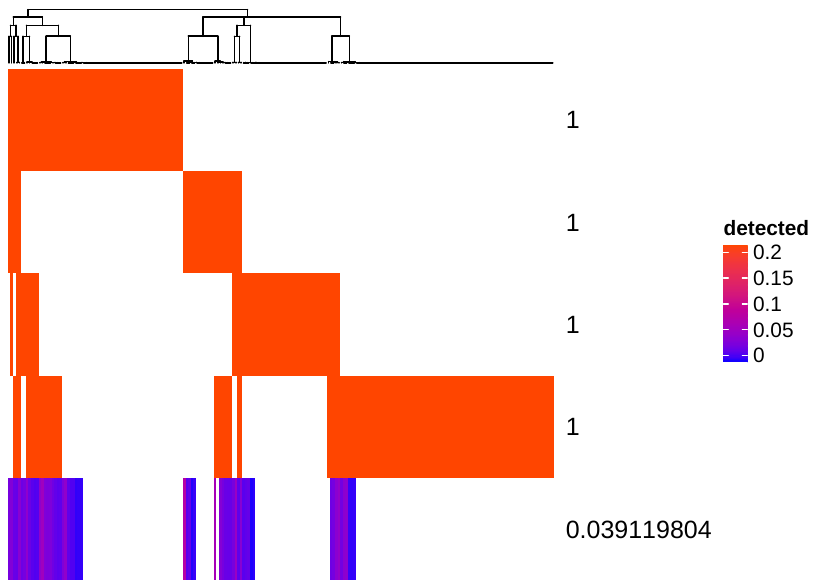

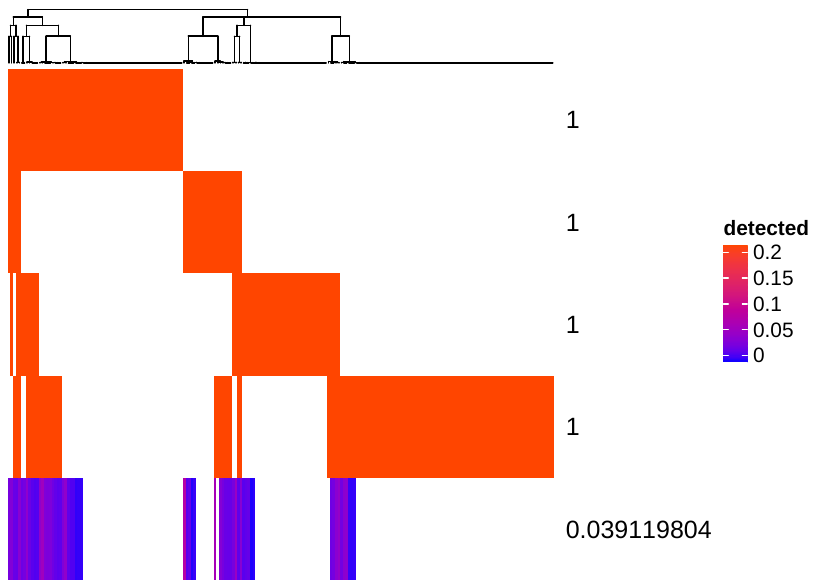

**(A)** Comparison of baseline ctDNA mAF in plasma from patient 5634, and the number of tumour regions that mutation was observed in. **(B)** Representation of private mutations in the plasma of patient 5634. All 4 tumour regions were represented in the plasma. A *DNMT1* mutation (chr19_10305566_G>A; R4C) private to the A21 region of the tumour sits as an outlier at baseline with a mAF of 7.8x10^-2^, the highest mAF of all patient specific mutations identified in this patient. It is unclear why this mutation has such a strong representation in plasma given its limited representation in the tissue (even within sample A21, it’s mAF does not sit as an outlier). It is possible that there is some other tumour clone where this mutation represents a key driver, and it is this clone which is being picked up in the plasma. **(C)** Comparison of the number of mutations private to different tumour regions, that were detected in patient plasma. All tumour regions had >10 private mutations called in plasma. **(D)** Heatmap of mutations detected across 4 FFPE tumour biopsies and baseline plasma, with vertical coloured lines indicating individual SNVs. Hierarchical clustering was by mutation according to Euclidean distance. Colour intensity corresponds to mutation mAF. All mutation clusters, even those private to individual regions, are represented by at least one mutation in plasma.

**Supplementary Materials References**

1.	Network, T. C. G. A. R. *et al.* Comprehensive molecular characterization of clear cell renal cell carcinoma. *Nature* **499,** 43 (2013).

2.	Pal, S. K. *et al.* Evolution of Circulating Tumor DNA Profile from First-line to Subsequent Therapy in Metastatic Renal Cell Carcinoma. *Eur. Urol.* **72,** 557–564 (2017).

3.	Alioto, T. S. *et al.* A comprehensive assessment of somatic mutation detection in cancer using whole-genome sequencing. *Nat. Commun.* **6,** 10001 (2015).

4.	Gerlinger, M. *et al.* Intratumor Heterogeneity and Branched Evolution Revealed by Multiregion Sequencing. *N. Engl. J. Med.* **366,** 883–892 (2012).

5.	LaFlamme, B. Mutational landscape of chromophobe renal cell carcinoma. *Nat. Genet.* **46,** 1050 (2014).

6.	Davis, C. F. *et al.* The somatic genomic landscape of chromophobe renal cell carcinoma. *Cancer Cell* **26,** 319–330 (2014).

7.	Joshi, S. *et al.* The Genomic Landscape of Renal Oncocytoma Identifies a Metabolic Barrier to Tumorigenesis. *Cell Rep.* **13,** 1895–1908 (2015).

8.	Mouliere, F. *et al.* Enhanced detection of circulating tumor DNA by fragment size analysis. *Sci. Transl. Med.* **10,** eaat4921 (2018).
